# Corallimorpharian genome supports the monophyly of Scleractinia and illuminates the cellular evolution of coral calcification

**DOI:** 10.64898/2026.09.25.754439

**Authors:** Thomas D. Lewin, Tosuke Sakagami, Victor M. Piñon-Gonzalez, Yuki Yoshioka, Li-Jung Kao, Yi-Ling Chiu, Kohsei Sasaki, Yi-Hua Chen, Jeng-Yi Li, Kai-Xuan Tin, Mei-Yeh Jade Lu, David J. Miller, Shinya Shikina, Mei-Fang Lin, Yi-Jyun Luo

**Author notes:** T.D.L., T.S., and V.M.P.-G. contributed equally to this work.

## Abstract

The production of calcified skeletons by stony corals is the cornerstone of reef ecosystems and the persistence of marine biodiversity. Corallimorpharians, sometimes described as naked corals, resemble stony corals in many respects but lack hardened skeletons. However, whether they are stony corals that secondarily lost their skeletons has been a subject of much debate. Here, we present a chromosome-level genome of the corallimorpharian *Ricordea yuma* and show that macrosynteny and microsynteny analyses provide strong support for corallimorpharians as the sister group to stony corals. This phylogenetic placement indicates that corallimorpharians were not ancestrally skeletonized and therefore occupy a critical outgroup position for reconstructing the origin of coral calcification. Guided by this phylogenetic context, we sequenced whole-polyp single-cell transcriptomes from *R. yuma* and the complex coral *Galaxea fascicularis* and found that the scleractinian skeleton evolved through the emergence of a derived calicoblast cell state in stony corals. A combination of comparative genomics and regeneration experiments in *G. fascicularis* show that the emergence of the calicoblast cell state was accompanied by the evolution of a biomineralization toolkit assembled from lineage-specific innovations and co-opted ancestral components. Although corallimorpharians retain some ancestral genes later incorporated into coral biomineralization, they lack a detectable calicoblast-like transcriptional state and many core calcification genes. Together, our findings indicate that coral calcification arose through coordinated changes in genome architecture, gene repertoire, and cell identity, highlighting this process as a major evolutionary transition driven by the integration of genomic and cellular innovations.

## Introduction

The deposition of calcium carbonate skeletons by stony corals (order Scleractinia) is fundamental to the formation of coral reefs, the most biodiverse marine ecosystems on Earth (Hoegh-Guldberg et al. 2007). Both major clades of scleractinians, known as robust and complex corals (Chen et al. 1995; Romano and Palumbi 1996; Romano and Cairns 2000; Chen et al. 2002; Kitahara et al. 2010), employ biomineralization to construct extracellular aragonitic skeletons (Drake et al. 2020) that support extraordinary biological diversity and sustain key ecological functions. However, a closely related group, the corallimorpharians (also known as mushroom anemones, coral-like anemones, or naked corals), are morphologically very similar to stony corals yet entirely lack calcified skeletons (den Hartog 1980; Daly et al. 2003; Stanley 2003; Medina et al. 2006) (Fig. 1a). Corallimorpharians can inhabit a wide variety of marine environments but most live in shallow, tropical waters, where they can rapidly colonise large areas through asexual reproduction and outcompete reef-building corals (Kuguru et al. 2004; Lin et al. 2013). Like stony corals, they can harbour symbiotic dinoflagellates (LaJeunesse 2002; LaJeunesse et al. 2004) and bleach in response to environmental stressors (Kuguru et al. 2007). However, in comparison to their stony relatives, the Corallimorpharia is species-poor and understudied, with fewer than 50 extant species described (Fautin 2016).

**Figure 1.**
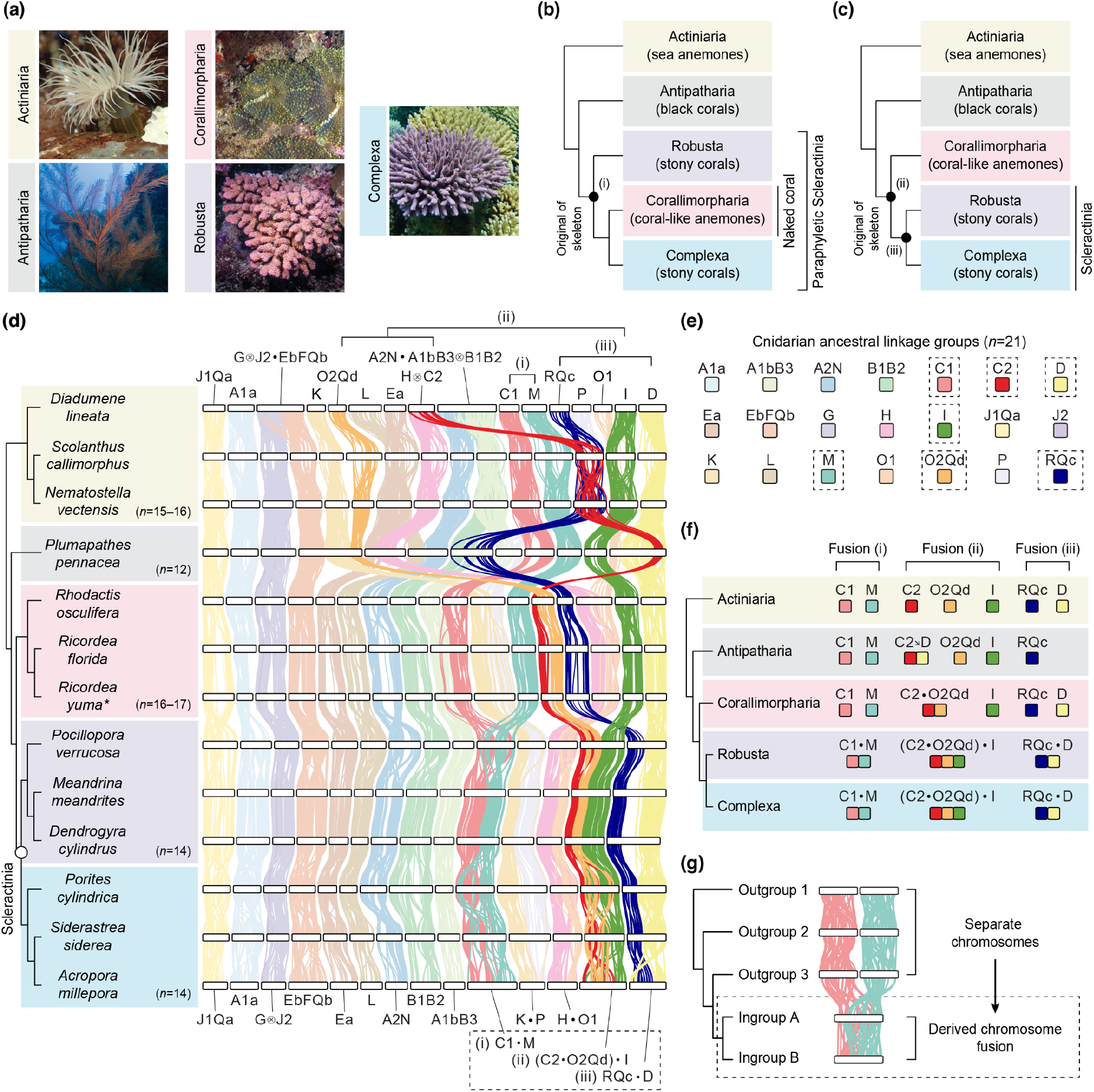
Macrosynteny analysis reveals three derived chromosome fusions unique to scleractinians. (a) Representative photos of species from each hexacorallian clade: *Diadumene lineata* (Actiniaria), *Plumapathes pennacea* (Antipatharia), *Ricordea yuma* (Corallimorpharia), *Pocillopora verrucosa* (Robusta), and *Acropora millepora* (Complexa). Photo credits are given in the Acknowledgments. (b, c) Two hypotheses for the phylogenetic position of corallimorpharians within Hexacorallia. (b) The ‘naked coral’ hypothesis places corallimorpharians within the Scleractinia as sister to complex corals. (c) The monophyletic Scleractinia hypothesis places corallimorpharians as sister to a clade containing both robust and complex corals. Scenarios for the origin of coral skeletons: (i) secondary skeleton loss in Corallimorpharia; (ii) skeleton origin in the common ancestor of Corallimorpharia and Scleractinia, followed by secondary loss in Corallimorpharia; and (iii) skeleton evolution as a scleractinian innovation. (d) Macrosynteny plot of hexacorallian genomes. Each white, horizontal bar represents a chromosome, with vertical lines linking orthologous genes, colored by cnidarian ancestral linkage groups (ALGs). All hexacorallians share the G⊗J2 fusion, and both corallimorpharians and scleractinians share C2●O2Qd, K●P, and H●O1. Three additional fusions (C1●M, (C2●O2Qd)●I, RQc●D) are present in all scleractinians but absent from corallimorpharians, black corals, and sea anemones. The genome sequenced in this study is marked with an asterisk. Phylogenies show the relationships determined in Supplementary Fig. 19. Scleractinian-specific fusions are numbered with Roman numerals. The symbols ●, ⊗ and 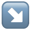 represent chromosome fusion, fusion-with-mixing, and centric insertion, respectively. n, the number of chromosomes. (e) Overview of the 21 cnidarian ALGs, derived from fusion of 29 metazoan ALGs. Several ALGs comprise segments that were separate in the metazoan ancestor but fused and mixed in the cnidarian ancestor (e.g., RQc denotes fusion-with-mixing of metazoan ALGs R and Qc). ALGs involved in coral-specific fusions are highlighted with dashed boxes. (f) Summary of the three derived fusions shared by robust and complex corals but absent from other hexacorallians. (g) Schematic showing phylogenetic information from irreversible fusion events.

Where corallimorpharians sit in the phylogeny of cnidarians has long been debated. They are members of Hexacorallia, together with scleractinians, black corals, and sea anemones, but two competing hypotheses exist for their placement within this group. First, corallimorpharians fall within Scleractinia as sister to Complexa, making true scleractians polyphyletic (the naked coral hypothesis) (Fig. 1b). Second, corallimorpharians are the sister group to all stony corals, including Complexa and Robusta (the monophyletic Scleractinia hypothesis) (Fig. 1c). Early phylogenetic analyses based on mitochondrial protein sequences for the most part supported the naked coral hypothesis (Romano and Cairns 2000; Medina et al. 2006; Kayal et al. 2013; Kitahara et al. 2014; Xiao et al. 2019), but more recent analyses of nuclear genes and mitochondrial nucleotide sequences have largely converged on a monophyletic Scleractinia (Fukami et al. 2008; Kitahara et al. 2014; Lin et al. 2016; Wang et al. 2017; Kayal et al. 2018; Quattrini et al. 2018; Quattrini et al. 2020; McFadden et al. 2021; Wang et al. 2021; Quek et al. 2023; Vaga et al. 2024; Vaga et al. 2025), and gene family and microsynteny analyses of draft genomes also supported scleractian monophyly (Wang et al. 2017).

In recent years, a new approach has emerged to resolving recalcitrant phylogenetic questions. Rare genomic changes can provide sequence-independent tests of phylogenetic hypotheses (Rokas and Holland 2000), and the increasing availability of chromosome-level genomes enables the use of interchromosomal rearrangements, such as chromosome fusions, fissions, and translocations, as molecular synapomorphies (Steenwyk and King 2024). This has proved a powerful technique to distinguishing between a small number of clearly defined phylogenetic hypotheses, clarifying relationships among teleost lineages (Parey et al. 2023), flatfishes (Gallego-Garcia et al. 2026), and animal phyla (Schultz et al. 2023; Lewin, Liao, Chen, et al. 2025; Lewin, Sakagami, et al. 2025). It may therefore also be useful for testing the phylogenetic position of corallimorpharians.

The phylogenetic position of corallimorpharians is critical for our interpretation of how calcification evolves over time and in response to climatic variation. This is because, depending on which hypothesis is accurate, corallimorpharians either lost the ability to form a skeleton (naked coral hypothesis) or never had this ability (monophyletic Scleractinia). Indeed, it has been proposed that corallimorpharians evolved from a calcifying ancestor that secondarily lost its skeleton during the acidifying oceans of the Cretaceous (Medina et al. 2006). In line with this, experimental studies have shown that stony corals can survive prolonged exposure to high acidity without their skeletons (Fine and Tchernov 2007), and corals’ calcifying cells repress calcification genes under temperature stress (Bonacolta et al. 2024; Grau-Bové et al. 2026; Lu et al. 2026), raising the possibility that skeleton loss may be an adaptive response to changing oceans. Under this scenario, corallimorpharians may represent an evolutionary analog for how modern corals will respond to ongoing ocean acidification (Medina et al. 2006; Lin et al. 2014) which, driven by rising carbon dioxide levels, is projected to have severe impacts on coral reefs because it inhibits biomineralization (Gattuso 1998; Guinotte et al. 2006; Hoegh-Guldberg et al. 2007; Guinotte and Fabry 2008; Doney et al. 2009; Mollica et al. 2018). Under the alternative monophyletic Scleractinia hypothesis, corallimorpharians diverged before the emergence of the scleractinian skeleton and instead provide an outgroup for reconstructing how coral calcification first evolved (Quattrini et al. 2020). Resolving the evolutionary position of corallimorpharians is therefore central to understanding both how coral calcification evolved and whether corallimorpharians represent genuine evolutionary analogs of reef corals under ocean acidification.

The scleractinian skeleton itself is produced through a specialized biomineralization program involving a derived ectodermal cell type (Levy et al. 2021) and skeletal organic matrix proteins (SOMPs). These skeletons are formed by specialized ectodermal cells known as calicoblasts (Drake et al. 2020; Levy et al. 2021; Wang et al. 2021) and are composed predominantly of aragonite embedded within an organic matrix of proteins, polysaccharides, and lipids (Stolarski et al. 2021). Calicoblasts concentrate calcium and inorganic carbon into transient amorphous calcium carbonate nanoparticles that are transported to the extracellular calcifying medium, where they crystallize into aragonite (Drake et al. 2013). This process is orchestrated by SOMPs, many of which are highly acidic,enriched in aspartic and glutamic acids, and regulate crystal nucleation, orientation, and growth (Drake et al. 2013; Mummadisetti et al. 2021). The coral skeleton proteome further includes carbonic anhydrases, structural proteins, and extracellular domains of transmembrane proteins incorporated into the growing biomineral (Ramos-Silva et al. 2013). Comparative genomic and proteomic studies suggest that this biomineralization toolkit evolved through the co-option of ancestral cnidarian genes together with lineage-specific innovations and extensive domain shuffling (Ramos-Silva et al. 2013; Takeuchi et al. 2016; Wang et al. 2021). However, it remains an open question whether the origin of the scleractinian skeleton involved the coordinated evolution of both a specialized skeleton-forming cell type and a novel biomineralization program.

This study aims to (i) sequence the genome of the corallimorpharian *Ricordea yuma* to chromosome-level; (ii) use chromosome fusions and gene order as sequence-independent methods of resolving the phylogenetic position of corallimorpharians; (iii) use whole-polyp single-cell transcriptomics on *R. yuma* and multiple reef-building corals to investigate the genetic and cellular basis of skeleton formation; (iv) perform comparative genomic analyses of skeletal organic matrix proteins to determine the origin of the genes integrated into this network; and (v) estimate the timing of corallimorpharian–scleractinian divergence and integrate these estimates with paleoenvironmental reconstructions to define the temporal context in which calcification evolved.

## Results

### A chromosome-level genome of the corallimorpharian *Ricordea yuma*

To study the phylogenetic position of corallimorpharians and the origins of calcification, we first collected a specimen of the corallimorpharian *Ricordea yuma* from Kenting, Taiwan; treated it with menthol to induce the loss of its symbiotic Symbiodiniaceae algae (J.-T. Wang et al. 2012); and sequenced its genome using PacBio HiFi long-read sequencing technology (43× coverage). Scaffolding with Phase Genomics Hi-C data produced a chromosome-level assembly with 96.0% of the genome incorporated into 16 chromosomal pseudomolecules (Figure S1). The final genome is 577 Mb in length, with a scaffold N50 of 35.1 Mb, a GC content of 39.5%, and 95.3% completeness as assessed with Benchmarking Universal Single-Copy Orthologs (BUSCO) Metazoa odb10 database (Figure S2). Genome annotation identified 20,409 protein-coding genes, and repetitive sequences accounted for 41.1% of the genome (Table S1).

### Scleractinians share three chromosome fusions absent in corallimorpharians

We then used our *R. yuma* genome to test whether interchromosomal rearrangements could distinguish between the two hypotheses for the position of corallimorpharians within hexacorallian phylogeny (Fig. 1b, c). To this end, we combined the new genome with 12 additional chromosome-level anthozoan genomes, adopting a ‘quality over quantity’ strategy and restricting the dataset to chromosome-scale genomes generated with PacBio HiFi sequencing. The final dataset consists of 13 members of Hexacorallia: three anemones (Actiniaria), one black coral (Antipatharia), three robust corals (Robusta), three complex corals (Complexa), and three corallimorpharians (Corallimorpharia) (Datasets S1, S2).

Across cnidarians, and more broadly across animals, orthologous genes are retained together on the same chromosomes among distantly related lineages over the course of hundreds of millions of years of divergence (Khalturin et al. 2019; Simakov et al. 2022; Schultz et al. 2023; Zimmermann et al. 2023; Levy et al. 2025; Lewin, Liao, and Luo 2025). Indeed, all cnidarians inherited 21 ancestral linkage groups (ALGs), defined as sets of genes that were linked on the same chromosome in their last common ancestor, and these remain readily identifiable in many extant species (Simakov et al. 2022). From our hexacorallian dataset, we retrieved single-copy orthologs and assigned them to these 21 ALGs, which we then used as markers of chromosomal homology to detect interchromosomal rearrangements (Fig. 1d, Figure S3, Table S2). This macrosynteny analysis revealed chromosome fusions occurring in a stepwise manner (Fig. 1d, Figure S4) throughout the evolutionary history of hexacorallians. First, one fusion (G⊗J2, where letters represent the ALGs and ⊗ represents fusion-with-mixing of the ALGs, when chromosomal fusion is followed by shuffling and interspersal of genes from the two parent chromosomes) is shared across all hexacorallians. Second, three fusions (H●O1, K●P and C2●O2Qd, where ● indicates ALG fusion without subsequent mixing) are shared by corallimorpharians and scleractinians, but are absent in black corals and sea anemones. These are synapomorphies of the clade consisting of Corallimorpharia and Scleractinia, confirming that it is monophyletic. Third, all scleractinians, including both robust and complex corals, share three additional fusions (C1●M, RQc●D, and (C2●O2Qd)●I) that are not found in corallimorpharians, black corals, or sea anemones (Fig. 1d–f). The most parsimonious explanation for this is that these three fusions arose once, in the common ancestor of complex and robust corals, after the split from corallimorpharians (Fig. 1g).

The presence of three fusions shared by complex and robust corals but absent in corallimorpharians strongly supports the monophyly of scleractinians. Despite this, two alternative scenarios could produce this result even under the naked coral hypothesis: independent origins of the three fusions in robust and complex corals, or secondary loss in corallimorpharians. However, unique characteristics of the C1●M fusion rule out both of these possibilities. Across both complex and robust corals, this fusion includes the identical translocation of eight ALG C1 genes into the M region (Figure S5). This demonstrates homology of the complex and robust C1●M fusions, ruling out independent origins. Moreover, corallimorpharians retain these genes on the C1 chromosome rather than M chromosome (Table S3). This rules out secondary loss of the fusion in corallimorpharians, since this scenario would have resulted in the presence of these eight translocated genes on the M chromosome. Thus, the three derived fusions can only be explained by a single origin in the ancestor of scleractinians after their divergence from corallimorpharians, providing rare genomic change evidence, independent of sequence-based phylogenetics, for scleractinian monophyly.

### Conservation of gene order supports the monophyly of scleractinians

We next examined whether conservation of microsynteny, the small-scale order of genes within chromosomes, could give further information as to the phylogenetic placement of corallimorpharians. Microsynteny is expected to decay over evolutionary time, predominantly through chromosomal inversions, so more closely related species typically share more collinear gene blocks (Li et al. 2022; Plessy et al. 2024; Lewin, Liao, Chen, et al. 2025; Schafran et al. 2025). The extent to which microsynteny is conserved is therefore a property that can be applied to phylogenetic inference (Zhao et al. 2021; Almeida-Silva et al. 2023). We hypothesized that, if scleractinians form a monophyletic clade, gene order should be more conserved within scleractinians than between scleractinians and corallimorpharians. Conversely, if corallimorpharians are nested within Scleractina as a sister to complex corals, then these two clades should exhibit the highest conservation.

To test this, we used GENESPACE (Lovell et al. 2022) to quantify collinear gene blocks across genomes. This revealed greater gene order conservation within scleractinians (83.7% conservation of gene blocks) than between scleractinians and corallimorpharians (74.4%) (Fig. 2a), consistent with scleractinian monophyly. To explore this further, we searched for clade-specific stretches of conserved microsynteny using syntenet (Almeida-Silva et al. 2023). This revealed 646 conserved microsyntenic blocks present in both robust and complex corals but absent from corallimorpharians (Fig. 2b). By comparison, corallimorpharians share only 20 and 95 blocks exclusively with robust or complex corals, respectively (Dataset S3). Moreover, a maximum likelihood phylogeny based on a presence–absence matrix of conserved blocks recovers scleractinians as monophyletic (Fig. 2c). Overall, microsynteny analysis reveals higher conservation of gene order within scleractinians than between scleractinians and corallimorpharians, providing another line of evidence for scleractinian monophyly.

**Figure 2.**
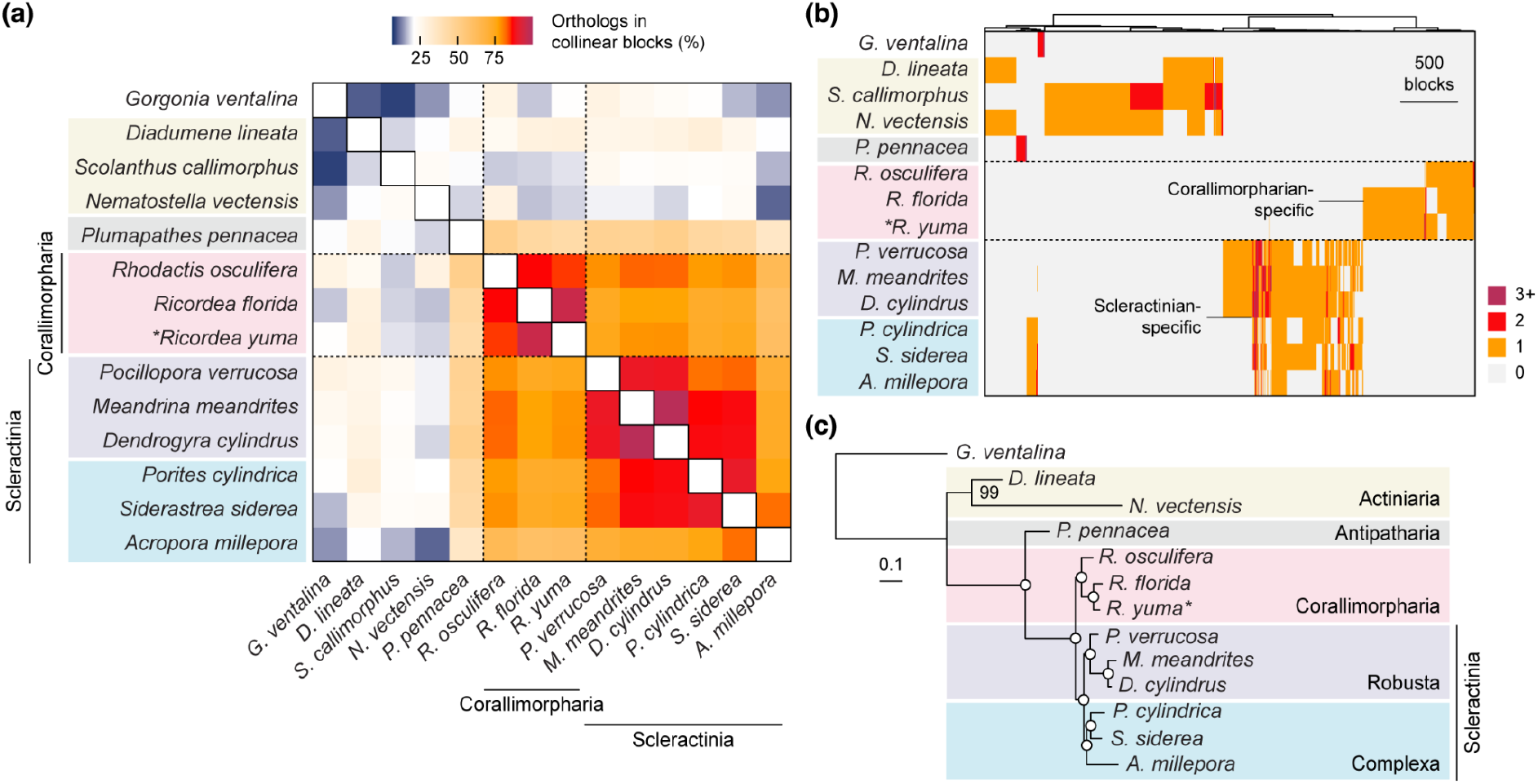
Conserved microsynteny supports the monophyly of scleractinians. (a) Heatmap shows the proportion of orthologs in collinear blocks of at least five genes in pairwise comparisons of species. Robust and complex stony corals have higher microsynteny conservation with each other than either group does with corallimorpharians. (b) Clade-specific microsyntenic gene blocks in Hexacorallia. Each column represents one block (*n* = 4,330). Colors represent the number of copies of a given block in each species. Stony corals share over 900 gene blocks of conserved microsynteny that are absent in corallimorpharians and other hexacorallians, suggesting their monophyly. The corallimorpharians *Rhodactis* and *Ricordea* also share a large number of unique collinear blocks. Columns clustered by Ward’s clustering on a matrix of Euclidean distances. (c) Microsynteny-based phylogenetic inference based on a binarised matrix representing the presence/absence of conserved microsyntenic gene clusters (Almeida-Silva et al. 2023). Maximum likelihood tree was built using the MK+FO+R model with 1,000 bootstrap replicates in IQ-TREE. This topology recovers a monophyletic Scleractinia as sister to Corallimorpharia. Open circles indicate 100% bootstrap support. The genome sequenced in this study is marked with an asterisk.

### Comparative single-cell transcriptomics reveals calicoblasts as a scleractinian innovation

Scleractinian skeleton formation depends on calicoblasts, a specialized epidermal cell type that establishes the physicochemical conditions required for aragonite precipitation (Johnston 1980; Le Tissier 1990; Drake et al. 2020). Although corallimorpharians lack a calcareous skeleton, it remains unclear whether they possess an ancestral or vestigial calicoblast-like cell type. To investigate the evolutionary origin of this skeleton-forming program at the cellular level, we generated whole-polyp single-cell transcriptomic profiles for the corallimorpharian *R. yuma* and the complex coral *G. fascicularis* using a workflow of acetic acid–methanol fixation (García-Castro et al. 2021), fluorescence-activated cell sorting, and 10x Genomics single-cell RNA-sequencing (see Materials and methods) (Figure S6). *G. fascicularis* was not included in the present phylogenetic reconstruction because genomic data for this species were not available at the time of analysis, but previous studies (Ying et al. 2018; Mason et al. 2021) have consistently placed this species within the Complexa.Unsupervised clustering identified 10 transcriptionally distinct populations of cells in each species, including epidermal, gastrodermal, algae-hosting, neural, immune, and cnidocyte cells (Fig. 3a, b). In our initial clustering analysis, a distinct calicoblast cluster could be identified in *G. fascicularis* but was absent in the *R. yuma* dataset, whereas most other major cell-type programs were broadly conserved between the two species (Fig. 3a, b).

**Figure 3.**
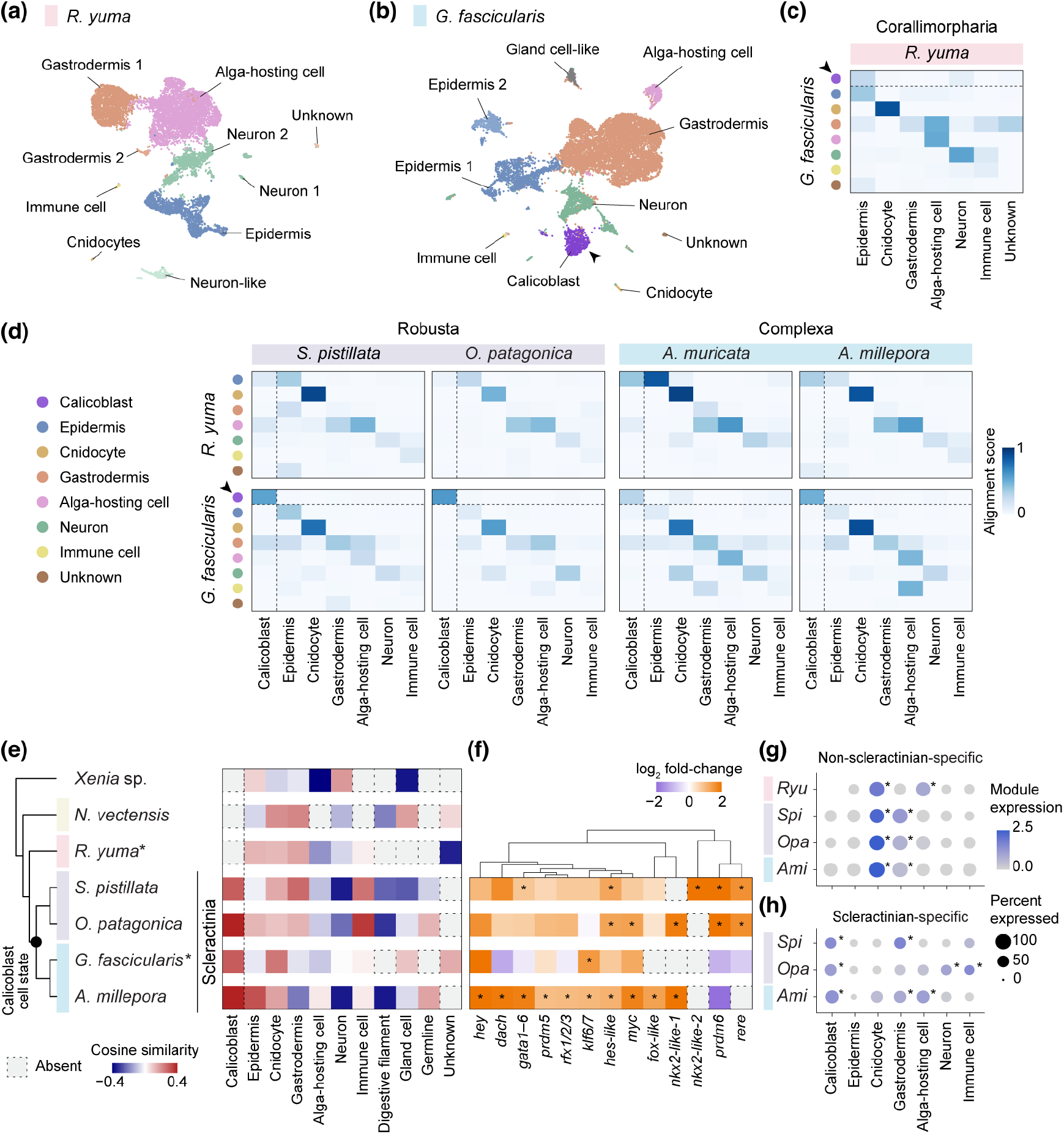
Cross-species single-cell analysis reveals calicoblasts as a novel cell type specific to scleractinian corals. (a) UMAP projection of single-cell transcriptomes from the corallimorpharian *Ricordea yuma* (9,822 cells), identifying 10 major cell types without calicoblasts. (b) Corresponding analysis of the scleractinian coral *Galaxea fascicularis* (12,750 cells), identifying 10 major cell types, including a distinct calicoblast cluster. (c) SAMap alignment between *R. yuma* and *G. fascicularis* cell types. Conserved correspondences were detected across major cell types. Although calicoblasts (marked by dashed lines and arrowheads) showed their closest similarity to epidermal cells in *R. yuma*, no transcriptionally homologous calicoblast-like state was identified. (d) Cross-species SAMap comparisons between *R. yuma* or *G. fascicularis* and four stony corals representing Robusta and Complexa. Calicoblasts showed strong alignment across scleractinian species but lacked a corresponding cell state in *R. yuma*, supporting a scleractinian origin of calicoblasts. (e) Evolutionary origin and cross-species similarity of the calicoblast state. Left, phylogenetic relationships among analyzed anthozoans. Right, heatmap showing the cosine similarity between cell types and a mean embedding vector derived from calicoblast populations within a shared embedding space generated by TranscriptFormer. Scleractinian calicoblasts showed high similarity to this reference state, whereas outgroup species lackeda transcriptionally homologous cell state. Asterisks denote species with single-cell RNA sequencing data generated in this study. (f) Differentially expressed transcription factors in calicoblasts. Heatmap with hierarchical clustering showing representative transcription factors enriched in calicoblasts relative to other cell types, highlighting a distinct regulatory program associated with calicoblast identity. Asterisks indicate significant enrichment (log_2_ fold-change > 1, expression detected in > 10% of cells, and *P* < 0.05). (g, h) Dot plots of average module scores for lineage-specific single-copy orthogroups in the scleractinian + *R. yuma* clade (454 genes; g) and scleractinians (147 genes; h). Asterisks indicate scores significantly higher than other cell types (*t*-test, *P* < 0.01).

To test the robustness of this result, we performed cross-species SAMap alignment (Tarashansky et al. 2021) based on 986 shared single-copy orthologs. This confirmed the conservation of most major cell-type relationships between *R. yuma* and *G. fascicularis*, including cnidocytes, neural cells, and algae-hosting cells (Fig. 3c). However, the *G. fascicularis* calicoblast cluster failed to align with any distinct cell population in *R. yuma*. Instead, calicoblasts showed weak similarity to *R. yuma* epidermal cells, consistent with the developmental origin of calicoblasts as a specialized epidermal derivative (Levy et al. 2021; Bonacolta et al. 2024), but without evidence for a conserved calicoblast-like transcriptional state in the skeleton-free corallimorpharian lineage.

To further investigate the evolutionary origin of the calicoblast cell state, we expanded our comparative analysis to include recently published single-cell atlases from the robust corals *Stylophora pistillata* (Levy et al. 2021) and *Oculina patagonica* (Levy et al. 2025), as well as the complex corals *Acropora millepora* (Levy et al. 2025) and *Acropora muricata* (Han et al. 2025). Across all pairwise comparisons between scleractinians, calicoblasts formed a highly conserved cell-type module shared among both Robusta and Complexa (Fig. 3d). In contrast, no equivalent correspondence was detected between scleractinian calicoblasts and any *R. yuma* cell population. Other major cell types, including epidermis, gastrodermis, cnidocytes, and neurons, remained broadly conserved across both scleractinians and corallimorpharians. To evaluate whether variation in cell number and sequencing depth could account for the apparent absence of calicoblasts in *R. yuma*, we performed a subsampling simulation using the published scRNA-seq datasets. Cells and sequencing depth were randomly downsampled to match those of the *R. yuma* dataset, and the proportion of detectable calicoblast marker genes was quantified (see Materials and methods). At the *R. yuma* cell count (9,822 cells), the proportion of detectable calicoblast marker genes remained nearly complete across all three species: 99.80% in *S. pistillata*, 99.99% in *O. patagonica*, and 99.99% in *A. millepora* (Figure S7), indicating that the absence of calicoblasts in *R. yuma* is unlikely to be explained by insufficient cell number or sequencing depth.

Given that SAMap relies on predefined ortholog relationships, which may limit comparisons across deeply diverged species, we further applied TranscriptFormer (Pearce et al. 2026) to explore the evolutionary origin of calicoblasts within Anthozoa. Unlike orthology-based integration approaches, TranscriptFormer compares genes across species using protein language model embeddings rather than predefined ortholog assignments, allowing comparisons even between distantly related species. Comparative integration across scleractinian corals, the corallimorpharian *R. yuma*, the sea anemone *Nematostella vectensis*, and the soft coral *Xenia* sp. recovered conserved epidermal, neuronal, and gastrodermal programs across Anthozoa (Figure S8). In contrast, no calicoblast-like transcriptional state was detected outside Scleractinia (Fig. 3e). Overall, our single-cell transcriptomic data suggests that calicoblasts represent a scleractinian-specific cellular innovation that is ancestral to both complex and robust corals but evolved after the divergence of corallimorpharians.

### Evolution of a distinct transcriptional identity in calicoblasts

To investigate the molecular features that underpinned the evolution of calicoblasts in scleractinians, we examined the genes enriched in calicoblast cell populations. Compared with epidermal cells, calicoblast-specifc genes were enriched for signaling- and cell adhesion-related functions, including protein kinase activity, receptor-mediated signaling, and cadherin binding (Figure S9). Differential expression analysis also identified a distinct set of transcription factors enriched in calicoblasts, including *gata1–6*, *hes-like*, and *nkx2-like* (Fig. 3f, Figure S10), consistent with previous findings (Levy et al. 2025). Although calicoblasts from robust and complex corals shared a core set of transcription factors, robust corals additionally recruited *rere* and *prdm6* into the calicoblast regulatory program (Fig. 3f, Figure S11). Members of the PRDM family are broadly conserved across Metazoa and have been implicated in developmental gene regulation in diverse animal lineages (Hohenauer and Moore 2012), while *rere* was recently found to be upregulated under heat stress in corals (Grau-Bové et al. 2026). The recruitment of additional transcription factors into calicoblast gene regulatory networks suggests that lineage-specific modifications of ancestral networks contributed to the diversification of this cell type within Scleractinia.

We also sought to test whether calicoblasts exhibit signatures of evolutionary novelty at the transcriptomic level. To this end, we identified scleractinian-specific and scleractinian/corallimorpharian-specific orthologs with OrthoFinder and asked in which cell cluster they are most highly expressed (Figure S12). Scleractinian-specific orthologs showed significantly higher expression in calicoblasts versus other cell types (Fig. 3g, h). In contrast, scleractinian/corallimorpharian-specific orthologs were most highly expressed in cnidocytes and gastrodermis (Fig. 3g, h). This is consistent with phylostratigraphic analyses, which showed an enrichment of hexacorallian and scleractinian genes in calicoblasts (Levy et al. 2021; Lu et al. 2026). Together with their restricted distribution to Scleractinia, this pattern supports the interpretation that calicoblasts represent a stony coral-specific cell-type innovation associated with the evolution of a biomineralized skeleton.

### A scleractinian-specific biomineralization toolkit emerged with the origin of calicoblasts

We next asked whether the genomic machinery underlying calicoblast evolution emerged concurrently with the origin of this skeleton-forming cell type, or instead predated the divergence of Scleractinia and Corallimorpharia. Previous proteomic studies in *Stylophora* and *Acropora* identified skeletal organic matrix proteins (SOMPs) involved in coral calcification, including extracellular matrix proteins, acidic proteins, enzymes, transporters, and lineage-restricted proteins (Moya et al. 2008; Drake et al. 2013; Ramos-Silva et al. 2013; Takeuchi et al. 2016; Mummadisetti et al. 2021). To investigate the origin of these proteins, we performed comparative genomic analyses to characterise the evolution of 38 key SOMPs (Drake et al. 2013; Ramos-Silva et al. 2013; Takeuchi et al. 2016) across the Hexacorallia (Figure S13–17, Dataset S4). This revealed that a substantial fraction of the calcification toolkit is restricted to Scleractinia (Fig. 4a). Among this set of 38 SOMPs, 20 were absent from corallimorpharians and were detected only in stony corals, including galaxins (*GXN* and *GXN2*), the bicarbonate transporter *SLC4γ*, several carbonic anhydrases (*CA*), and skeletal aspartic acid-rich proteins (*SAARs*). Several genes, including *SLC4γ* and *SAAR2*, were conserved across all sampled scleractinians, whereas others showed narrower phylogenetic distributions within robust corals. By contrast, the remaining 18 SOMPs, including mucins, coadhesins, and collagens, were also present in corallimorpharians, of which 11 were broadly distributed across Hexacorallia, suggesting that components of the coral biomineralization toolkit were co-opted from existing genes with other functions.

**Figure 4.**
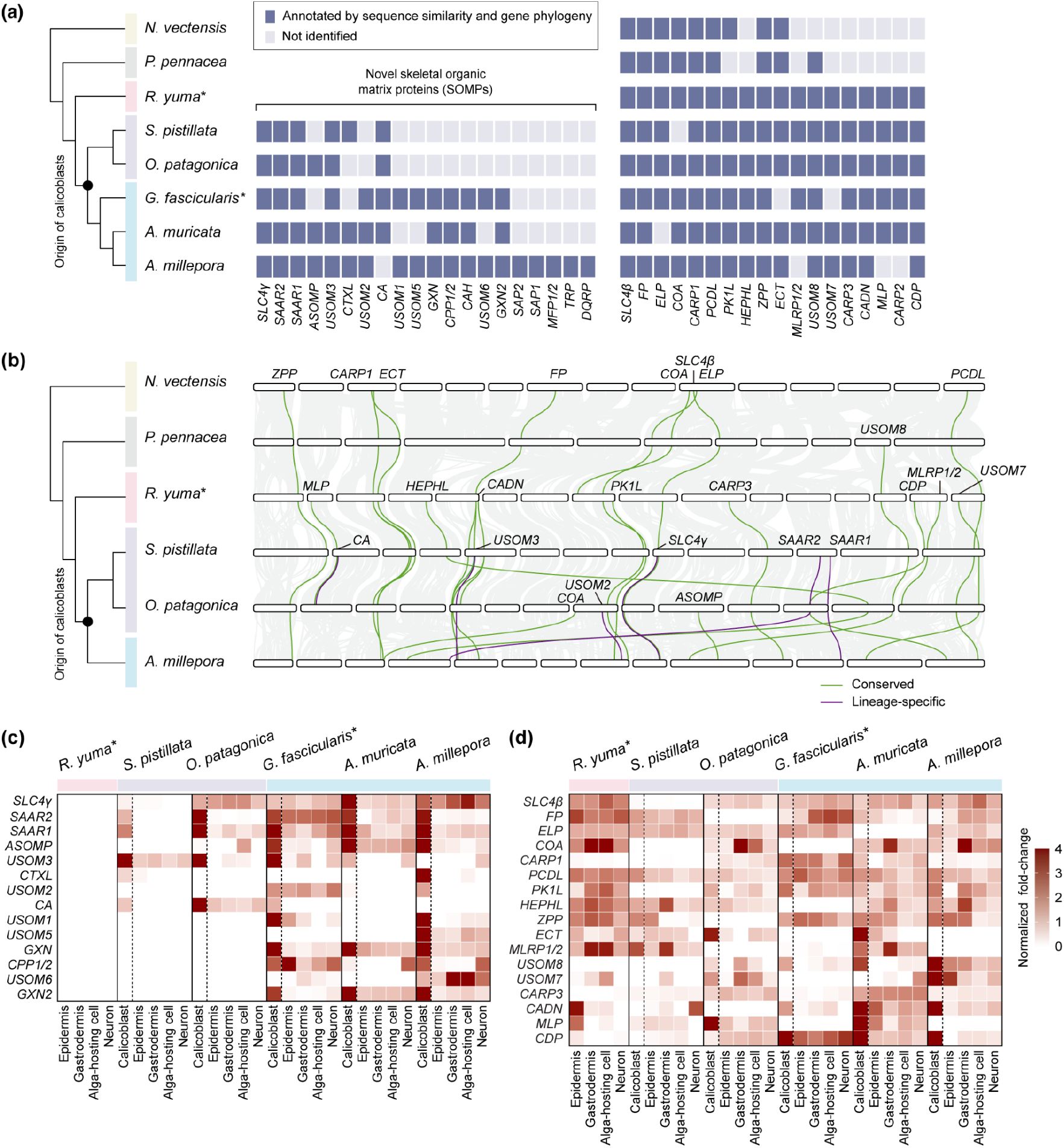
Evolutionary conservation, chromosomal organisation and cell-type specialisation of skeletal organic matrix proteins (SOMPs). (a) Distribution of 38 curated SOMPs across hexacorallians. Genes were annotated using reciprocal BLAST, phylogenetic reconstruction, and synteny analyses. Conserved SOMPs are shown on the left and scleractinian-specific SOMPs on the right, revealing extensive lineage-specific expansion of the coral biomineralization toolkit. The curated SOMPs include acid-rich proteins (*CARPs*, *SAARs*, and *SAPs*), carbonate transport proteins, carbonic anhydrases (*CA*), extracellular matrix proteins, galaxins (*GXNs*), and uncharacterized skeletal organic matrix proteins (*USOMs*). Asterisks denote species with single-cell RNA sequencing data generated in this study. (b) Macrosynteny map showing the chromosomal positions of SOMPs. Vertical lines connect orthologous genes between species. Conserved and scleractinian-specific SOMPs are highlighted in green and purple, respectively, whereas other genes are shown in gray. Gene names are indicated adjacent to the corresponding lines. *CA* and *ECT* are duplicated in some coral species. Most SOMPs are retained on orthologous chromosomes, whereas *HEPHL* and *CDP* in *O. patagonica* and *SAAR2*, *USOM7*, and *USOM8* in *A. millepora* are located on non-orthologous chromosomes. (c) Cross-species single-cell expression heatmap of scleractinian-specific SOMPs across major corallimorpharian and scleractinian cell types. Most lineage-specific SOMPs show strong enrichment in calicoblasts, linking lineage-specific SOMPs to the evolution of the skeleton-forming cell program. (d) Expression heatmap of non-scleractinian-specific SOMPs across coral cell types. Conserved SOMPs show expression across gastrodermal, epidermal, neuronal, and calicoblast populations (dashed lines), consistent with ancestral functions predating coral biomineralization.

Given the importance of 3D genome organization to gene expression (Misteli 2020; Llorens-Giralt et al. 2026), we then investigated the syntenic relationships of SOMPs across hexacorallians. This revealed that certain genes, including *CA* and *ECT*, exhibit lineage-specific tandem duplications in several stony coral lineages (Figure 4b and Figure S13). Moreover, although the majority of SOMPs are localized on orthologous chromosomes across hexacorallians, notable exceptions exist: *HEPHL* and *CDP* in *O. patagonica* as well as *SAAR2*, *USOM7*, and *USOM8* in *A. millepora* are located on non-orthologous chromosomes (Figure 4b). In all other hexacorallian species examined, these genes retain positions on orthologous chromosomes (Figure S18), suggesting that species-specific chromosomal rearrangements have occurred at SOMP loci. Even following species-specific chromosomal rearrangement events, these SOMPs appear to retain their functional roles; for example, *SAAR2* seemingly continues to function as a SOMP in *A. millepora*, as evidenced by its calicoblast-specific expression pattern (Figure 4c).

We next asked how these lineage-specific and ancestral components are integrated to form a coherent calicoblast transcriptional program. Across the six single-cell datasets from corallimorpharians and scleractinians, calicoblasts showed significantly higher SOMP expression than other cell types in all five sampled stony corals (*S. pistillata*: *r* = 0.509; *O. patagonica*: *r* = 0.786; *G. fascicularis*: *r* = 0.665; *A. muricata*: *r* = 0.840; *A. millepora*: *r* = 0.644; *P* < 0.001).

At the gene level, 21 of the 38 analyzed SOMPs showed preferential expression in calicoblasts in at least one coral species (Fig. 4c, d). Fourteen of these 21 genes were scleractinian-specific, including *GXN*, *GXN2*, *SAAR1*, *SAAR2*, and *CA*, directly linking lineage-specific genomic innovations to the calicoblast transcriptional program (Fig. 4c). This pattern was most pronounced in complex corals, which may reflect the fact that many SOMPs were originally identified from *Acropora* skeleton proteomes (Ramos-Silva et al. 2013; Takeuchi et al. 2016).

The expression pattern of non-scleractinian-specific SOMPs contrasted with that of lineage-specific genes. In *R. yuma*, some of these ancestral SOMPs, such as *CADN* (coadhesin), *CDP* (CUB domain protein), and *MLP* (mucin-like protein), were expressed most highly in the epidermis (Fig. 4d). Given the epidermal origin of calicoblasts, this implies the co-option of existing epidermal genes into the calicoblast transcriptional programme. Other non-scleractinian-specific SOMPs, including *COA* (collagen α chain) and *ZPP* (zona pellucida domain-containing protein), were broadly expressed across multiple cell types in *R. yuma*, with the highest enrichment observed in gastrodermal and algae-hosting cells (*r* = 0.725, *P* < 0.001) (Fig. 4d). This pattern is mirrored in stony corals themselves, where these non-scleractinian-specific SOMPs lacked consistent calicoblast-specific expression. It stands in contrast to lineage-specific SOMPs, which showed strong and conserved calicoblast enrichment across scleractinian species (Fig. 4c). Overall, the calicoblast transcriptional program combines lineage-specific biomineralization genes which are strongly restricted to calicoblasts with ancestrally conserved components that remain broadly deployed across multiple cell types.

### Coral calcification recruited genes from diverse ancestral expression contexts

Given the lack of calicoblast-specific expression in many of the non-scleractinian-specific SOMPs, we sought to establish what functions these genes perform outside of calcification and ultimately determine what type of genes were co-opted to function in calcification. To achieve this, we chose to study their expression during tissue regeneration in stony corals. This is because this is a cellular process that predates the evolutionary origin of coral skeleton formation and does not include skeletonization in its early stages, providing an independent biological context in which these genes can be examined outside of the calcification program.

We induced wounds in the coenosarc tissue connecting adjacent polyps in our model stony coral, *G. fascicularis*, by mechanically isolating individual polyps from the colony. We then performed bulk RNA sequencing of the regenerating coenosarc at five time points spanning 0 h to 16 days post-fragmentation (Fig. 5a–e′). During this process, 1,087 genes were differentially expressed, and these could be clustered into five groups, broadly corresponding to baseline tissue maintenance genes (cluster C1) and then early (C3, C4), mid (C2) and late (C5) response genes (Fig. 5f, Dataset S5). Gene ontology (GO) analysis reveals that baseline tissue maintenance genes are enriched for cell cycle and metabolism functions (Fig 5g). In contrast, early response genes are enriched for cell structure remodelling; mid response genes for genome maintenance and signal regulation; and late-acting ones for cell differentiation, organ morphogenesis and calcification (Fig 5g, Dataset S6).

**Figure 5.**
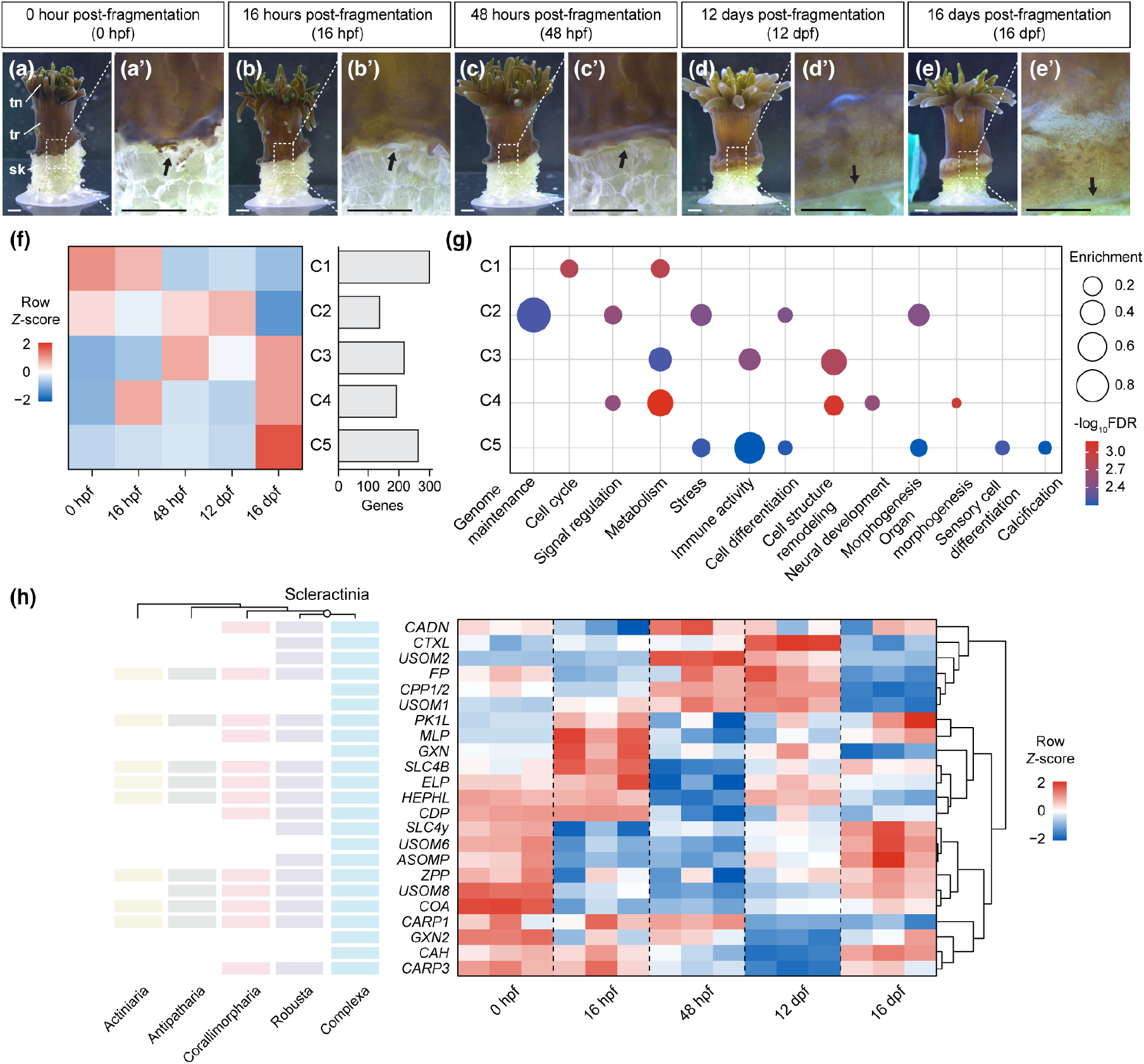
Bulk transcriptomics reveal ancestral roles of skeletal organic matrix protein genes later co-opted for calcification. (a–e) Representative images of *G. fascicularis* polyp regeneration following colony fragmentation at five time points: 0, 16, and 48 hours post-fragmentation, and 12 and 16 days post-fragmentation. (a’–e’) Enlarged views of the boxed regions highlighting regeneration. Black arrows indicate the boundary of regenerating tissues. tn, tentacles; tr, trunk; sk, skeleton. Scale bars: 1 mm (low magnification) and 500 µm (high magnification). (f) Heatmap of 1,087 differentially expressed genes during the regeneration process following fragmentation, clustered into five groups with values representing the mean expression per cluster (C1–C5). Bar plot on the right shows the number of genes in each cluster. (g) Dot plot of enriched biological process gene ontology biological process terms among the top 50 differentially expressed genes in each cluster (C1–C5). (h) Heatmap showing expression profiles of the 23 skeletal organic matrix proteins (SOMPs) identified in *G. fascicularis* across the regeneration time course. The left panel indicates the presence–absence of these SOMPs across representative hexacorallians. The 23 SOMPs analysed in (h) include: *ASOMP* (acidic skeletal organic matrix protein), *CAH* (carbonic anhydrase), *CADN* (coadhesin), *CARP1/3* (coral acid-rich proteins 1 and 3), *CDP* (CUB domain protein), *COA* (collagen α chain), *CPP1/2* (CUB and peptidase domain proteins 1 and 2), *CTXL* (cephalotoxin-like protein), *ELP* (EGF-laminin G domain protein), *FP* (neuroglian FNIII domain protein), *GXN/GXN2* (galaxins 1 and 2), *HEPHL* (hephaestin-like protein), *MLP* (mucin-like protein), *PK1L* (polycystin-1-related protein), *SLC4β/γ* (anion exchangers β/γ), *USOM1/2/6/8* (uncharacterized SOMPs), and *ZPP* (ZP domain protein).

**Figure 6.**
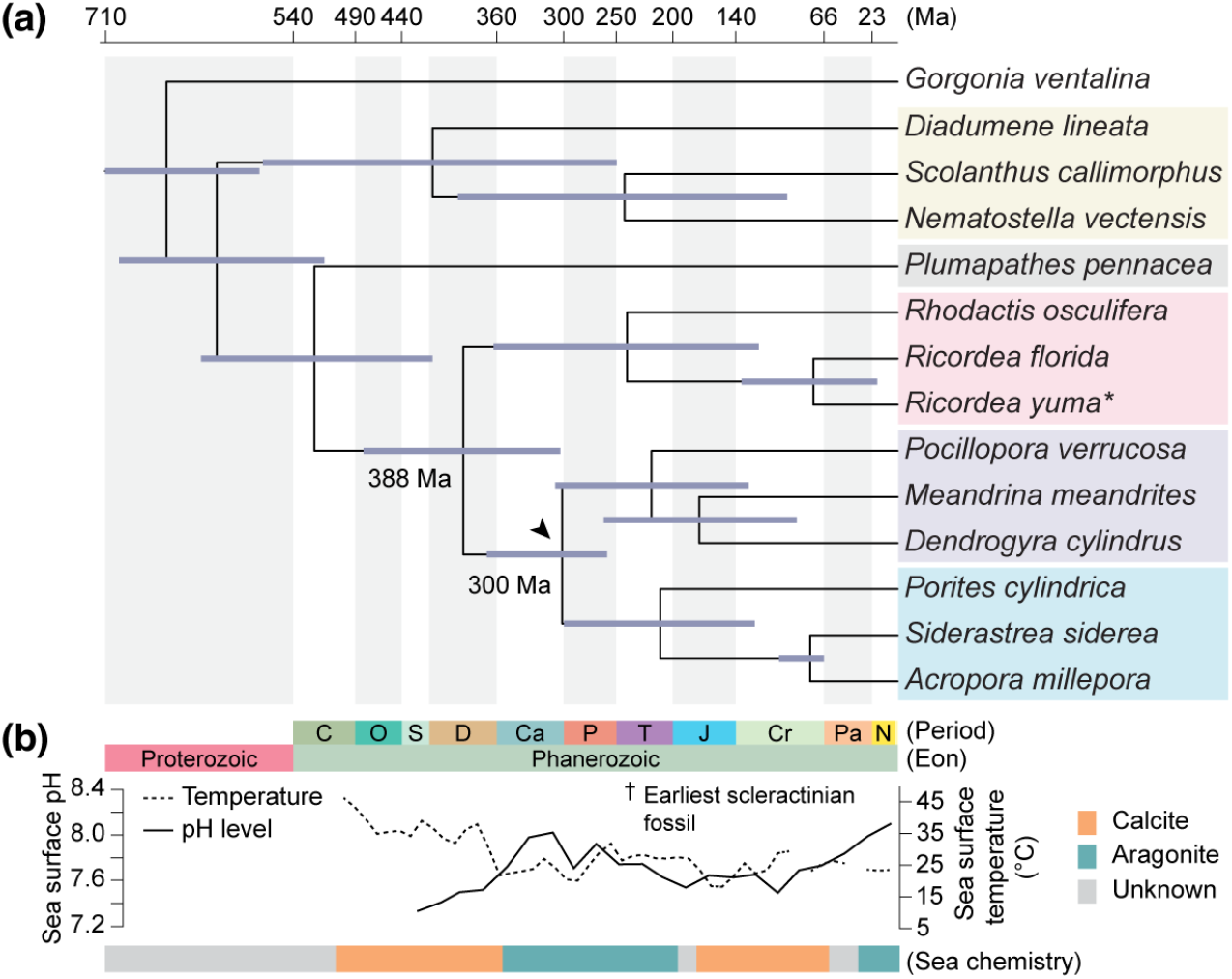
Time-calibrated phylogeny supports an origin of coral calcification in aragonite-favouring oceans. (a) Time-calibrated phylogenetic tree of selected hexacorallians, inferred with MCMCTree. The tree was initially constructed using single-copy orthologs with 100% species occupancy (*n* = 2,335) and the Q.PFAM+F+R6 model in IQ-TREE 3. Light blue bars indicate 95% highest posterior density intervals for node ages. Arrowhead indicates the possible origin of the coral skeleton during an aragonite sea interval, pre-dating the earliest fossil evidence of skeleton formation. Ma, million years ago. The genome sequenced in this study is marked with an asterisk. (b) Reconstruction of past ocean climates in alignment with the time-calibrated phylogeny. Sea surface temperature (dotted line) is reconstructed from δ^18^O models of Phanerozoic fossils (Grossman and Joachimski 2022). Model-predicted ocean surface pH (black line) is based on atmospheric CO_2_ and seawater Ca^2+^ reconstructions (Ridgwell and Zeebe 2005). Periods of aragonite (blue) versus calcite (orange) seas are indicated from Mg/Ca ratios (Balthasar and Cusack 2015), with gray boxes marking intervals of uncertainty. Geological timescale abbreviations: C, Cambrian; O, Ordovician; S, Silurian; D, Devonian; Ca, Carboniferous; P, Permian; T, Triassic; J, Jurassic; Cr, Cretaceous; P, Paleogene; N, Neogene. Dagger indicates the earliest fossil evidence of a scleractinian skeleton.

Of the 23 *G. fascicularis* SOMPs that were differentially expressed, approximately half were most highly expressed at 0 hours post-fragmentation (hpf) (Fig. 5h). This suggests that they are genes utilised during baseline coenosarc tissue functioning and maintenance rather than regeneration. The remaining SOMPs were not partitioned into a single cluster and instead showed highly variable expression patterns across the regeneration timecourse, with subsets upregulated in stages, at 16 hpf, 48 hpf, and 16 days post-fragmentation. This implies that calcification did not arise through wholesale co-option of an existing regulatory program (such as that underlying regeneration) but that SOMPs were integrated into the calcification programme through the recruitment of genes with diverse ancestral roles. Together, our genomic, single-cell, and regeneration analyses support a model in which coral biomineralization evolved through the assembly of a novel calicoblast cell state in a subset of epidermal cells utilising newly emerged scleractinian-specific genes and selectively co-opted ancestral hexacorallian components.

### The emergence of the scleractinian skeleton coincided with a shift to aragonite-favouring ocean chemistry

Finally, we used geological conditions to further explore the timing of the emergence of coral skeletonization. To this end, we first generated a maximum likelihood phylogenetic tree of our hexacorallian dataset in IQ-TREE 3 (Minh et al. 2020; Wong et al. 2026) with 2,335 single-copy orthologues and then performed molecular clock analyses using MCMCTree (Yang and Rannala 2006; Rannala and Yang 2007; dos Reis and Yang 2011) (Fig. 4a, Figure S19). This placed the divergence between Corallimorpharia and Scleractinia at approximately 388 million years ago (Ma) and the divergence of complex and robust scleractinians at approximately 300 Ma (Fig. 4a). Comparative analyses across biomineralizing animals suggest that skeletal mineralogy often reflects the prevailing seawater chemistry at the time skeletons first evolved, with few subsequent transitions despite later fluctuations in ocean chemistry (Porter 2010). Reconstructions of oceanic chemistry (Balthasar and Cusack 2015) place the divergence of corallimorpharians and scleractinians during a time of calcite-favouring chemistry but the origin of crown group scleractinians in an aragonite-favouring period that persisted through to the end of the Triassic (Fig. 4b). This supports the hypothesis that scleractinians’ aragonite skeletons emerged at the base of stony corals rather than in the ancestor of corallimorpharians and scleractinians. Taken with our single-cell transcriptomic data, it suggests that calcification is a synapomorphy of scleractinians and was not present in the ancestor of corallimorpharians.

## Discussion

Resolving the phylogenetic position of corallimorpharians is central to understanding both the initial emergence and potential for loss of calcification in hexacorallians. Early mitochondrial protein analyses supported the naked coral hypothesis, in which corallimorpharians were interpreted as skeleton-lost corals nested within Scleractinia (Medina et al. 2006; Kayal et al. 2013), but subsequent studies based on nuclear genes, mitochondrial nucleotide sequences, and phylogenomic datasets have increasingly recovered Corallimorpharia as the sister group to a monophyletic Scleractinia (Fukami et al. 2008; Kitahara et al. 2014; Lin et al. 2016; Wang et al. 2017; Kayal et al. 2018; Quattrini et al. 2018; Quattrini et al. 2020; McFadden et al. 2021; Wang et al. 2021; Quek et al. 2023; Vaga et al. 2024; Vaga et al. 2025). Our macrosynteny and microsynteny analyses each provide sequence-independent support for scleractinian monophyly. Though there are limitations to macrosynteny-based phylogenetic inferences, such as the potential for homoplasy (Kulkarni et al. 2026; Struck 2026) and limitations in the statistical approaches to determining what constitutes a ‘significant’ ALG (Copley 2025; Struck 2026), we can rule out homoplasy due to common derived features of the C1●M fusion in complex and robust corals, and the highly conserved nature of ALGs in hexacorallians eliminates any ambiguity over ALG assignments. The three chromosome fusion events shared by robust and complex corals therefore provide robust support for the monophyly of scleractinians. This places corallimorpharians outside the skeleton-forming group and rejects the hypothesis that they represent skeleton-lost stony corals. This means that corallimorpharians do not provide evidence of skeleton loss in response to ocean acidification and are unlikely to provide a direct evolutionary analog for how modern reef-building corals might respond to extreme ocean acidification.

Having established that corallimorpharians represent the nearest outgroup to skeleton-forming corals, we used comparative single-cell transcriptomics to explore the origins of calcification at cellular and molecular levels. Across orthology-based and orthology-independent integrations, calicoblasts formed a conserved transcriptional state shared by robust and complex corals but not corallimorpharians, sea anemones, or soft corals. Calicoblasts therefore represent a scleractinian-specific cell-type innovation associated with biomineralization rather than the retention of an ancestral anthozoan cell type. The limited but detectable similarity between calicoblasts and epidermal populations is consistent with their developmental origin as specialized ectodermal cells (Johnston 1980; Le Tissier 1990; Drake et al. 2020; Levy et al. 2021; Bonacolta et al. 2024; Han et al. 2025), yet our cross-species analyses suggest that calicoblasts have evolved a distinct cellular identity beyond a generalized epithelial state. Furthermore, though calicoblasts were conserved across scleractinian corals, distinct suites of transcription factors define them in robust and complex clades, suggesting that their regulatory programs continued to diverge following the origin of the cell type. Such divergence may have contributed to the emergence of lineage-specific skeletal traits observed among major coral clades (Budd and Stolarski 2011). Interpretation of these patterns remains tentative because currently available single-cell datasets, including our *G. fascicularis* atlas, differ in sequencing depth, cell recovery, and tissue dissociation efficiency. Broader taxonomic sampling together with optimized coral-specific single-cell protocols will be required to determine the extent and evolutionary significance of regulatory diversification in calicoblasts.

Comparative analyses of SOMPs indicate that the biomineralization toolkit used by calicoblasts was assembled through both lineage-specific innovation and the co-option of ancestral genes, consistent with previous hypotheses regarding the evolutionary assembly of coral biomineralization machinery (Ramos-Silva et al. 2013). Several key components, including SLC4γ, galaxins, skeletal aspartic acid-rich proteins and some carbonic anhydrases, are restricted to scleractinians and likely originated in association with skeleton formation. SLC4γ provides a particularly clear example, having arisen through tandem duplication of the ancestral SLC4β transporter before becoming functionally integrated into the coral biomineralization program (Zoccola et al. 2015; Tinoco et al. 2023). By contrast, many other skeletal organic matrix proteins are shared with corallimorpharians and broader hexacorallians, where they likely fulfilled ancestral epithelial and extracellular functions prior to their recruitment into calcification. Notably, our bulk RNA-sequencing data suggest that many of these non-scleractinian-specific components remain active during baseline tissue maintenance and at various stages of polyp regeneration. Coral biomineralization therefore appears to have evolved not through the invention of an entirely new molecular system, but through the integration of lineage-specific innovations into pre-existing cellular machinery (Arendt et al. 2016). This pattern is consistent with a broader evolutionary principle in which novel traits emerge through the redeployment and reorganization of ancestral developmental programmes rather than through *de novo* invention (True and Carroll 2002; Shubin et al. 2009).

Finally, we examined when in geological time the scleractinian skeleton first emerged and whether its origin was associated with major environmental transitions. Our molecular clock analyses support a deep Palaeozoic divergence between Corallimorpharia and Scleractinia, followed by the later emergence and ecological expansion of skeletonized corals. While our estimated stem age is consistent with previous phylogenomic reconstructions (e.g., 383 Ma) (Quattrini et al. 2020), our crown age is younger than estimates derived from dense, order-wide taxonomic sampling, which place the crown radiation between the Carboniferous (332–357 Ma) (Quattrini et al. 2020) and the Ordovician (∼460 Ma) (Vaga et al. 2025). This discrepancy is expected since our focused taxon sampling strategy prioritized high-quality genomic resolution over exhaustive phylogenetic density, which generally produces deeper node age estimates (Chen et al. 2025). This notwithstanding, the temporal pattern is broadly consistent with previous phylogenomic studies, indicating that the stem lineage of Scleractinia predates the earliest unequivocal coral skeleton fossils by tens to hundreds of millions of years (Stanley 2003; Stolarski et al. 2011; Campoy et al. 2020; Quattrini et al. 2020; Campoy et al. 2023; Vaga et al. 2025). Recent phylogenetic analyses suggest that early scleractinians within this interval were likely solitary, azooxanthellate corals inhabiting deep-water refugia during Paleozoic environmental crises (Campoy et al. 2023). This evolutionary interval also overlapped with major long-term shifts in ocean chemistry, including declining atmospheric CO_2_, increasing ocean pH, and the transition towards aragonite-favoring oceans (Porter 2010; Stolarski et al. 2011). Although these correlations do not establish direct environmental causation, they suggest that the ecological radiation of skeletonized scleractinians occurred under conditions increasingly permissive for aragonite biomineralization. Our results therefore support a scenario in which components of the calcification machinery evolved along the scleractinian stem lineage before the later ecological expansion of heavily skeletonized reef corals during Triassic aragonite seas.

In summary, we present a chromosome-level genome for the corallimorpharian *R. yuma* and integrate it with available corallimorpharian and hexacorallian genomes to resolve Corallimorpharia as the sister group to a monophyletic Scleractinia. This rejects the hypothesis that corallimorpharians are secondarily non-calcifying stony corals and provides an evolutionary context for reconstructing the origin of coral calcification. By generating comparative single-cell datasets from a corallimorpharian and a reef-building coral, we show that the scleractinian skeleton evolved through the emergence of a derived calicoblast cell state together with a biomineralization toolkit assembled from lineage-specific innovations and co-opted ancestral components. Although corallimorpharians retain some ancestral genes later incorporated into coral biomineralization, they lack a detectable calicoblast-like transcriptional state. Together, our findings indicate that coral calcification arose through coordinated changes in genome architecture, gene repertoire, and cell identity, highlighting this process as a major evolutionary transition driven by the integration of genomic and cellular innovations.

## Materials and methods

### Specimen collection and genomic sample preparation

Specimens of the native Indo-Pacific corallimorpharian *Ricordea yuma* were collected from Kenting, Taiwan, and reared in a tank at the National Museum of Marine Biology and Aquarium (NMMBA), Pingtung, Taiwan. Two individual *R. yuma* polyps from the same colony were maintained in the lab in seawater filtered through 1-μm filters, where they reproduced asexually by budding or fragmentation. Genomic DNA and Hi-C nuclei samples were prepared from menthol-bleached polyps collected from the same colony to minimize genetic variation and remove symbiotic dinoflagellates. Aposymbiotic polyps were produced following menthol treatment at 200 mM for approximately one week (J.-T. Wang et al. 2012). High molecular weight DNA was extracted using DNAzol Reagent (Thermo Fisher Scientific 10503027), followed by the manufacturer’s protocol. For Hi-C library preparation, *R. yuma* polyps were dissected into small fragments and dissociated in calcium- and magnesium-free seawater (CMF-ASW) to obtain a single-cell suspension. The resulting cells were snap-frozen in liquid nitrogen and stored at −80 °C until further processing.

### Genome sequencing and assembly

To achieve a chromosome-level genome assembly of the corallimorpharian *R. yuma*, we combined Pacific Biosciences long-read sequencing with long-range chromatin interaction data from Phase Genomics Hi-C. Genomic DNA concentration was measured using a Qubit 4 Fluorometer (Thermo Fisher Scientific), and DNA size distribution and concentration were further verified with a Fempto Pulse System using the Genomic DNA 165 kb Kit (Agilent). The whole-genome sequencing library was prepared using the Genomic DNA SMRTbell Prep Kit (v3.0), and the SMRTbell library was size-selected (5–50 kb) using BluePippin 0.75% agarose gel cassettes (Sage Science). A total of 25.0 Gb of HiFi long reads (∼43-fold coverage) were generated from a single SMRT flow cell on the PacBio Sequel IIe platform. Hi-C libraries were prepared from frozen nuclei samples using the Phase Genomics Proximo Hi-C Animal Kit (KT2045), and 148 million read pairs of Hi-C data were sequenced on the Illumina NextSeq 2000 platform (150 bp paired-end). The PacBio HiFi reads were assembled into contigs using Hifiasm (v0.20.0-r639) (Cheng et al. 2021), with redundancy removed by Purge_Dups (v1.2.5) (Guan et al. 2020) to collapse haplotypic sequences. Hi-C data were integrated to scaffold the contigs into chromosome-scale pseudomolecules using YaHS (v1.2a.1) (Zhou et al. 2022). Manual curation was performed using Juicebox (v1.11.08) (Durand et al. 2016) to correct misjoins and refine contig orientation. Genome statistics were calculated with Gfastats (v1.3.9) (Formenti et al. 2022). Genome assembly completeness was assessed using BUSCO (v6.0.0) against the metazoa_odb10 datasets (Simão et al. 2015).

### Transcriptome sequencing for genome annotation

To generate a reference transcriptome for gene annotation, nine *R. yuma* polyps, all asexually derived from a single parent colony maintained at the NMMBA and therefore assumed to share an identical genotype, were acclimated in a closed-circuit tank containing 10 L of filtered seawater for seven days under controlled conditions (25 °C, salinity 3.5%, pH 8.0, 12:12 h light cycle). Following acclimation, the polyps were divided into three groups of three individuals and exposed to different temperature conditions (25 °C, 28 °C, or 31 °C) for an additional seven days to maximize transcript diversity and capture a broader range of gene expression profiles for annotation. Tissue samples were collected using RNase AWAY-treated scissors (Thermo Fisher Scientific) and snap-frozen in liquid nitrogen for RNA extraction. Total RNA was extracted using TRIzol reagent (Thermo Fisher Scientific) according to the manufacturer’s instructions. For transcriptome sequencing, cDNA libraries were prepared from 100 ng of total RNA using the Illumina Stranded mRNA Prep Kit and sequenced on the Illumina NovaSeq 6000 platform with 150 bp paired-end reads.

### Genome dataset curation and gene prediction

We curated a dataset of exclusively chromosome-level genomes for phylogenomic and synteny analysis (Dataset S1). The dataset consisted of one octocoral outgroup (*Gorgonia ventalina*), three members of Actiniaria (sea anemones: *Diadumene lineata (Wood et al. 2022)*, *Nematostella vectensis (Fletcher et al. 2023)*, and *Scolanthus callimorphus* (Zimmermann et al. 2023); one member of Antipatharia (black corals: *Plumapathes pennace*a); three members of Corallimorpharia (coral-like anemones: *Rhodactis osculifera*, *Ricordea florida* and our new *Ricordea yuma* genome); three members of the scleractinian clade Robusta (*Dendrogyra cylindrus*, *Meandrina meandrites,* and *Pocillopora verrucosa*); and three members of the scleractinian clade Complexa (*A. millepora (Fuller et al. 2020)*, *Porites cylindrica*, and *Siderastrea siderea*). Ten of the 14 genomes were sequenced as part of the Aquatic Symbiosis Genomics (AGS) Project of the Tree of Life Programme (Threlfall and Blaxter 2021; The Darwin Tree of Life Project Consortium et al. 2022; McKenna et al. 2024).

Repeat and gene prediction were performed following our previously published pipeline (Lewin et al. 2024; Lewin et al. 2026). Briefly, repeats were annotated de novo with RepeatModeler (v2.0.5) (Flynn et al. 2020) and then soft-masked with RepeatMasker (v4.1.5) (Smit et al. 2015). Gene prediction was performed using BRAKER3 (v3.0.8) (Stanke et al. 2006; Stanke et al. 2008; Li et al. 2009; Barnett et al. 2011; Lomsadze et al. 2014; Buchfink et al. 2015; Hoff et al. 2016; Hoff et al. 2019; Brůna et al. 2021; Gabriel et al. 2024). RNA sequencing data was available for all species except *Scolanthus callimorphus*. In these cases, RNA sequencing data (Table S4) was downloaded with fastq-dump (v3.1.1) (Leinonen et al. 2011), trimmed with fastp (v0.23.4) (Chen et al. 2018), mapped to the soft-masked genome with HISAT2 (v2.2.1) (Kim et al. 2019) and fed to BRAKER3 as hints in .bam format. OrthoDB (Kuznetsov et al. 2023) protein hints were used for *S. callimorphus* in the absence of RNA sequencing data.

### Macrosynteny analysis

The SyntenyFinder (v1.0) (Lewin, Liao, Chen, et al. 2025) pipeline was used to perform macrosynteny-based analyses. SyntenyFinder identifies single-copy orthologs using OrthoFinder (Emms and Kelly 2019) and assigns them to one of the 21 cnidarian ancestral linkage groups (Simakov et al. 2022) based on orthology. It then uses an R (v4.3.0) script (R Core Team 2024) to perform pairwise analyses of macrosynteny using dot plots and ribbon plots. RIdeogram (v0.2.2) (Hao et al. 2020) is used to construct ribbon plots. Dot plots show all single-copy orthologs; ribbon plots trim links between chromosomes with fewer than 10 shared genes for clarity. Due to our use of relatively closely related species, inference of fusions is not reliant on a small number of genes per ALG, a known problem with previous studies (Schultz et al. 2023; Copley 2025), and pairwise comparisons consist of an average of 2,161 genes with a known ALG (approximately 103 per ALG) (Table S2).

### Microsynteny analysis

First, the presence of conserved microsynteny, referring to small-scale gene order within a chromosome, was quantified using GENESPACE (v1.3.1) (Lovell et al. 2022). GENESPACE uses OrthoFinder (Emms and Kelly 2019) to determine homology and MCScanX (Y. Wang et al. 2012) to detect conserved blocks of collinear genes across genomes. GENESPACE was run in R with a block size of 5 to measure the proportion of homologues in collinear blocks in pairwise comparisons of species. Plots were produced with ggplot2 (v3.5.1) (Wickham 2009). Second, the network-based approach of syntenet (v1.4.2) (Almeida-Silva et al. 2023) was used to identify conserved taxon-specific microsyntenic gene blocks. Finally, microsynteny-based phylogenetic inference was performed with IQ-TREE 3 (v3.0.1) using a binarized matrix representing the presence/absence of microsyntenic gene clusters found by syntenet. *S. callimorphus* had a large number of species-specific duplications and was therefore removed from the dataset prior to tree building.

### Coral polyp isolation and culture conditions

Experimental validation of calcification-associated gene expression was conducted in the reef-building coral *Galaxea fascicularis*, originally collected from the northeast coast of Taiwan and maintained under an outdoor pond (Shikina et al. 2024). As one of the species included in our single-cell atlas, *G. fascicularis* possesses a distinct calcifying cell population (the calicoblasts) and was selected for functional validation of candidate calcification genes identified in our comparative analyses of biomineralizing cell types. Its ability to propagate through polyp separation (Puntin et al. 2022) and suitability for long-term laboratory culture enabled controlled experimental studies. Coral colonies approximately 5 cm in diameter were kept under controlled conditions in an indoor fibre-reinforced plastic tank at the New Taipei City Marine Resource Restoration Park. For sample preparation, individual polyps were gently excised from colonies using bone cutters. Polyps were attached to 0.1-mm-thick film sheets using instant adhesive and transferred to glass containers containing 800 mL of artificial seawater, with up to 30 polyps per container. Cultures were maintained under a 12 h light/12 h dark photoperiod with a light intensity of 100 μE/m^2^s, at 25 °C and 35 ppt salinity. Coral husbandry, including *Artemia* feeding and complete seawater replacement, was performed twice weekly as described previously (Shikina et al. 2023).

### Single-cell preparation and RNA sequencing

Whole polyps of *R. yuma* and *G. fascicularis* were dissected into small tissue fragments, and excess seawater was carefully removed prior to dissociation. All dissociation and fixation steps were performed in 15 ml Falcon tubes pre-coated with 1% BSA/PBS to minimize cell adhesion. Tissue fragments were then incubated for 5 min in 10 ml calcium- and magnesium-free artificial seawater (CMF-ASW; 450 mM NaCl, 9 mM KCl, 30 mM Na_2_SO_4_, 2.5 mM NaHCO_3_, 10 mM Tris-HCl, 2.5 mM EDTA, and 25 mM HEPES; adjusted to pH 8.0 using 10 N NaOH) to deplete residual calcium and magnesium ions. Following incubation, CMF-ASW was removed and replaced with 1 ml fresh CMF-ASW, after which tissue fragments were gently dissociated by repeated pipetting using low-retention P1000 pipette tips to generate single-cell suspensions. Cell suspensions were filtered through a 40 μm cell strainer (Corning 431750) to remove aggregates prior to fixation.

Filtered cell suspensions were collected into fresh 15 ml Falcon tubes pre-coated with 1% BSA/PBS and fixed using a modified ACME protocol (García-Castro et al. 2021), an acetic acid-and methanol-based fixation method compatible with fluorescence-activated cell sorting (FACS) and downstream single-cell RNA sequencing. Briefly, 1 ml cell suspension in CMF-ASW was mixed with 9 ml freshly prepared ACME solution (1.5 ml methanol, 1 ml glacial acetic acid, 1 ml glycerol, and 0.2 ml 7.5% *N*-acetyl-L-cysteine). Samples were incubated for 20 min at room temperature with gentle rocking, followed by centrifugation at 1,000 × *g* for 5 min at 4 °C using a swinging-bucket rotor to pellet cells and remove the fixative. Cell pellets were washed once with 1% BSA/PBS and resuspended in 1% BSA/PBS supplemented with 10% DMSO for cryopreservation at −80 °C until FACS sorting.

Prior to FACS sorting, frozen fixed-cell suspensions were thawed and centrifuged at 1,000 × *g* for 5 min at 4 °C. Cell pellets were resuspended in chilled 1% BSA/PBS at a final concentration of approximately 2 × 10^6^ cells/ml and passed through a 40 μm cell strainer immediately before sorting to remove residual aggregates. FACS acquisition and sorting were performed on a BD FACSAria IIIu (BD Biosciences) using BD FACSDiva software (v8.0.3). Sorting was conducted in 4-way purity mode using a 100 μm nozzle at a sheath pressure of 30 psi and a drop drive frequency of 41,000 Hz. Both sample and collection chambers were maintained at 4 °C throughout the procedure. Initial gating based on forward scatter area (FSC-A) and side scatter area (SSC-A) was used to exclude debris and retain intact cellular events. Doublets and aggregates were further excluded using FSC-H versus FSC-W and SSC-H versus SSC-W gating.

To separate animal cells from symbiotic algae, chlorophyll autofluorescence was detected in the PerCP-Cy5.5 channel following excitation with a 488 nm laser. Symbiodiniaceae cells exhibited strong autofluorescence, whereas animal cells displayed substantially lower fluorescence intensity (Figure S6). Animal cells were therefore isolated as FSC/SSC-positive, low-autofluorescence singlet events. Sorted animal cells were collected into chilled Eppendorf Protein LoBind tubes (Eppendorf 022431102) containing 1% BSA/PBS supplemented with RNase inhibitor (Thermo Scientific N8080119) at a final concentration of 2 U/μl. Following FACS sorting, cells were centrifuged at 1,000 × *g* for 5 min at 4 °C, resuspended in chilled 1% BSA/PBS, and mixed with trypan blue for cell counting and morphological quality assessment using the EVOS M7000 Imaging System (Thermo Scientific).

Single-cell RNA sequencing was performed using the 10x Genomics Chromium platform. Single-cell libraries were generated using the Chromium GEM-X Single Cell 3′ v4 platform (PN-1000691), including the Chromium GEM-X Single Cell 3′ GEM Kit v4 (PN-1000693), Library Construction Kit (PN-1000694), Gel Bead Kit v4 (PN-1000692), Chromium GEM-X Single Cell 3′ Chip Kit v4 (PN-1000690), and Dual Index Kit TT Set A (PN-1000215), following the manufacturer’s instructions. Cell suspensions were loaded targeting recovery of 20,000 cells per sample. The *R. yuma* and *G. fascicularis* suspensions had final cell concentrations of approximately 950 and 1,800 cells/μl, respectively, and both samples passed quality control with low aggregate levels prior to GEM generation. Libraries were sequenced on an Illumina NextSeq 2000 platform using XLEAP-SBS chemistry with a paired-end sequencing configuration of 28 bp for Read 1, 10 bp for i7 index reads, 10 bp for i5 index reads, and 90 bp for Read 2 on a P4 flow cell. The *R. yuma* and *G. fascicularis* libraries generated 88,113 Mb and 83,355 Mb of sequencing data.

### Single-cell transcriptomic analysis

Single-cell RNA-seq raw reads were processed with Cell Ranger (v10.0.0) using --include-introns=true and --expect-cells=20000. Since 10x Genomics libraries are enriched for transcript 3′ ends, incomplete annotation of terminal exons and 3′ UTRs can substantially reduce transcript recovery in non-model genomes and bias comparative single-cell analyses across species. To improve transcript assignment and recover reads mapping to incompletely annotated 3′ regions, we refined the *R. yuma* annotation using GeneExt (Zolotarov et al. 2026) together with BRAKER3-derived gene models. GeneExt leverages single-cell RNA-seq coverage profiles to extend and homogenize transcript annotations, thereby improving recovery of biologically informative transcripts from poorly annotated loci.

Illumina sequencing generated 746,719,665 reads for *R. yuma* and 706,399,235 reads for *G. fascicularis*. *R. yuma* reads were mapped against this optimized annotation set, yielding 11,003 estimated cells with a mean of 67,865 reads per cell. Overall, 35.3% of reads mapped to the genome and 25.9% were assigned to annotated transcripts, corresponding to 33,071 detected genes. *G. fascicularis* reads were mapped against a Trinity-derived transcriptome reference, yielding 16,202 estimated cells with a mean of 43,600 reads per cell. Overall, 71.9% of reads mapped to the genome and 47.5% were assigned to annotated transcripts, corresponding to 29,892 detected genes.

To minimize confounding effects from ambient RNA contamination, which commonly arises from cell-free transcripts captured during droplet-based single-cell RNA-seq, raw count matrices were corrected using SoupX (v1.6.2) (Young and Behjati 2020). Contamination fractions were estimated automatically with the autoEstCont function following preliminary SCTransform-based clustering using 30 principal components and a Louvain resolution of 0.1. Corrected count matrices were subsequently analyzed in Seurat (v5.3.0) (Hao et al. 2024). Cells were filtered using species-specific thresholds based on the number of detected features, total UMI counts, and mitochondrial read fraction (*R. yuma*: nFeature 223–1,658, nCount 326–2,347, mitochondrial reads <0.51%; *G. fascicularis*: nFeature 192–1,612, nCount 657–5,342, mitochondrial reads <10%).

Data were normalized with SCTransform (v2) using glmGamPoi (v1.24.0) (Ahlmann-Eltze and Huber 2021) while regressing mitochondrial read percentage, followed by principal component analysis, shared nearest-neighbor graph construction, Louvain clustering, and UMAP embedding. Final datasets comprised 9,822 cells and 27,168 genes for *R. yuma* and 12,750 cells and 28,059 genes for *G. fascicularis*. Clustering identified ten major cell types in each species (*R. yuma*: 25 principal components, resolution 0.1; *G. fascicularis*: 30 principal components, resolution 0.1).

### Subsampling analysis of calicoblast marker detectability

To assess whether the absence of calicoblast marker genes in *R. yuma* could be explained by technical limitations rather than a true biological difference, a subsampling simulation was performed using single-cell RNA sequencing datasets from three scleractinian corals: *S. pistillata*, *O. patagonica*, and *A. millepora (Levy et al. 2025)*. Since the *R. yuma* dataset contained 9,822 cells, this cell number was included as one of the subsampling levels for comparison. Cells were randomly subsampled to target cell numbers ranging from 200 to 20,000. Unique molecular identifier (UMI) counts in calicoblast cells were then downsampled to match the mean sequencing depth of *R. yuma* (885.3 UMIs per cell). This procedure was repeated 50 times for each cell-count level, with a fixed random seed (seed = 42). Calicoblast marker genes were defined as genes with log_2_ fold-change > 1, expression detected in >10% of cells, and *P* < 0.05. In each iteration, calicoblast marker genes with a summed UMI count >10 across all calicoblast cells were considered detectable, and the proportion of detectable marker genes was calculated. Data are expressed as mean ± SD.

### Cross-species single-cell analysis

Publicly available single-cell RNA sequencing datasets for *S. pistillata*, *O. patagonica*, and *A. millepora* were retrieved from NCBI GEO (accession GSE289546) (Levy et al. 2025). Single-cell data for *A. muricata* were obtained from Figshare (Han et al. 2025). All datasets were analyzed using the original processed count matrices and cell-type annotations, including *S. pistillata* (15,053 cells, 49,751 genes, 25 annotated subtypes), *O. patagonica* (29,723 cells, 51,131 genes, 31 annotated subtypes), *A. muricata* (4,941 cells, 12,916 genes, 13 annotated subtypes), and *A. millepora* (28,736 cells, 43,454 genes, 27 annotated subtypes). To facilitate cross-species comparisons, fine-grained subtype annotations were consolidated into broader biological categories, including calicoblasts, epidermal cells, algae-hosting cells, gastrodermal cells, muscle cells, digestive filament cells, gland cells, neurons, cnidocytes, immune cells, and germline/progenitor cells.

Cell types were annotated through cross-species correlation analyses with previously published coral single-cell RNA-seq datasets using nine independent pairwise SAMap (v2.0.0) comparisons (Tarashansky et al. 2021), implemented with SAMalg (v2.0.0), scanpy (v1.11.5), anndata (v0.12.10), and scipy (v1.17.1). *R. yuma* and *G. fascicularis* were each independently compared with *S. pistillata*, *O. patagonica*, *A. muricata*, and *A. millepora*, together with a direct comparison between *R. yuma* and *G. fascicularis*. Each comparison was performed independently (pairwise=True) from unmodified species-specific h5ad files to avoid cross-run interference and preserve species-specific transcriptomic structure during pairwise alignment.

Gene homology bridges between species were constructed using reciprocal best-hit (RBH) searches implemented in DIAMOND (v2.1.11) (Buchfink et al. 2021). Because evolutionary distances varied substantially among the compared cnidarian species, sequence similarity thresholds were adjusted to maximize recovery of biologically meaningful orthologs while minimizing spurious matches. Coral–coral comparisons were conducted using the sensitive search mode with a minimum amino acid identity threshold of 60%, whereas comparisons involving *R. yuma* employed the more-sensitive mode with a 40% identity threshold. Searches used an *E*-value cutoff of ≤1e−6 and retained a single target per query. Only reciprocal matches recovered in both search directions were retained as RBH orthologs. The numbers of RBH ortholog pairs recovered across pairwise comparisons ranged from 3,164 to 6,866 genes. Pairwise cell-type alignment scores (0–1) were calculated using the SAMap get_mapping_scores function and visualized as heatmaps after collapsing fine-grained subtypes into broad biological categories.

For sequence-independent cell-type comparisons (i.e., orthology-independent integration), TranscriptFormer (v0.6.1) was used with the pre-trained TF-Metazoa model, which contains 444 million trainable parameters and was trained on 112 million cells across 12 species (Pearce et al. 2026). TranscriptFormer generates per-cell embeddings by jointly encoding gene expression profiles and gene identity represented by protein embeddings derived from Evolutionary Scale Model 2 (ESM-2), a transformer-based protein language model trained on large-scale protein sequence data (Lin et al. 2023). Protein embeddings were generated *de novo* for all cnidarian species analyzed in this study using predicted protein sequences.

To quantify transcriptomic similarity among cell types across species, pairwise cosine similarities between cell-type centroids were computed. For each species, centroid embeddings were calculated by averaging embeddings across all cells assigned to the same cell type, followed by L2 normalization. A consensus calicoblast centroid was constructed by pooling calicoblasts from four scleractinian coral species, including *S. pistillata* (629 cells), and *O. patagonica* (1,195 cells), *G. fascicularis* (724 cells), and *A. millepora* (1,483 cells). To reduce species-specific biases, embedding vectors from each species were independently mean-centered prior to pooling. For broader cnidarian cross-species comparisons, publicly available single-cell RNA sequencing datasets from the sea anemone *N. vectensis (Sebé-Pedrós et al. 2018)* and the soft coral *Xenia* sp. (Hu et al. 2020) were included. The *N. vectensis* and *Xenia* sp. datasets were retrieved from NCBI GEO (GSE95723) and BioProject (PRJNA548325), respectively, and comprised 20,808 cells across 32 annotated cell types and 25,271 cells across 24 annotated cell types.

Calicoblast-enriched genes were identified independently for each of the four scleractinian coral species using the FindMarkers function implemented in Seurat with a Wilcoxon rank-sum test (calicoblasts versus all other cell types). Genes were retained if they showed an average log_2_ fold-change > 1, expression in >10% of calicoblast cells, and an adjusted *P*-value < 0.05. Transcription factors were annotated based on KEGG Orthology assignments generated by the KEGG Automatic Annotation Server using an RBH approach (Moriya et al. 2007). Transcriptome age indices (TAI) were calculated using GenEra (v1.4.2) and myTAI (v1.0.1.9000) (Drost et al. 2018; Barrera-Redondo et al. 2023). GO terms were assigned with EggNOG-mapper (v2.1.3) (Cantalapiedra et al. 2021), and GO enrichment analysis was conducted using topGO (v2.62.0) (Alexa et al. 2006).

### Skeletal organic matrix protein annotation

A set of 43 reference skeletal organic matrix proteins (SOMPs) was compiled from previous studies in *A. millepora* (38 genes) (Ramos-Silva et al. 2013) and *S. pistillata* (5 genes) (Drake et al. 2013). These reference sequences included both full-length and partial gene models. Orthology inference was conducted using three complementary approaches: reciprocal sequence similarity, phylogenetic analysis, and microsynteny conservation. First, each reference SOMP was used in a reciprocal BLASTP (v2.15.0) (Camacho et al. 2009) search against the predicted proteomes of all species included in this study (see Dataset acquisition and gene prediction). Candidate hits were retained based on reciprocal best hits and significant sequence similarity. For each SOMP, homologous sequences were then aligned using MAFFT (v7.525), and the resulting alignments were trimmed with trimAl (v1.5.0) (Capella-Gutiérrez et al. 2009) in automated mode. Phylogenetic relationships were reconstructed using IQ-TREE 3, enabling ModelFinder and performing ultrafast bootstrap and SH-aLRT support estimation. Phylogenetic trees were constructed for 32 SOMPs; in the remaining cases, only one or few species showed a significant hit, making tree construction unnecessary. To further assess orthology, microsynteny analysis was performed manually using the genomic coordinates and BRAKER3-predicted gene models for each species. A flexible window size was used depending on the local genomic context: in many cases, 10 genes upstream and downstream of each SOMPs candidate were sufficient, while in more complex or less conserved regions, up to 20–25 neighboring genes were compared to confidently identify syntenic blocks. A presence–absence matrix was constructed to summarize the detection of SOMPs across species. Genes with weak or ambiguous support (e.g., a BLAST hit with no clear phylogenetic or syntenic support) were recorded as uncertain.

The 38 annotated SOMPs include: ASOMP (acidic skeletal organic matrix protein), CA (carbonic anhydrase), CAH (putative carbonic anhydrase), CADN (coadhesin), CARP1 (coral acid-rich protein 1), CARP2/3/4 (coral acid-rich proteins 2, 3, and 4), CDP (CUB domain protein), COA (collagen α chain), CPP1/2 (CUB and peptidase domain proteins 1 and 2), CTXL (cephalotoxin-like protein), DQRP (Asp/Glu-rich protein), ECT (ectin), ELP (EGF– laminin G domain protein), FP (neuroglian FNIII domain protein), GXN/GXN2 (galaxins 1 and 2), HEPHL (hephaestin-like protein), MLP (mucin-like protein), MLRP1/2 (MAM and LDL-receptor domain proteins 1 and 2), MFP1/2 (MAM and FNIII domain proteins 1 and 2), PCDL (protocadherin-like protein), PK1L (polycystin-1-related protein), SAP1A/B and SAP2 (secreted acidic proteins), SAAR2 (skeletal Asp-rich protein 2), SLC4β/γ (anion exchangers β/γ), TRP (threonine-rich protein), USOM1–8 (uncharacterized SOMPs), and ZPP (ZP domain protein).

### Bulk RNA sequencing for regeneration experiments and transcriptome assembly

To generate a reference transcriptome for single-cell analysis, coenosarc tissues surrounding each *G. fascicularis* polyp, excluding the polyp body, were dissected under a stereomicroscope and collected for RNA extraction at designated time points (0, 16, and 48 h post-fragmentation, and 12 and 16 d post-fragmentation). Tissues were immediately transferred into 500 μL of TRI reagent (Sigma-Aldrich T9424) and homogenized, then stored at −20 °C. Total RNA was extracted using TRIzol reagent according to the manufacturer’s instructions. RNA libraries were prepared using the Illumina TruSeq Stranded mRNA Library Prep Kit following the standard protocol. Briefly, mRNA was isolated from 1 μg of total RNA using oligo(dT)-attached magnetic beads and fragmented under elevated temperature. First-strand cDNA synthesis was performed using reverse transcriptase and random primers, followed by second-strand synthesis and 3’ adenylation. Adaptors were ligated to the cDNA fragments, which were then PCR-amplified and purified using the AMPure XP system (Beckman Coulter). Library quality was evaluated using the Qsep400 System (Bioptic), and concentrations were measured with a Qubit 2.0 Fluorometer (Thermo Scientific). Sequencing was performed on an Illumina NovaSeq X Plus platform with 150 bp paired-end reads.

### Transcriptome assembly

A *G. fascicularis* transcriptome assembly was generated using Trinity (v2.15.2) (Grabherr et al. 2011; Henschel et al. 2012; Haas et al. 2013) in genome-guided *de novo* mode. Illumina NovaSeq 150 bp paired-end reads from one control and four time points (three biological replicates each; 458,400,255 read pairs after filtering) were quality-checked with FastQC (v0.12.1) and MultiQC (v1.21) (Ewels et al. 2016), then adapter- and quality-trimmed with fastp (v0.23.4) (Chen et al. 2018; Chen 2023). Trimmed reads were aligned to the *Galaxea fascicularis* genome (GenBank assembly GCA_948470475.1) using STAR (v2.7.11b), achieving a final alignment rate of 39.56%. The initial assembly comprised 337,241 transcripts (174,497 > 500 bp) with an average alignment fraction of 92.9% to the reference, and 28,862 putative chimeric transcripts, as determined by RNAQUAST (v2.3.0) (Bushmanova et al. 2016). Transcript filtering was performed with EvidentialGene (Gilbert 2016) to retain primary models.

### Bulk transcriptomic analysis

Transcript quantification was performed using Salmon (v1.10.3) (Patro et al. 2017) in quasi-mapping mode with automatic library type detection and a *k*-mer size of 31. The Salmon index was constructed from the filtered coding sequences generated by EvidentialGene. Individual quantification runs were executed in parallel for each sample using quality-filtered paired-end reads. The resulting quantification files were imported into R using the tximport (v.1.30.0) package. Differential expression analysis was carried out using DESeq2 (v1.42.1) (Love et al. 2014), applying a likelihood ratio test model suitable for time-course experiments. Genes with low expression were filtered using the filterByExpr function from the edgeR package (v4.0.16), which accounts for both library size and experimental group structure. Genes were considered differentially expressed if they exhibited an adjusted *P*-value < 0.01 and an absolute log_2_ fold-change (|LFC|) > 1.5. Differentially expressed genes were averaged and grouped into clusters using hierarchical *k*-means clustering, and expression values were averaged and *Z*-score normalized prior to visualization. Expression patterns across clusters were visualized in a heatmap using the pheatmap package (v1.0.12) in R.

To functionally characterize differentially expressed genes, we annotated the filtered proteome using EggNOG-mapper (v2.1.12) (Huerta-Cepas et al. 2019; Cantalapiedra et al. 2021), which assigned Gene Ontology (GO) terms to each predicted gene model. GO enrichment analysis was performed using topGO (v2.54.0) (Alexa et al. 2006), and statistical significance was evaluated using Fisher’s exact test. The top 50 enriched Biological Process (BP) GO terms for each cluster were manually curated to remove redundancy and merge semantically similar categories. A summary of enriched terms was visualized using a dotplot generated with ggplot2 (v3.5.2) To investigate the expression dynamics of SOMPs, we used the SOMP gene set generated in this study (see SOMP annotation methods). Homologous sequences were identified in the *G. fascicularis* predicted proteome using reciprocal BLASTP with an *E*-value threshold of 1e-5. SOMPs homologs were cross-referenced with the full DESeq2 output and filtered using an adjusted *P*-value threshold of < 0.05, without applying a fold-change cutoff. This broader filtering was used to include SOMPs not captured in the more strictly filtered and clustered DEG set. Expression profiles of the identified SOMPs were visualized as a clustered heatmap using pheatmap.

### Phylogenomics and divergence time estimation

For divergence time estimation, a tree was generated using chromosome-level genomes and IQ-TREE 3 (v3.0.1) (Minh et al. 2020; Wong et al. 2026). Single-copy orthologs with 100% species occupancy (*n* = 2,335) were identified with OrthoFinder (v2.5.4) (Emms and Kelly 2019). Protein sequences were then aligned with MAFFT (v7.520) (Katoh et al. 2002; Katoh and Standley 2013), trimmed with ClipKIT (v1.4.1) to maintain phylogenetically informative sites (Steenwyk et al. 2020), and concatenated to form a supermatrix with PhyKIT (v1.11.7) (Steenwyk et al. 2021). The phylogeny was inferred from the supermatrix by using ModelFinder (Kalyaanamoorthy et al. 2017) to select an optimal model for sequence evolution (model = Q.PFAM+F+R6) and then running IQ-TREE 3 on the complete concatenated alignment. UFBoot2 (Hoang et al. 2018) was used to perform 1,000 bootstrap replicates.

To test the robustness of this result to the tree-building methodology used, phylogenomic analysis was repeated using the same dataset with three further methods. First, a phylogeny was constructed with a partitioned analysis (-p option) using the phylogenetic terrace aware (PTA) data structure (Chernomor et al. 2016) in IQ-TREE 3. This permits each partition to have its own rate of evolution. Second, phylogenomic analysis was repeated using the Bayesian method (aamodelpr=mixed) in MrBayes (v3.2.6) (Huelsenbeck and Ronquist 2001; Ronquist and Huelsenbeck 2003; Ronquist et al. 2012). The Markov chain Monte Carlo (MCMC) analysis was run using 1 chain for 400,000 generations. A consensus tree is presented along with posterior probabilities. Third, unrooted ML trees of each of the single-copy orthologs were produced with IQ-TREE 3 (1,000 bootstrap replicates) and then the supertree method of weighted Accurate Species TRee ALgorithm (wASTRAL) was implemented to infer a species tree from the 2,335 individual gene trees. wASTRAL is an improvement on the popular tool ASTRAL (Mirarab et al. 2014; Zhang et al. 2018) that, rather than treating all input gene trees equally, weights each quartet by its statistical support (Zhang and Mirarab 2022). All trees were rendered in iToL (v6) (Letunic and Bork 2024).

Divergence time estimation was performed using MCMCTree (v4.10.6), part of the PAML package (Yang and Rannala 2006; Rannala and Yang 2007; dos Reis and Yang 2011). To prepare the molecular dataset for tree calibration, we used the 2,335 single-copy orthologs identified from the previous OrthoFinder analysis as an initial baseline. For each orthogroup, a phylogenetic tree was constructed using IQ-TREE 3, enabling the ModelFinder option to select the best-fit substitution model and activating write per-site rates to calculate per-site rates for subsequent clock-likeness analysis. Orthologs producing trees with average bootstrap support values below 70 were excluded from subsequent analysis. To assess the clock-likeness of each remaining ortholog, we calculated the Degree of Violation of the Molecular Clock (DVMC) (Liu et al. 2017) for all trees using PhyKIT. DVMC values were plotted to visualise the distribution of genes with respect to their deviation from clock-like behaviour. Based on this distribution, we applied a cutoff of 0.8 for DVMC, which represented a balance between minimising ortholog loss and eliminating orthologs with unacceptable molecular clock properties. These filtering steps reduced the dataset from 2,335 to 1,722 orthologs.

Fossil calibration data were obtained from the Paleobiology Database (https://paleobiodb.org/; accessed March 2025). The root of the tree was calibrated using the oldest known fossil of a sea anemone from the early Silurian period (Barroso et al. 2025), representing the earliest divergence within Anthozoa (Table S5). Before running MCMCTree, we estimated the prior parameters for rate variation: sigma2_gamma and rgene_gamma. For sigma2_gamma, which models the variance in substitution rates across lineages, we used a Gamma distribution with parameters Γ (2, 10). This setting allows for high rate heterogeneity, which is appropriate for deep phylogenies where strict molecular clock assumptions may not hold. For rgene_gamma, which specifies the prior on the overall evolutionary rate, we estimated the mean rate (μ) using the tree height obtained from a maximum likelihood tree built with the filtered gene set (1,722 genes) and the selected root age (from selected priors). The scale parameter (β) of the Gamma distribution was calculated using the relation β = α / μ. We plotted the resulting prior distributions using different α values to ensure the prior was not unrealistically narrow or overly diffuse, given the estimated depth and the calibration points used in the tree.

With the molecular data, fossil calibrations in place, and prior distributions defined, we first ran MCMCTree to sample from the prior alone. This step allowed us to verify that the specified priors produced a reasonable distribution of divergence times, independent of the molecular data. Subsequently, we ran MCMCTree to calculate the gradient and Hessian matrices. This step involves estimating the maximum likelihood branch lengths (b), as well as the gradient (g) and Hessian (H) of the log-likelihood function, evaluated at the maximum likelihood estimates (MLEs). These matrices are required to approximate the likelihood function during MCMC sampling efficiently. For the amino acid substitution model, we used the WAG (Whelan and Goldman) model, which is appropriate for protein sequences and commonly used in deep metazoan phylogenies. Once the gradient and Hessian were computed, we ran the final MCMC analysis using prior information with the approximate likelihood to sample divergence times. The resulting time-calibrated phylogenetic tree was visualised using the R package MCMCtreeR (v1.1) (Puttick 2019).

## Acknowledgments

We thank the High-Throughput Genomics Core at the Biodiversity Research Center, Academia Sinica for sequencing support. We thank Tzu-Wen Tai, Hui-Ju Lin, and the Flow Cytometry Cell Sorting Core at the Institute of Biomedical Sciences, Academia Sinica for technical assistance with cell sorting. We are grateful to the Aquatic Symbiosis Genomics Project of the Tree of Life Programme at the Wellcome Sanger Institute for making high-quality genomes openly accessible to the community. We gratefully acknowledge the NMMBA for providing the corallimorpharian specimens utilized in this study. We thank the members of the Symbiosis Genomics & Evolution Lab for their assistance and support. Photo acknowledgments for Fig. 1a: *D. lineata* by Oscar Dove; *P. pennacea* by Neil DeMaster; *R. yuma* by craigjhowe; *P. verrucosa* by Damien Brouste; and *A. millepora* by brigitteleroy (iNaturalist; licensed under CC BY-NC 4.0).

## Supplementary material

Supplementary material is available at *Molecular Biology and Evolution* online.

## Author contributions

T.D.L., S.S., M.-F.L., and Y.-J.L. conceived the project. Y.-L.C. and M.-F.L. collected and prepared *R. yuma* samples. Y.-L.C., K.S., and S.S. collected and prepared *G. fascicularis* samples. L.-J.K., Y.-L.C., J.-Y.L., K.-X.T., and M.-Y.J.L. performed genome sequencing. T.D.L., V.M.P.-G., Y.Y., and Y.-J.L. assembled and annotated the *R. yuma* genome. T.D.L. and V.M.P.-G. curated and annotated publicly available genomes. T.D.L., V.M.P.-G., Y.Y., and Y.-J.L. carried out synteny analyses. L.-J.K., Y.-L.C., and Y.-H.C. performed single-cell preparation and library construction. T.S., V.M.P.-G., and Y.-J.L. analyzed single-cell data. T.D.L., T.S., V.M.P.-G., and Y.Y. carried out comparative genomic analyses. T.D.L. and V.M.P.-G. conducted phylogenetic analyses. T.S. and V.M.P.-G. performed transcriptomic analyses. T.D.L., T.S., V.M.P.-G., and Y.-J.L. interpreted the data, prepared figures, and wrote the manuscript. Y.Y., Y.-L.C., D.J.M., S.S., and M.-F.L. discussed the results and edited the manuscript.

## Data availability

Genome sequencing and RNA sequencing datasets of the corallimorpharian *R. yuma* are accessible through NCBI BioProject (PRJNA1532872). Single-cell RNA sequencing data for *R. yuma* and *G. fascicularis* polyps are available through NCBI GEO (GSEXXXXXX). Gene models, including coding sequences, protein sequences, and gene annotations for anthozoan genomes, together with newly generated *G. fascicularis* transcriptome and single-cell data, are deposited on Zenodo (https://doi.org/10.5281/zenodo.22929641). Code for analyses in this study is available on GitHub (https://github.com/symgenoevolab/ricordea_genome).

## References

Ahlmann-Eltze C, Huber W. 2021. glmGamPoi: fitting Gamma-Poisson generalized linear models on single cell count data. Bioinformatics 36:5701–5702.

Alexa A, Rahnenführer J, Lengauer T. 2006. Improved scoring of functional groups from gene expression data by decorrelating GO graph structure. Bioinformatics 22:1600– 1607.

Almeida-Silva F, Zhao T, Ullrich KK, Schranz ME, Van de Peer Y. 2023. syntenet: an R/Bioconductor package for the inference and analysis of synteny networks. Bioinformatics 39:btac806.

Arendt D, Musser JM, Baker CVH, Bergman A, Cepko C, Erwin DH, Pavlicev M, Schlosser G, Widder S, Laubichler MD, et al. 2016. The origin and evolution of cell types. Nat. Rev. Genet. 17:744–757.

Balthasar U, Cusack M. 2015. Aragonite-calcite seas—Quantifying the gray area. Geology 43:99–102.

Barnett DW, Garrison EK, Quinlan AR, Strömberg MP, Marth GT. 2011. BamTools: a C++ API and toolkit for analyzing and managing BAM files. Bioinformatics 27:1691–1692.

Barrera-Redondo J, Lotharukpong JS, Drost H-G, Coelho SM. 2023. Uncovering gene-family founder events during major evolutionary transitions in animals, plants and fungi using GenEra. Genome Biol. 24:54.

Barroso FRG, Viana MSS, Agostinho S, Daly M, Fairchild TR, Marques AC, Pacheco MLAF. 2025. Insights into the lifestyle and preservation of Arenactinia ipuensis n. gen. et n. sp. (Anthozoa, Actiniaria) from the Early Silurian (Ipu Formation, Parnaíba Basin, Brazil). Earth History and Biodiversity 3:100017.

Bonacolta A, Snyder G, Karp R, Yeager E, Wen A, Dennison C, Nonell J, Traylor-Knowles N, Baker A, Campo J del. 2024. A single-cell atlas of coral bleaching. Research Square [Internet]. Available from: https://www.researchsquare.com/article/rs-5397639/latest.pdf

Brůna T, Hoff KJ, Lomsadze A, Stanke M, Borodovsky M. 2021. BRAKER2: automatic eukaryotic genome annotation with GeneMark-EP+ and AUGUSTUS supported by a protein database. NAR Genom Bioinform 3:lqaa108.

Buchfink B, Xie C, Huson DH. 2015. Fast and sensitive protein alignment using DIAMOND. Nat. Methods 12:59–60.

Budd AF, Stolarski J. 2011. Corallite wall and septal microstructure in scleractinian reef corals: comparison of molecular clades within the family Faviidae. J. Morphol. 272:66– 88.

Bushmanova E, Antipov D, Lapidus A, Suvorov V, Prjibelski AD. 2016. rnaQUAST: a quality assessment tool for de novo transcriptome assemblies. Bioinformatics 32:2210–2212.

Camacho C, Coulouris G, Avagyan V, Ma N, Papadopoulos J, Bealer K, Madden TL. 2009. BLAST+: architecture and applications. BMC Bioinformatics 10:421.

Campoy AN, Addamo AM, Machordom A, Meade A, Rivadeneira MM, Hernández CE, Venditti C. 2020. The origin and correlated evolution of symbiosis and coloniality in scleractinian corals. Front. Mar. Sci. 7:461.

Campoy AN, Rivadeneira MM, Hernández CE, Meade A, Venditti C. 2023. Deep-sea origin and depth colonization associated with phenotypic innovations in scleractinian corals. Nat. Commun. 14:7458.

Cantalapiedra CP, Hernández-Plaza A, Letunic I, Bork P, Huerta-Cepas J. 2021. EggNOG-mapper v2: Functional annotation, orthology assignments, and domain prediction at the metagenomic scale. Mol. Biol. Evol. 38:5825–5829.

Capella-Gutiérrez S, Silla-Martínez JM, Gabaldón T. 2009. trimAl: a tool for automated alignment trimming in large-scale phylogenetic analyses. Bioinformatics 25:1972–1973.

Chen CA, Odorico DM, ten Lohuis M, Veron JE, Miller DJ. 1995. Systematic relationships within the Anthozoa (Cnidaria: Anthozoa) using the 5’-end of the 28S rDNA. Mol. Phylogenet. Evol. 4:175–183.

Chen CA, Wallace CC, Wolstenholme J. 2002. Analysis of the mitochondrial 12S rRNA gene supports a two-clade hypothesis of the evolutionary history of scleractinian corals. Mol. Phylogenet. Evol. 23:137–149.

Cheng H, Concepcion GT, Feng X, Zhang H, Li H. 2021. Haplotype-resolved de novo assembly using phased assembly graphs with hifiasm. Nat. Methods 18:170–175.

Chen S. 2023. Ultrafast one-pass FASTQ data preprocessing, quality control, and deduplication using fastp. Imeta 2:e107.

Chen S, Zhou Y, Chen Y, Gu J. 2018. fastp: an ultra-fast all-in-one FASTQ preprocessor. Bioinformatics 34:i884–i890.

Chen Z, Baeza JA, Chen C, Gonzalez MT, González VL, Greve C, Kocot KM, Arbizu PM, Moles J, Schell T, et al. 2025. A genome-based phylogeny for Mollusca is concordant with fossils and morphology. Science 387:1001–1007.

Chernomor O, von Haeseler A, Minh BQ. 2016. Terrace aware data structure for phylogenomic inference from supermatrices. Syst. Biol. 65:997–1008.

Copley RR. 2025. Sponges, ctenophores, and the statistical significance of syntenies. Mol. Biol. Evol. 42:msaf321.

Daly M, Fautin DG, Cappola VA. 2003. Systematics of the Hexacorallia (Cnidaria: Anthozoa). Zool. J. Linn. Soc. 139:419–437.

Doney SC, Fabry VJ, Feely RA, Kleypas JA. 2009. Ocean Acidification: The Other CO 2 Problem. Annu. Rev. Mar. Sci. 1:169–192.

Drake JL, Mass T, Haramaty L, Zelzion E, Bhattacharya D, Falkowski PG. 2013. Proteomic analysis of skeletal organic matrix from the stony coral Stylophora pistillata. Proc. Natl. Acad. Sci. U. S. A. 110:3788–3793.

Drake JL, Mass T, Stolarski J, Von Euw S, van de Schootbrugge B, Falkowski PG. 2020. How corals made rocks through the ages. Glob. Chang. Biol. 26:31–53.

Drost H-G, Gabel A, Liu J, Quint M, Grosse I. 2018. myTAI: evolutionary transcriptomics with R. Bioinformatics 34:1589–1590.

Durand NC, Robinson JT, Shamim MS, Machol I, Mesirov JP, Lander ES, Aiden EL. 2016. Juicebox Provides a Visualization System for Hi-C Contact Maps with Unlimited Zoom. Cell Syst 3:99–101.

Emms DM, Kelly S. 2019. OrthoFinder: phylogenetic orthology inference for comparative genomics. Genome Biol. 20:238.

Ewels P, Magnusson M, Lundin S, Käller M. 2016. MultiQC: summarize analysis results for multiple tools and samples in a single report. Bioinformatics 32:3047–3048.

Fautin DG. 2016. Catalog to families, genera, and species of orders Actiniaria and Corallimorpharia (Cnidaria: Anthozoa). Zootaxa 4145:1–449.

Fine M, Tchernov D. 2007. Scleractinian coral species survive and recover from decalcification. Science 315:1811.

Fletcher C, Pereira da Conceicoa L, Natural History Museum Genome Acquisition Lab, Darwin Tree of Life Barcoding collective, Wellcome Sanger Institute Tree of Life programme, Wellcome Sanger Institute Scientific Operations: DNA Pipelines collective, Tree of Life Core Informatics collective, Darwin Tree of Life Consortium. 2023. The genome sequence of the starlet sea anemone, *Nematostella* vectensis (Stephenson, 1935). Wellcome Open Res. 8:79.

Flynn JM, Hubley R, Goubert C, Rosen J, Clark AG, Feschotte C, Smit AF. 2020. RepeatModeler2 for automated genomic discovery of transposable element families. Proc. Natl. Acad. Sci. U. S. A. 117:9451–9457.

Formenti G, Abueg L, Brajuka A, Brajuka N, Gallardo-Alba C, Giani A, Fedrigo O, Jarvis ED. 2022. Gfastats: conversion, evaluation and manipulation of genome sequences using assembly graphs. Bioinformatics 38:4214–4216.

Fukami H, Chen CA, Budd AF, Collins A, Wallace C, Chuang Y-Y, Chen C, Dai C-F, Iwao K, Sheppard C, et al. 2008. Mitochondrial and nuclear genes suggest that stony corals are monophyletic but most families of stony corals are not (Order Scleractinia, Class Anthozoa, Phylum Cnidaria). PLoS One 3:e3222.

Fuller ZL, Mocellin VJL, Morris LA, Cantin N, Shepherd J, Sarre L, Peng J, Liao Y, Pickrell J, Andolfatto P, et al. 2020. Population genetics of the coral Acropora millepora: Toward genomic prediction of bleaching. Science 369:eaba4674.

Gabriel L, Brůna T, Hoff KJ, Ebel M, Lomsadze A, Borodovsky M, Stanke M. 2024. BRAKER3: Fully automated genome annotation using RNA-seq and protein evidence with GeneMark-ETP, AUGUSTUS, and TSEBRA. Genome Res. 34:769–777.

Gallego-Garcia J, Hays D, Tongboonkua P, Minich JJ, Hilgers L, Michael TP, Hiller M, Zhang C, Orti G, Arcila D, et al. 2026. Whole-genome phylogenomics and synteny resolve a single origin of body-plan asymmetry in flatfishes. bioRxiv [Internet]:2026.04.07.717116. Available from: https://www.biorxiv.org/content/10.64898/2026.04.07.717116v2.abstract

García-Castro H, Kenny NJ, Iglesias M, Álvarez-Campos P, Mason V, Elek A, Schönauer A, Sleight VA, Neiro J, Aboobaker A, et al. 2021. ACME dissociation: a versatile cell fixation-dissociation method for single-cell transcriptomics. Genome Biol. 22:89.

Gattuso J. 1998. Effect of calcium carbonate saturation of seawater on coral calcification. Glob. Planet. Change 18:37–46.

Gilbert D. 2016. Gene-omes built from mRNA seq not genome DNA. F1000Research [Internet] 5. Available from: 10.7490/f1000research.1112594.1

Grabherr MG, Haas BJ, Yassour M, Levin JZ, Thompson DA, Amit I, Adiconis X, Fan L, Raychowdhury R, Zeng Q, et al. 2011. Full-length transcriptome assembly from RNA-Seq data without a reference genome. Nat. Biotechnol. 29:644–652.

Grau-Bové X, Montes-Espuña L, Księżopolska E, Najle SR, Mass T, Levy S, Sebé-Pedrós A. 2026. Cell-resolved transcriptional responses during heat-induced coral bleaching and recovery. bioRxiv [Internet]:2026.07.18.739195. Available from: https://www.biorxiv.org/content/10.64898/2026.07.18.739195v1.abstract

Grossman EL, Joachimski MM. 2022. Ocean temperatures through the Phanerozoic reassessed. Sci. Rep. 12:8938.

Guan D, McCarthy SA, Wood J, Howe K, Wang Y, Durbin R. 2020. Identifying and removing haplotypic duplication in primary genome assemblies. Bioinformatics 36:2896–2898.

Guinotte JM, Fabry VJ. 2008. Ocean Acidification and Its Potential Effects on Marine Ecosystems. Annals of the New York Academy of Sciences 1134:320–342.

Guinotte JM, Orr J, Cairns S, Freiwald A, Morgan L, George R. 2006. Will human-induced changes in seawater chemistry alter the distribution of deep-sea scleractinian corals? Front. Ecol. Environ. 4:141–146.

Haas BJ, Papanicolaou A, Yassour M, Grabherr M, Blood PD, Bowden J, Couger MB, Eccles D, Li B, Lieber M, et al. 2013. De novo transcript sequence reconstruction from RNA-seq using the Trinity platform for reference generation and analysis. Nat. Protoc. 8:1494–1512.

Han T, Li Y, Zhao H, Chen J, He C, Lu Z. 2025. In-depth single-cell transcriptomic exploration of the regenerative dynamics in stony coral. *Commun*. Biol. 8:652.

Hao Y, Stuart T, Kowalski MH, Choudhary S, Hoffman P, Hartman A, Srivastava A, Molla G, Madad S, Fernandez-Granda C, et al. 2024. Dictionary learning for integrative, multimodal and scalable single-cell analysis. Nat. Biotechnol. 42:293–304.

Hao Z, Lv D, Ge Y, Shi J, Weijers D, Yu G, Chen J. 2020. RIdeogram: drawing SVG graphics to visualize and map genome-wide data on the idiograms. PeerJ Comput Sci 6:e251.

den Hartog JC. 1980. Caribbean shallow water Corallimorpharia. Zoologische Verhandelingen 176:1–83.

Henschel R, Lieber M, Wu L-S, Nista PM, Haas BJ, LeDuc RD. 2012. Trinity RNA-Seq assembler performance optimization. In: Proceedings of the 1st Conference of the Extreme Science and Engineering Discovery Environment: Bridging from the eXtreme to the campus and beyond. New York, NY, USA: ACM. p. 1–8.

Hoang DT, Chernomor O, von Haeseler A, Minh BQ, Vinh LS. 2018. UFBoot2: Improving the Ultrafast Bootstrap Approximation. Mol. Biol. Evol. 35:518–522.

Hoegh-Guldberg O, Mumby PJ, Hooten AJ, Steneck RS, Greenfield P, Gomez E, Harvell CD, Sale PF, Edwards AJ, Caldeira K, et al. 2007. Coral reefs under rapid climate change and ocean acidification. Science 318:1737–1742.

Hoff KJ, Lange S, Lomsadze A, Borodovsky M, Stanke M. 2016. BRAKER1: Unsupervised RNA-Seq-Based Genome Annotation with GeneMark-ET and AUGUSTUS. Bioinformatics 32:767–769.

Hoff KJ, Lomsadze A, Borodovsky M, Stanke M. 2019. Whole-Genome Annotation with BRAKER. Methods Mol. Biol. 1962:65–95.

Hohenauer T, Moore AW. 2012. The Prdm family: expanding roles in stem cells and development. Development 139:2267–2282.

Huelsenbeck J, Ronquist F. 2001. MRBAYES: Bayesian inference of phylogenetic trees. Bioinformatics 17:754–755.

Huerta-Cepas J, Szklarczyk D, Heller D, Hernández-Plaza A, Forslund SK, Cook H, Mende DR, Letunic I, Rattei T, Jensen LJ, et al. 2019. eggNOG 5.0: a hierarchical, functionally and phylogenetically annotated orthology resource based on 5090 organisms and 2502 viruses. Nucleic Acids Res. 47:D309–D314.

Hu M, Zheng X, Fan C-M, Zheng Y. 2020. Lineage dynamics of the endosymbiotic cell type in the soft coral Xenia. Nature 582:534–538.

Johnston IS. 1980. The Ultrastructure of Skeletogenesis in Hermatypic Corals. In: International Review of Cytology. International Review of Cytology. Elsevier. p. 171– 214.

Kalyaanamoorthy S, Minh BQ, Wong TKF, von Haeseler A, Jermiin LS. 2017. ModelFinder: fast model selection for accurate phylogenetic estimates. Nat. Methods 14:587–589.

Katoh K, Misawa K, Kuma K-I, Miyata T. 2002. MAFFT: a novel method for rapid multiple sequence alignment based on fast Fourier transform. Nucleic Acids Res. 30:3059–3066.

Katoh K, Standley DM. 2013. MAFFT multiple sequence alignment software version 7: improvements in performance and usability. Mol. Biol. Evol. 30:772–780.

Kayal E, Bentlage B, Sabrina Pankey M, Ohdera AH, Medina M, Plachetzki DC, Collins AG, Ryan JF. 2018. Phylogenomics provides a robust topology of the major cnidarian lineages and insights on the origins of key organismal traits. BMC Evol. Biol. [Internet] 18. Available from: 10.1186/s12862-018-1142-0

Kayal E, Roure B, Philippe H, Collins AG, Lavrov DV. 2013. Cnidarian phylogenetic relationships as revealed by mitogenomics. BMC Evol. Biol. 13:5.

Khalturin K, Shinzato C, Khalturina M, Hamada M, Fujie M, Koyanagi R, Kanda M, Goto H, Anton-Erxleben F, Toyokawa M, et al. 2019. Medusozoan genomes inform the evolution of the jellyfish body plan. *Nat*. Ecol. Evol. 3:811–822.

Kim D, Paggi JM, Park C, Bennett C, Salzberg SL. 2019. Graph-based genome alignment and genotyping with HISAT2 and HISAT-genotype. Nat. Biotechnol. 37:907–915.

Kitahara MV, Cairns SD, Stolarski J, Blair D, Miller DJ. 2010. A comprehensive phylogenetic analysis of the Scleractinia (Cnidaria, Anthozoa) based on mitochondrial CO1 sequence data. PLoS One 5:e11490.

Kitahara MV, Lin M-F, Forêt S, Huttley G, Miller DJ, Chen CA. 2014. The “naked coral” hypothesis revisited--evidence for and against scleractinian monophyly. PLoS One 9:e94774.

Kuguru BL, Mgaya YD, ◻hman MC, Wagner GM. 2004. The reef environment and competitive success in the Corallimorpharia. Mar. Biol. 145:875–884.

Kuguru B, Winters G, Beer S, Santos SR, Chadwick NE. 2007. Adaptation strategies of the corallimorpharian Rhodactis rhodostoma to irradiance and temperature. Mar. Biol. 151:1287–1298.

Kulkarni SS, Klementz BC, Ballesteros JA, Abshire KM, Cunha TJ, Hassan MK, Laumer EM, De Medeiros BAS, Neu SM, Pankey MS, et al. 2026. No silver bullet: Patterns of macrosynteny recapitulate systemic conflicts in the higher-level relationships of the arachnids. bioRxiv [Internet]:2026.06.22.733561. Available from: https://www.biorxiv.org/content/10.64898/2026.06.22.733561v1.abstract

Kuznetsov D, Tegenfeldt F, Manni M, Seppey M, Berkeley M, Kriventseva EV, Zdobnov EM. 2023. OrthoDB v11: annotation of orthologs in the widest sampling of organismal diversity. Nucleic Acids Res. 51:D445–D451.

LaJeunesse. 2002. Diversity and community structure of symbiotic dinoflagellates from Caribbean coral reefs. Mar. Biol. 141:387–400.

LaJeunesse TC, Bhagooli R, Hidaka M, deVantier L, Done T, Schmidt GW, Fitt WK, Hoegh-Guldberg O. 2004. Closely related Symbiodinium spp. differ in relative dominance in coral reef host communities across environmental, latitudinal and biogeographic gradients. Mar. Ecol. Prog. Ser. 284:147–161.

Leinonen R, Sugawara H, Shumway M, International Nucleotide Sequence Database Collaboration. 2011. The sequence read archive. Nucleic Acids Res. 39:D19–D21.

Le Tissier MDA. 1990. The ultrastructure of the skeleton and skeletogenic tissues of the temperate coral Caryophyllia smithii. J. Mar. Biol. Assoc. U. K. 70:295–310.

Letunic I, Bork P. 2024. Interactive Tree of Life (iTOL) v6: recent updates to the phylogenetic tree display and annotation tool. Nucleic Acids Res. 52:W78–W82.

Levy S, Elek A, Grau-Bové X, Menéndez-Bravo S, Iglesias M, Tanay A, Mass T, Sebé-Pedrós A. 2021. A stony coral cell atlas illuminates the molecular and cellular basis of coral symbiosis, calcification, and immunity. Cell 184:2973–2987.e18.

Levy S, Grau-Bové X, Kim IV, Najle SR, Księżopolska E, Elek A, Montes-Espuña L, Montgomery SA, Mass T, Sebé-Pedrós A. 2025. The evolution of facultative symbiosis in stony corals. Nature 648:368–376.

Lewin TD, Liao IJ-Y, Chen M-E, Bishop JDD, Holland PWH, Luo Y-J. 2025. Fusion, fission, and scrambling of the bilaterian genome in Bryozoa. Genome Res. 35:78–92.

Lewin TD, Liao IJ-Y, Luo Y-J. 2024. Annelid comparative genomics and the evolution of massive lineage-specific genome rearrangement in bilaterians. Mol. Biol. Evol. 41:msae172.

Lewin TD, Liao IJ-Y, Luo Y-J. 2025. Conservation of bilaterian genome structure is the exception, not the rule. Genome Biol. 26:247.

Lewin TD, Sakagami T, Kao L-J, Liao IJ-Y, Chiu Y-L, Piñon-Gonzalez VM, Li J-Y, Tin K-X, Lu M-YJ, Endo K, et al. 2025. Phoronid genome supports a monophyletic Lophophorata. Curr. Biol. 35:5633–5645.e7.

Lewin TD, Sakagami T, Shimizu K, Kao L-J, Chiu Y-L, Liao IJ-Y, Chen M-E, Hisata K, Endo K, Satoh N, et al. 2026. Brachiopod genome unveils the evolution of BMP signalling in bilaterian body patterning. Nat. Commun. 17:3856.

Li H, Handsaker B, Wysoker A, Fennell T, Ruan J, Homer N, Marth G, Abecasis G, Durbin R. 2009. The Sequence Alignment/Map format and SAMtools. Bioinformatics 25:2078– 2079.

Lin M-F, Chen CA, Miller DJ. 2013. Asexual reproduction by marginal budding in the tropical corallimorpharian, Ricordea yuma (Corallimorpharia; Ricordeidae). Galaxea (Tokyo) 15:41–42.

Lin MF, Chou WH, Kitahara MV, Chen CLA, Miller DJ, Forêt S. 2016. Corallimorpharians are not “naked corals”: insights into relationships between Scleractinia and Corallimorpharia from phylogenomic analyses. PeerJ 4:e2463.

Lin M-F, Kitahara MV, Luo H, Tracey D, Geller J, Fukami H, Miller DJ, Chen CA. 2014. Mitochondrial genome rearrangements in the scleractinia/corallimorpharia complex: implications for coral phylogeny. Genome Biol. Evol. 6:1086–1095.

Lin Z, Akin H, Rao R, Hie B, Zhu Z, Lu W, Smetanin N, Verkuil R, Kabeli O, Shmueli Y, et al. 2023. Evolutionary-scale prediction of atomic-level protein structure with a language model. Science 379:1123–1130.

Liu L, Zhang J, Rheindt FE, Lei F, Qu Y, Wang Y, Zhang Y, Sullivan C, Nie W, Wang J, et al. 2017. Genomic evidence reveals a radiation of placental mammals uninterrupted by the KPg boundary. Proc. Natl. Acad. Sci. U. S. A. 114:E7282–E7290.

Li Y, Liu H, Steenwyk JL, LaBella AL, Harrison M-C, Groenewald M, Zhou X, Shen X-X, Zhao T, Hittinger CT, et al. 2022. Contrasting modes of macro and microsynteny evolution in a eukaryotic subphylum. Curr. Biol. 32:5335–5343.e4.

Llorens-Giralt P, Camilleri-Robles C, Zuber L, Khadka J, Marti-Marimon M, Crespo-Pera A, Serras F, Gambetta MC, Marti-Renom MA, Corominas M. 2026. 3D genome organization in tissue regeneration involves long-range chromatin loops. Sci. Adv. 12:eaea8281.

Lomsadze A, Burns PD, Borodovsky M. 2014. Integration of mapped RNA-Seq reads into automatic training of eukaryotic gene finding algorithm. Nucleic Acids Res. 42:e119.

Lovell JT, Sreedasyam A, Schranz ME, Wilson M, Carlson JW, Harkess A, Emms D, Goodstein DM, Schmutz J. 2022. GENESPACE tracks regions of interest and gene copy number variation across multiple genomes. Elife 11:e78526.

Love MI, Huber W, Anders S. 2014. Moderated estimation of fold change and dispersion for RNA-seq data with DESeq2. Genome Biol. 15:550.

Lu C-Y, Chang Y-P, Piñon-Gonzalez VM, Lewin TD, Shih K-N, Chuang P-S, Wada N, Yu S-P, Yu Z-R, Chen Y-H, et al. 2026. Endozoicomonas acroporae enhances coral thermal resilience through host-microbe coordination. ISME J. 20:wrag173.

Mason B, Cooke I, Moya A, Augustin R, Lin M-F, Satoh N, Bosch TCG, Bourne DG, Hayward DC, Andrade N, et al. 2021. AmAMP1 from Acropora millepora and damicornin define a family of coral-specific antimicrobial peptides related to the Shk toxins of sea anemones. Dev. Comp. Immunol. 114:103866.

McFadden CS, Quattrini AM, Brugler MR, Cowman PF, Dueñas LF, Kitahara MV, Paz-García DA, Reimer JD, Rodríguez E. 2021. Phylogenomics, origin, and diversification of anthozoans (phylum Cnidaria). Syst. Biol. 70:635–647.

McKenna V, Archibald JM, Beinart R, Dawson MN, Hentschel U, Keeling PJ, Lopez JV, Martín-Durán JM, Petersen JM, Sigwart JD, et al. 2024. The Aquatic Symbiosis Genomics Project: probing the evolution of symbiosis across the Tree of Life. Wellcome Open Res. 6:254.

Medina M, Collins AG, Takaoka TL, Kuehl JV, Boore JL. 2006. Naked corals: skeleton loss in Scleractinia. Proc. Natl. Acad. Sci. U. S. A. 103:9096–9100.

Minh BQ, Schmidt HA, Chernomor O, Schrempf D, Woodhams MD, von Haeseler A, Lanfear R. 2020. IQ-TREE 2: New Models and Efficient Methods for Phylogenetic Inference in the Genomic Era. Mol. Biol. Evol. 37:1530–1534.

Mirarab S, Reaz R, Bayzid MS, Zimmermann T, Swenson MS, Warnow T. 2014. ASTRAL: genome-scale coalescent-based species tree estimation. Bioinformatics 30:i541–i548.

Misteli T. 2020. The self-organizing genome: Principles of genome architecture and function. Cell 183:28–45.

Mollica NR, Guo W, Cohen AL, Huang K-F, Foster GL, Donald HK, Solow AR. 2018. Ocean acidification affects coral growth by reducing skeletal density. Proc. Natl. Acad. Sci. U. S. A. 115:1754–1759.

Moriya Y, Itoh M, Okuda S, Yoshizawa AC, Kanehisa M. 2007. KAAS: an automatic genome annotation and pathway reconstruction server. Nucleic Acids Res. 35:W182–W185.

Moya A, Tambutté S, Bertucci A, Tambutté E, Lotto S, Vullo D, Supuran CT, Allemand D, Zoccola D. 2008. Carbonic anhydrase in the scleractinian coral Stylophora pistillata: characterization, localization, and role in biomineralization. J. Biol. Chem. 283:25475– 25484.

Mummadisetti MP, Drake JL, Falkowski PG. 2021. The spatial network of skeletal proteins in a stony coral. J. R. Soc. Interface 18:20200859.

Parey E, Louis A, Montfort J, Bouchez O, Roques C, Iampietro C, Lluch J, Castinel A, Donnadieu C, Desvignes T, et al. 2023. Genome structures resolve the early diversification of teleost fishes. Science 379:572–575.

Patro R, Duggal G, Love MI, Irizarry RA, Kingsford C. 2017. Salmon provides fast and bias-aware quantification of transcript expression. Nat. Methods 14:417–419.

Pearce JD, Simmonds SE, Mahmoudabadi G, Krishnan L, Palla G, Istrate A-M, Tarashansky A, Nelson B, Valenzuela O, Li D, et al. 2026. TranscriptFormer: A generative cell atlas across 1.5 billion years of evolution. Science:eaec8514.

Plessy C, Mansfield MJ, Bliznina A, Masunaga A, West C, Tan Y, Liu AW, Grašič J, Del Río Pisula MS, Sánchez-Serna G, et al. 2024. Extreme genome scrambling in marine planktonic Oikopleura dioica cryptic species. Genome Res. 34:426–440.

Porter SM. 2010. Calcite and aragonite seas and the de novo acquisition of carbonate skeletons. Geobiology 8:256–277.

Puntin G, Craggs J, Hayden R, Engelhardt KE, McIlroy S, Sweet M, Baker DM, Ziegler M. 2022. The reef-building coral Galaxea fascicularis: a new model system for coral symbiosis research. Coral Reefs 42:239–252.

Puttick MN. 2019. MCMCtreeR: functions to prepare MCMCtree analyses and visualize posterior ages on trees. Bioinformatics 35:5321–5322.

Quattrini AM, Faircloth BC, Dueñas LF, Bridge TCL, Brugler MR, Calixto-Botía IF, DeLeo DM, Forêt S, Herrera S, Lee SMY, et al. 2018. Universal target-enrichment baits for anthozoan (Cnidaria) phylogenomics: New approaches to long-standing problems. Mol. Ecol. Resour. 18:281–295.

Quattrini AM, Rodríguez E, Faircloth BC, Cowman PF, Brugler MR, Farfan GA, Hellberg ME, Kitahara MV, Morrison CL, Paz-García DA, et al. 2020. Palaeoclimate ocean conditions shaped the evolution of corals and their skeletons through deep time. *Nat*. Ecol. Evol. 4:1531–1538.

Quek ZBR, Jain SS, Richards ZT, Arrigoni R, Benzoni F, Hoeksema BW, Carvajal JI, Wilson NG, Baird AH, Kitahara MV, et al. 2023. A hybrid-capture approach to reconstruct the phylogeny of Scleractinia (Cnidaria: Hexacorallia). Mol. Phylogenet. Evol. 186:107867.

Ramos-Silva P, Kaandorp J, Huisman L, Marie B, Zanella-Cléon I, Guichard N, Miller DJ, Marin F. 2013. The skeletal proteome of the coral Acropora millepora: the evolution of calcification by co-option and domain shuffling. Mol. Biol. Evol. 30:2099–2112.

Rannala B, Yang Z. 2007. Inferring speciation times under an episodic molecular clock. Syst. Biol. 56:453–466.

R Core Team. 2024. R: A language and environment for statistical computing. Vienna: R foundation for statistical computing. *Vienna: R foundation for statistical computing* [Internet]. Available from: https://www.R-project.org/

dos Reis M, Yang Z. 2011. Approximate likelihood calculation on a phylogeny for Bayesian estimation of divergence times. Mol. Biol. Evol. 28:2161–2172.

Ridgwell A, Zeebe R. 2005. The role of the global carbonate cycle in the regulation and evolution of the Earth system. Earth Planet. Sci. Lett. 234:299–315.

Rokas A, Holland PW. 2000. Rare genomic changes as a tool for phylogenetics. Trends Ecol. Evol. 15:454–459.

Romano SL, Cairns SD. 2000. Molecular phylogenetic hypotheses for the evolution of scleractinian corals. Bulletin of Marine Science 67:1043–1068.

Romano SL, Palumbi SR. 1996. Evolution of Scleractinian Corals Inferred from Molecular Systematics. Science 271:640–642.

Ronquist F, Huelsenbeck J. 2003. MrBayes 3: Bayesian phylogenetic inference under mixed models. Bioinformatics 19:1572–1574.

Ronquist F, Teslenko M, P. VDM, Ayres DL, Darling A, Höhna S, Huelsenbeck J. 2012. MrBayes 3.2: efficient Bayesian phylogenetic inference and model choice across a large model space. Systematic biology 61:539–542.

Schafran P, Hauser DA, Nelson JM, Xu X, Mueller LA, Kulshrestha S, Smalley I, de Vries S, Irisarri I, de Vries J, et al. 2025. Pan-phylum genomes of hornworts reveal conserved autosomes but dynamic accessory and sex chromosomes. Nat. Plants 11:49–62.

Schultz DT, Haddock SHD, Bredeson JV, Green RE, Simakov O, Rokhsar DS. 2023. Ancient gene linkages support ctenophores as sister to other animals. Nature 618:110– 117.

Sebé-Pedrós A, Saudemont B, Chomsky E, Plessier F, Mailhé M-P, Renno J, Loe-Mie Y, Lifshitz A, Mukamel Z, Schmutz S, et al. 2018. Cnidarian cell type diversity and regulation revealed by whole-organism single-cell RNA-seq. Cell 173:1520–1534.e20.

Shikina S, Cheng Y-C, Lin T-C, Chang Y-E, Tsai P-H, Chen E, Chang Y-C, Leung Y-Y, Yu Y-C, Chiu Y-L. 2024. Coral mariculture using abandoned abalone farming ponds in northeastern Taiwan. Aquaculture 592:Not Available.

Shikina S, Lin T-C, Chu Y-L, Cheng Y-C, Chang Y-E, Wada N, Tang S-L, Iizuka Y, Chiu Y-L. 2023. Culturing reef-building corals on a laboratory dish: a simple experimental platform for stony corals. Front. Mar. Sci. [Internet] 10. Available from: 10.3389/fmars.2023.1149495

Shubin N, Tabin C, Carroll S. 2009. Deep homology and the origins of evolutionary novelty. Nature 457:818–823.

Simakov O, Bredeson J, Berkoff K, Marletaz F, Mitros T, Schultz DT, O’Connell BL, Dear P, Martinez DE, Steele RE, et al. 2022. Deeply conserved synteny and the evolution of metazoan chromosomes. Sci Adv 8:eabi5884.

Simão FA, Waterhouse RM, Ioannidis P, Kriventseva EV, Zdobnov EM. 2015. BUSCO: assessing genome assembly and annotation completeness with single-copy orthologs. Bioinformatics 31:3210–3212.

Smit AFA, Hubley R, Green P. 2015. RepeatMasker Open-4.0.

Stanke M, Diekhans M, Baertsch R, Haussler D. 2008. Using native and syntenically mapped cDNA alignments to improve de novo gene finding. Bioinformatics 24:637–644.

Stanke M, Schöffmann O, Morgenstern B, Waack S. 2006. Gene prediction in eukaryotes with a generalized hidden Markov model that uses hints from external sources. BMC Bioinformatics 7:62.

Stanley GD Jr. 2003. The evolution of modern corals and their early history. Earth Sci. Rev. 60:195–225.

Steenwyk JL, Buida TJ 3rd, Li Y, Shen X-X, Rokas A. 2020. ClipKIT: A multiple sequence alignment trimming software for accurate phylogenomic inference. PLoS Biol. 18:e3001007.

Steenwyk JL, Buida TJ, Labella AL, Li Y, Shen X-X, Rokas A. 2021. PhyKIT: a broadly applicable UNIX shell toolkit for processing and analyzing phylogenomic data. Bioinformatics 37:2325–2331.

Steenwyk JL, King N. 2024. The promise and pitfalls of synteny in phylogenomics. PLoS Biol. 22:e3002632.

Stolarski J, Coronado I, Murphy JG, Kitahara MV, Janiszewska K, Mazur M, Gothmann AM, Bouvier A-S, Marin-Carbonne J, Taylor ML, et al. 2021. A modern scleractinian coral with a two-component calcite-aragonite skeleton. Proc. Natl. Acad. Sci. U. S. A. 118:e2013316117.

Stolarski J, Kitahara MV, Miller DJ, Cairns SD, Mazur M, Meibom A. 2011. The ancient evolutionary origins of Scleractinia revealed by azooxanthellate corals. BMC Evol. Biol. 11:316.

Struck TH. 2026. Progress and challenges in phylogenomics and genomics of Lophotrochozoa/Spiralia. Zool. Scr. [Internet]. Available from: 10.1111/zsc.70056

Takeuchi T, Yamada L, Shinzato C, Sawada H, Satoh N. 2016. Stepwise evolution of coral biomineralization revealed with genome-wide proteomics and transcriptomics. PLoS One 11:e0156424.

Tarashansky AJ, Musser JM, Khariton M, Li P, Arendt D, Quake SR, Wang B. 2021. Mapping single-cell atlases throughout Metazoa unravels cell type evolution. Elife 10:e66747.

The Darwin Tree of Life Project Consortium, Blaxter M, Mieszkowska N, Palma FD, Holland P, Durbin R, Richards T, Berriman M, Kersey P, Hollingsworth P, et al. 2022. Sequence locally, think globally: The Darwin Tree of Life Project. Proceedings of the National Academy of Sciences 119:e2115642118.

Threlfall J, Blaxter M. 2021. Launching the Tree of Life Gateway. Wellcome Open Res. 6:125.

Tinoco AI, Mitchison-Field LMY, Bradford J, Renicke C, Perrin D, Bay LK, Pringle JR, Cleves PA. 2023. Role of the bicarbonate transporter SLC4γ in stony-coral skeleton formation and evolution. Proc. Natl. Acad. Sci. U. S. A. 120:e2216144120.

True JR, Carroll SB. 2002. Gene co-option in physiological and morphological evolution. Annu. Rev. Cell Dev. Biol. 18:53–80.

Vaga CF, Quattrini AM, Galvão de Lossio E, Seiblitz I, Huang D, Quek ZBR, Stolarski J, Cairns SD, Kitahara MV. 2025. A global coral phylogeny reveals resilience and vulnerability through deep time. Nature 648:377–382.

Vaga CF, Seiblitz IG de L e., Capel KCC, Quattrini AM, Stolarski J, Cairns SD, Kitahara MV. 2024. Untangling deep-sea corals systematics: Description of a new family, Stephanocyathidae (Anthozoa, Scleractinia), through a genomic approach. Zool. Scr. 53:473–486.

Wang J -T, Chen Y-Y, Tew KS, Meng P-J, Chen CA. 2012. Physiological and biochemical performances of menthol-induced aposymbiotic corals. PLoS One 7:e46406.

Wang X, Drillon G, Ryu T, Voolstra CR, Aranda M. 2017. Genome-based analyses of six hexacorallian species reject the “naked coral” hypothesis. Genome Biol. Evol. 9:2626– 2634.

Wang X, Zoccola D, Liew YJ, Tambutte E, Cui G, Allemand D, Tambutte S, Aranda M. 2021. The evolution of calcification in reef-building corals. Mol. Biol. Evol. 38:3543–3555.

Wang Y, Tang H, Debarry JD, Tan X, Li J, Wang X, Lee T-H, Jin H, Marler B, Guo H, et al. 2012. MCScanX: a toolkit for detection and evolutionary analysis of gene synteny and collinearity. Nucleic Acids Res. 40:e49.

Wickham H. 2009. Ggplot2: Elegant graphics for data analysis. 1st ed. New York, NY: Springer

Wong TKF, Ly-Trong N, Ren H, Demotte P, Baños H, Roger AJ, Susko E, Bielow C, De Maio N, Goldman N, et al. 2026. IQ-TREE 3: phylogenomic inference software using complex evolutionary models. Mol. Biol. Evol. [Internet] 43. Available from: 10.1093/molbev/msag117

Wood C, Bishop J, Harley J, Mrowicki R, Marine Biological Association Genome Acquisition Lab, Darwin Tree of Life Barcoding collective, Wellcome Sanger Institute Tree of Life programme, Wellcome Sanger Institute Scientific Operations: DNA Pipelines collective, Tree of Life Core Informatics collective, Darwin Tree of Life Consortium. 2022. The genome sequence of the orange-striped anemone, Diadumene lineata (Verrill, 1869). Wellcome Open Res. 7:93.

Xiao M, Brugler MR, Broe MB, Gusmão LC, Daly M, Rodríguez E. 2019. Mitogenomics suggests a sister relationship of Relicanthus daphneae (Cnidaria: Anthozoa: Hexacorallia: incerti ordinis) with Actiniaria. Sci. Rep. 9:18182.

Yang Z, Rannala B. 2006. Bayesian estimation of species divergence times under a molecular clock using multiple fossil calibrations with soft bounds. Mol. Biol. Evol. 23:212–226.

Ying H, Cooke I, Sprungala S, Wang W, Hayward DC, Tang Y, Huttley G, Ball EE, Forêt S, Miller DJ. 2018. Comparative genomics reveals the distinct evolutionary trajectories of the robust and complex coral lineages. Genome Biol. 19:175.

Young MD, Behjati S. 2020. SoupX removes ambient RNA contamination from droplet-based single-cell RNA sequencing data. Gigascience 9:giaa151.

Zhang C, Mirarab S. 2022. Weighting by gene tree uncertainty improves accuracy of quartet-based species trees. Mol. Biol. Evol. 39:msac215.

Zhang C, Rabiee M, Sayyari E, Mirarab S. 2018. ASTRAL-III: polynomial time species tree reconstruction from partially resolved gene trees. BMC Bioinformatics 19:153.

Zhao T, Zwaenepoel A, Xue J-Y, Kao S-M, Li Z, Schranz ME, Van de Peer Y. 2021. Whole-genome microsynteny-based phylogeny of angiosperms. Nat. Commun. 12:3498.

Zhou C, McCarthy SA, Durbin R. 2022. YaHS: yet another Hi-C scaffolding tool. Bioinformatics 39:btac808.

Zimmermann B, Montenegro JD, Robb SMC, Fropf WJ, Weilguny L, He S, Chen S, Lovegrove-Walsh J, Hill EM, Chen C-Y, et al. 2023. Topological structures and syntenic conservation in sea anemone genomes. Nat. Commun. 14:8270.

Zoccola D, Ganot P, Bertucci A, Caminiti-Segonds N, Techer N, Voolstra CR, Aranda M, Tambutté E, Allemand D, Casey JR, et al. 2015. Bicarbonate transporters in corals point towards a key step in the evolution of cnidarian calcification. Sci. Rep. 5:9983.

Zolotarov G, Grau-Bové X, Sebé-Pedrós A. 2026. GeneExt: a gene model extension tool for enhanced single-cell RNA-seq analysis. Bioinformatics [Internet] 42. Available from: 10.1093/bioinformatics/btag094

