## Supplementary material for "Corallimorpharian genome supports the monophyly of Scleractinia and illuminates the cellular evolution of coral calcification"

**This PDF file includes**

- Supplementary Figures 1–19
- Supplementary Tables 1–5
- Legends for Supplementary Datasets 1–6

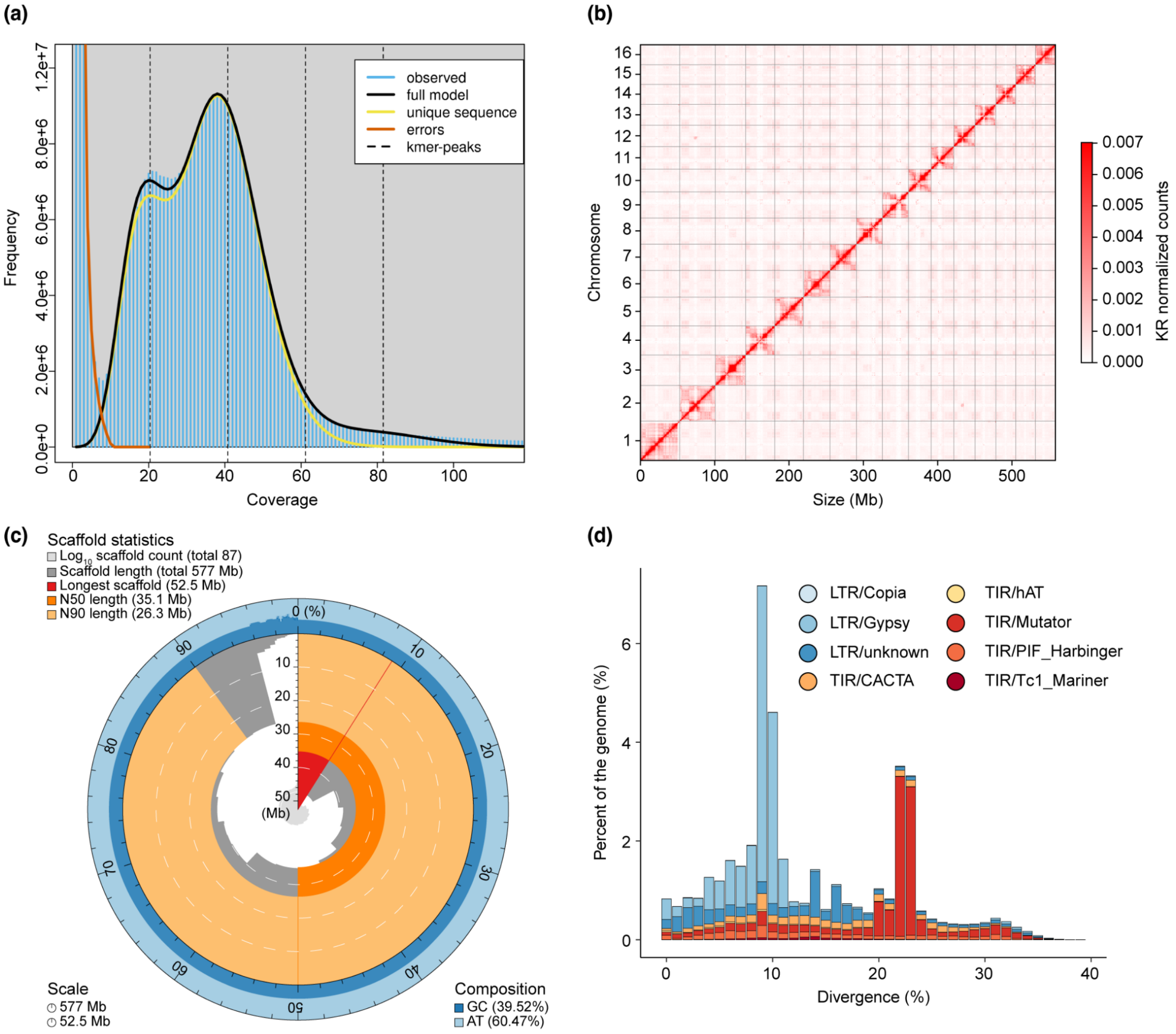


**Supplementary Figure 1.** A chromosome-level genome for the corallimorpharian *Ricordea yuma*. (a) GenomeScope profile based on *R. yuma* PacBio HiFi sequencing reads. Genome size (410 Mb) and heterozygosity (1.02%) are measured using a kmer-based approach ($k$ = 21). (b) Hi-C contact map (bin size = 500 kb) for the *R. yuma* assembly generated with HapHiC, normalized with the Knight–Ruiz algorithm to remove technical biases. Darker colors indicate stronger interactions. (c) Snailplot for the final *R. yuma* assembly showing total length (577 Mb), scaffold N50 (35.1 Mb), scaffold N90 (26.1 Mb) and GC content (39.5%). (d) Repeat landscape of the *R. yuma* genome. Divergence is used as a measure of time—higher levels of divergence indicate an older element. Peaks indicate a transposon invasion at a specific timepoint.

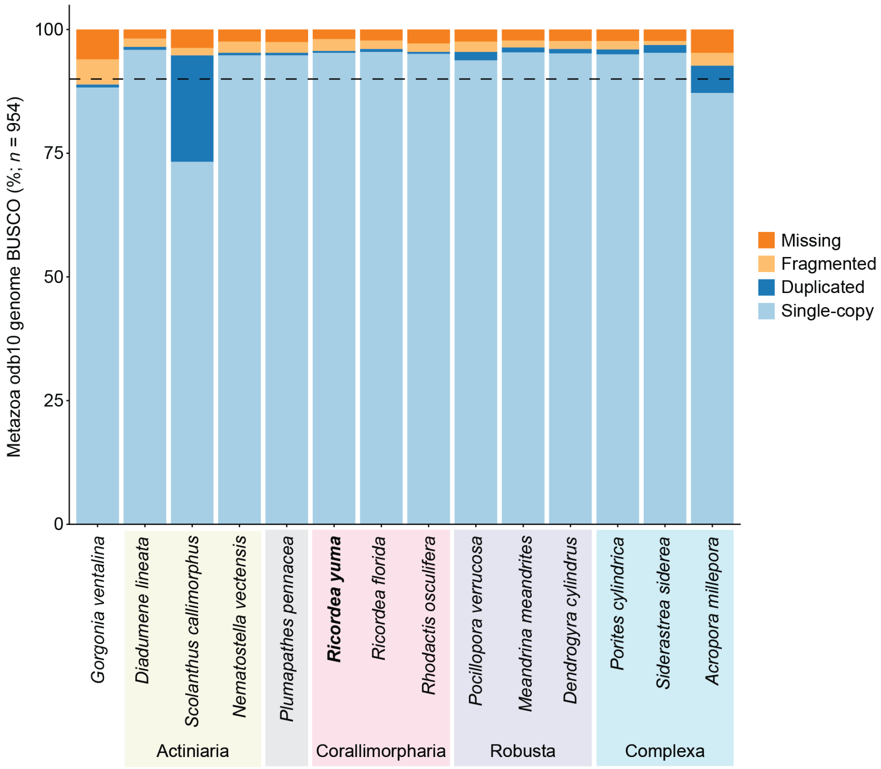


**Supplementary Figure 2.** Benchmarking Universal Single Copy Ortholog (BUSCO) scores for genomes used in this work. The genome of *R. yuma* (bold) was sequenced as part of this study. BUSCO scores were assessed using the metazoa_odb10 database, consisting of 954 universal metazoan genes. All genomes used in synteny analyses have BUSCO complete rates (single-copy plus duplicated) over 90% (dashed line).

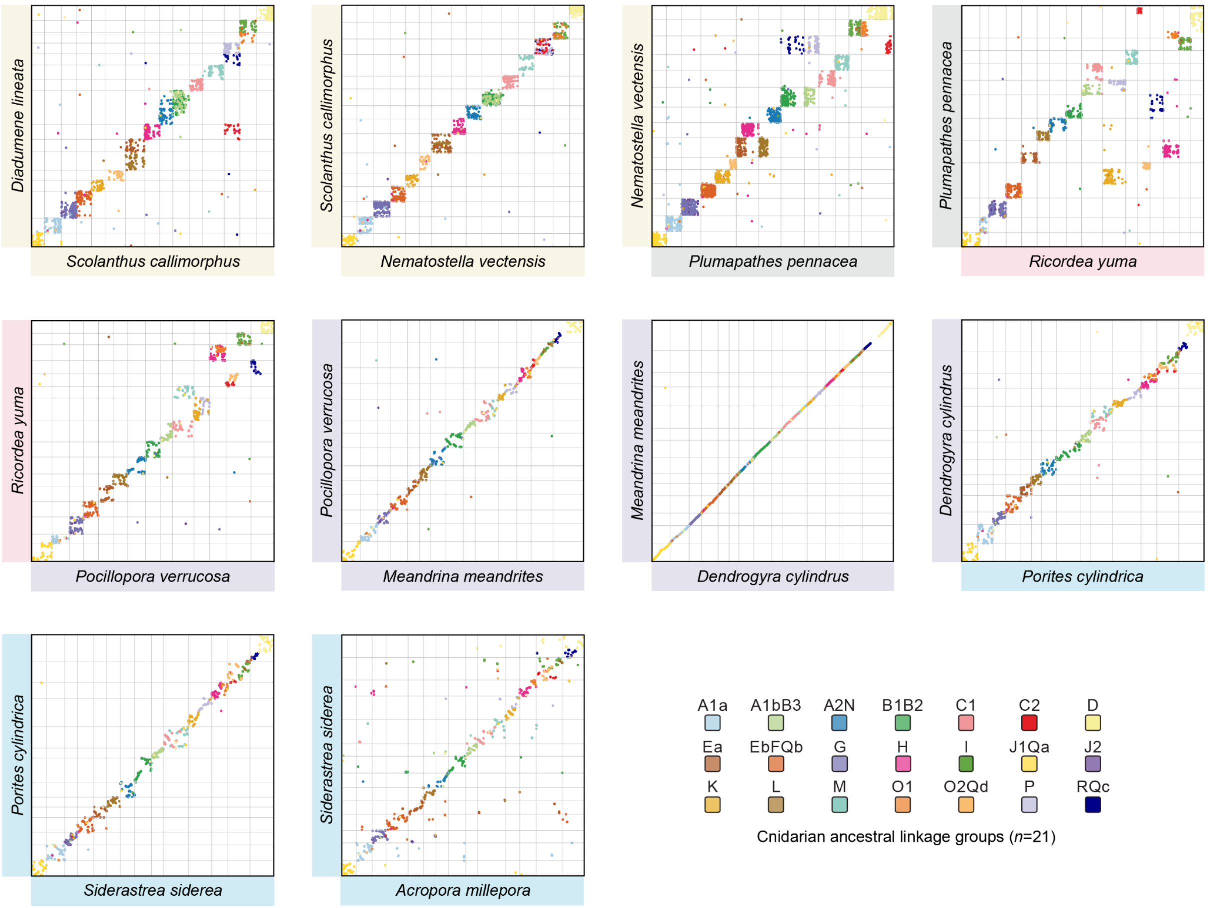


**Supplementary Figure 3.** Dot plots associated with the main text Fig. 1d. Each plot shows a pairwise comparison of genomes, with points representing the position of an ortholog in both genomes. Points are colored by the cnidarian ALG to which the gene belongs. Gray lines delineate chromosomal boundaries.

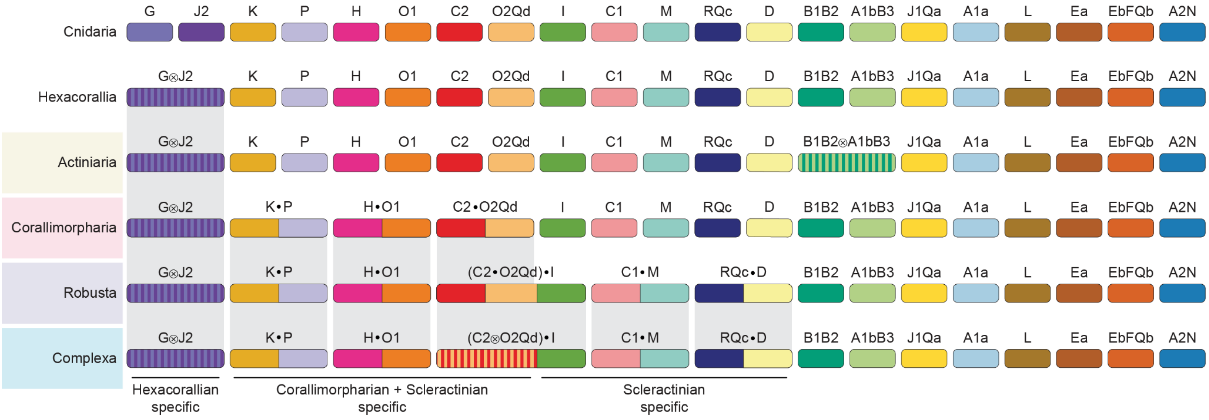


**Supplementary Figure 4.** Chromosome fusion events shared by hexacorallian clades. The last common ancestor of cnidarians had 21 ALGs. The last common ancestor of hexacorallians had a reduced complement of 20 ALGs due to an irreversible fusion-with-mixing event (G⨂J2) that all hexacorals have subsequently inherited. Corallimorpharians and scleractinians of both complex and robust clades share three subsequent fusions: K●P, H●O1 and C2●O2Qd. Finally, three additional fusions are shared by both robust and complex scleractinians but not corallimorpharians: I●(C2●O2Qd), C1●M and D●RQc. Actiniarians have a lineage-specific B1B2⨂A1bB3 fusion. Striped chromosomes represent fusion-with-mixing. Partitioned chromosomes represent fusion-without-mixing.

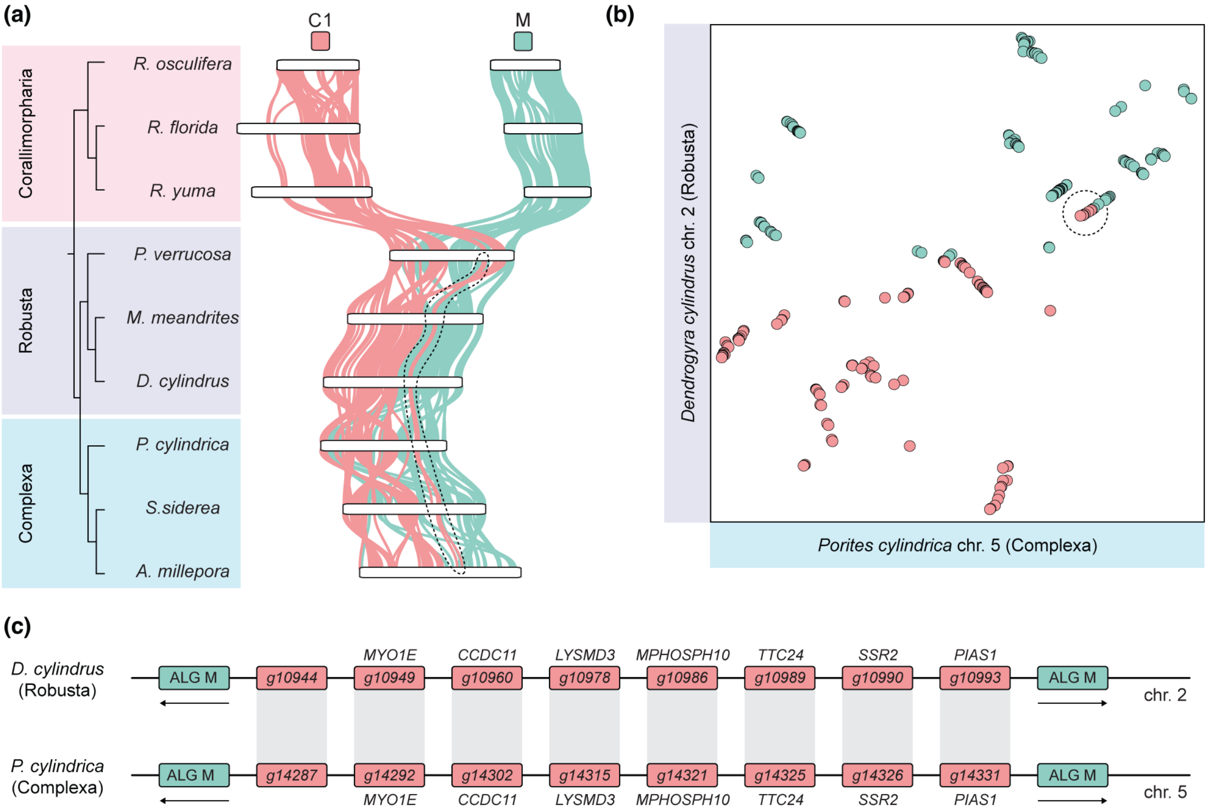


**Supplementary Figure 5.** The C1●M fusion occurred only once, in the ancestor of scleractinians, and did not occur independently in robust and complex corals. (a) Ribbon plot of chromosomes containing ALGs C1 and M in stony coral and corallimorpharian genomes. Dashed outline highlights a group of eight orthologs from the C1 ALG that have been translocated from the C1 portion of the chromosome to the M portion. This translocation is itself a rare genomic change. Since it is highly unlikely that exactly the same translocation of the same eight genes to the same location would occur independently in robust and complex corals, it must have happened in the ancestor of scleractinians. This means that the C1●M fusion must predate the complex/robust split, and therefore occurred only once, not twice, independently in each lineage. (b) Dot plot showing position of genes from the C1 and M ALGs on *Dendrogyra cylindrus* (Robusta) chromosome 2 and *Porites cylindrica* (Complexa) chromosome 5. The eight translocated ALG C1 genes are marked by the dotted circle. Each point represents one gene, colored by the ALG to which it belongs. (c) Simplified representation of the eight translocated ALG C1 orthologs on *Dendrogyra cylindrus* (Robusta) chromosome 2 and *Porites cylindrica* (Complexa) chromosome 5. Black line represents the chromosome, colored squares represent genes.

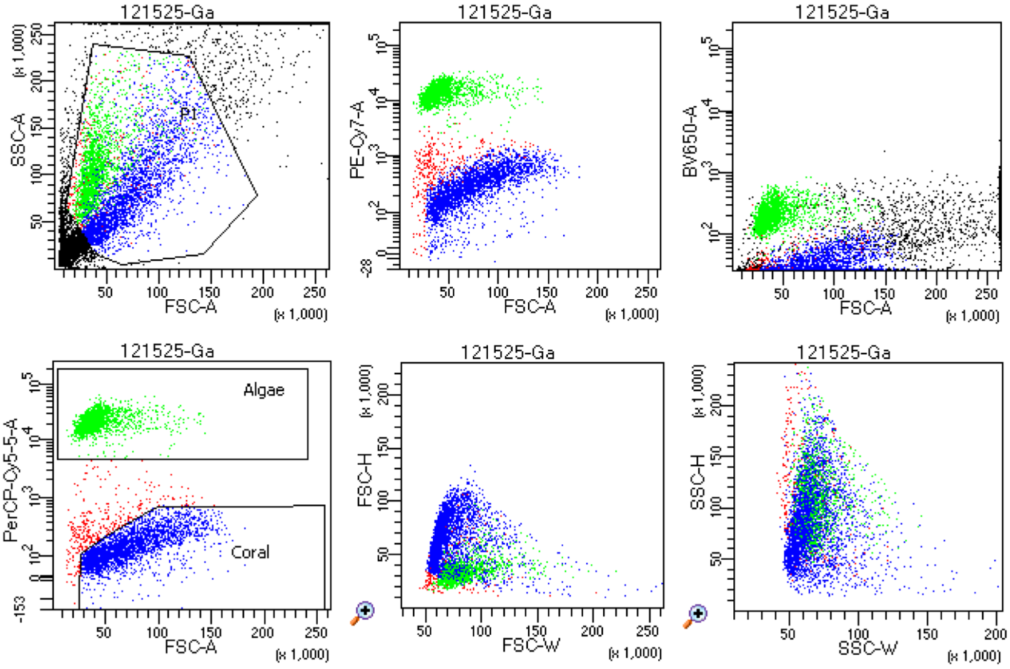


Tube: Ga

| Population | #Events | %Parent | %Total |
|---|---|---|---|
| All Events | 10,000 | #### | 100.0 |
| P1 | 5,019 | 50.2 | 50.2 |
| Algae | 1,991 | 39.7 | 19.9 |
| Coral | 2,674 | 53.3 | 26.7 |

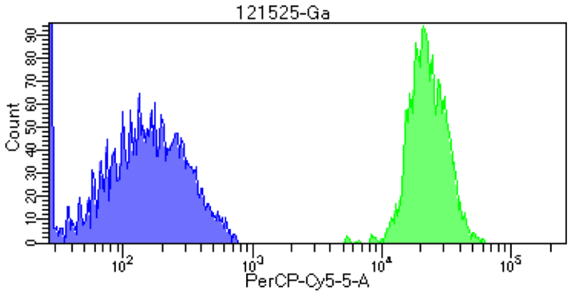


**Supplementary Figure 6.** FACS gating strategy for separating animal host cells and symbiotic Symbiodiniaceae from *Galaxea fascicularis*. Cellular events were first gated using forward scatter area (FSC-A) and side scatter area (SSC-A) to exclude debris, followed by singlet selection based on FSC-H/FSC-W and SSC-H/SSC-W profiles. Symbiotic algae (green) and coral animal cells (blue) were distinguished by chlorophyll autofluorescence detected in the PerCP-Cy5.5 channel following 488 nm laser excitation. Symbiodiniaceae cells exhibited strong autofluorescence, whereas coral animal cells displayed substantially lower fluorescence intensity. Low-autofluorescence singlet coral cells were isolated for downstream analyses. To determine the optimal filter combination for resolving autofluorescence signals from symbiotic algae, cells were analyzed using three optical configurations: PE-Cy7 (Ex: 561 nm, Em: 780/60 nm), BV650 (Ex: 405 nm, Em: 660/20 nm) and PerCP-Cy5.5 (Ex: 488 nm, Em: 695/40 nm).

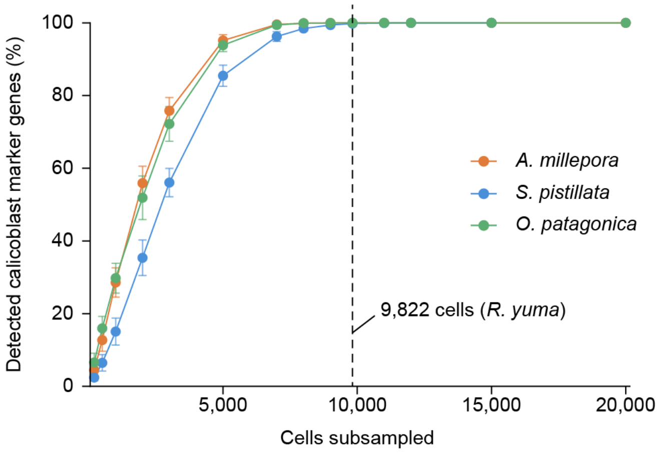


**Supplementary Figure 7.** Subsampling analysis supports the biological absence of calicoblasts in the corallimorpharian *R. yuma*. Cells from *S. pistillata*, *O. patagonica*, and *A. millepora* were randomly subsampled and downsampled to match the sequencing depth of *R. yuma*. At the *R. yuma* cell count (9,822 cells), the proportion of detectable calicoblast marker genes remained nearly complete (99.80% in *S. pistillata*, 99.99% in *O. patagonica*, and 99.99% in *A. millepora*), indicating that the absence of calicoblast markers in *R. yuma* is unlikely to be explained by insufficient cell number or sequencing depth.

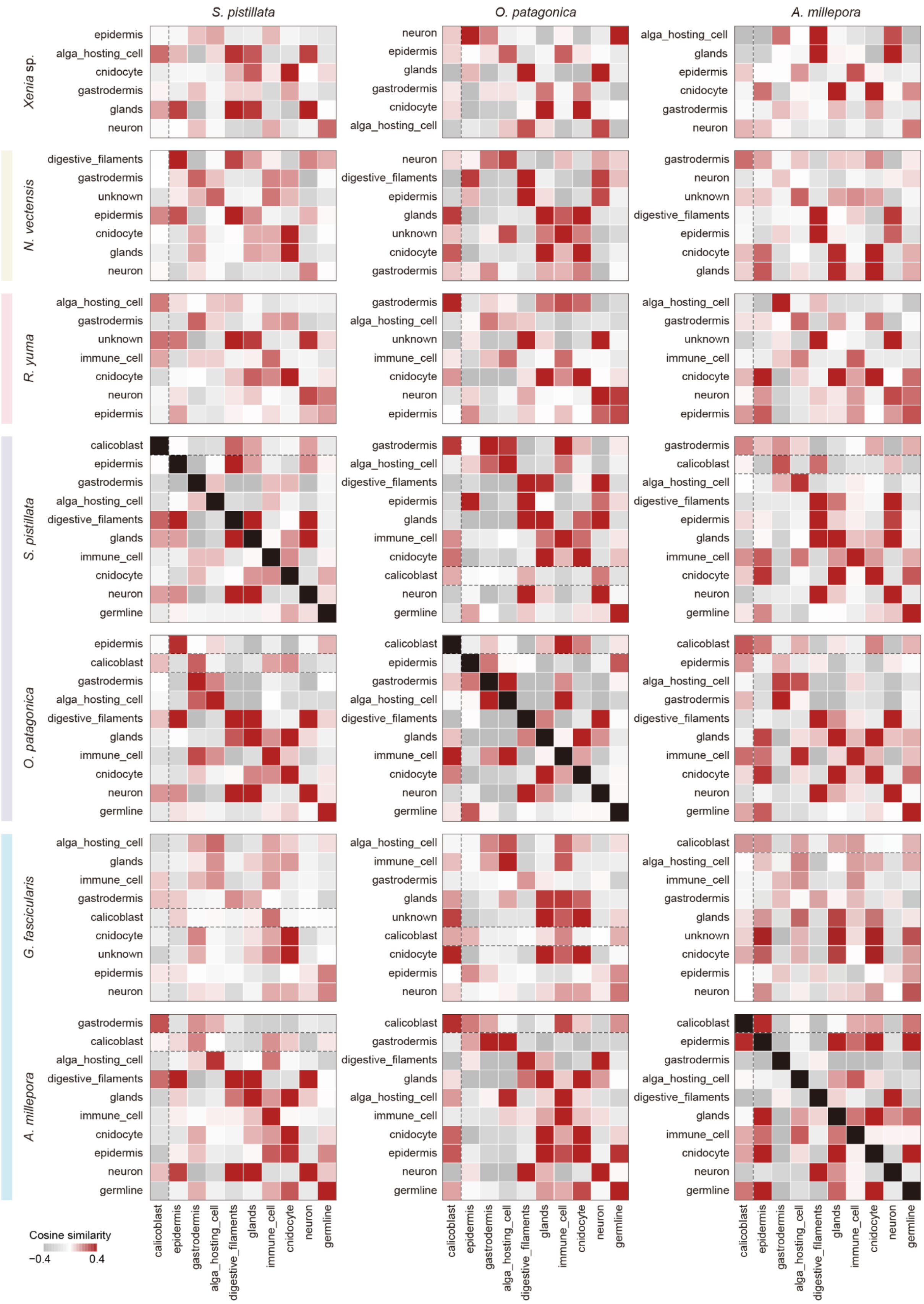


**Supplementary Figure 8.** All-to-all cross-species comparison of anthozoan cell types based on cosine similarity in a shared embedding space generated by TranscriptFormer.

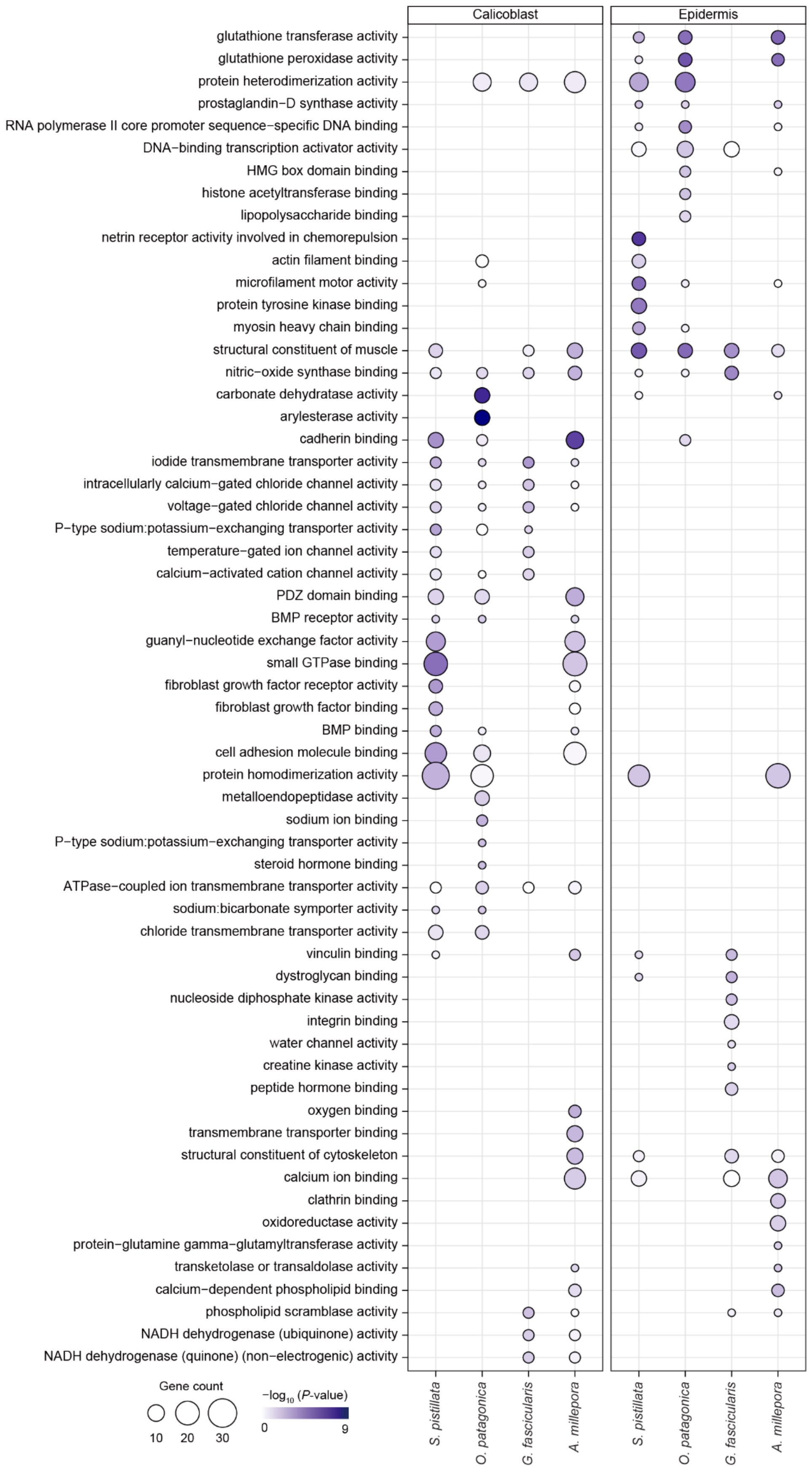


**Supplementary Figure 9.** Gene Ontology (GO) enrichment analysis of calicoblasts and epidermis in scleractinian corals. Dot plot shows the top 10 enriched molecular function GO terms for each species.

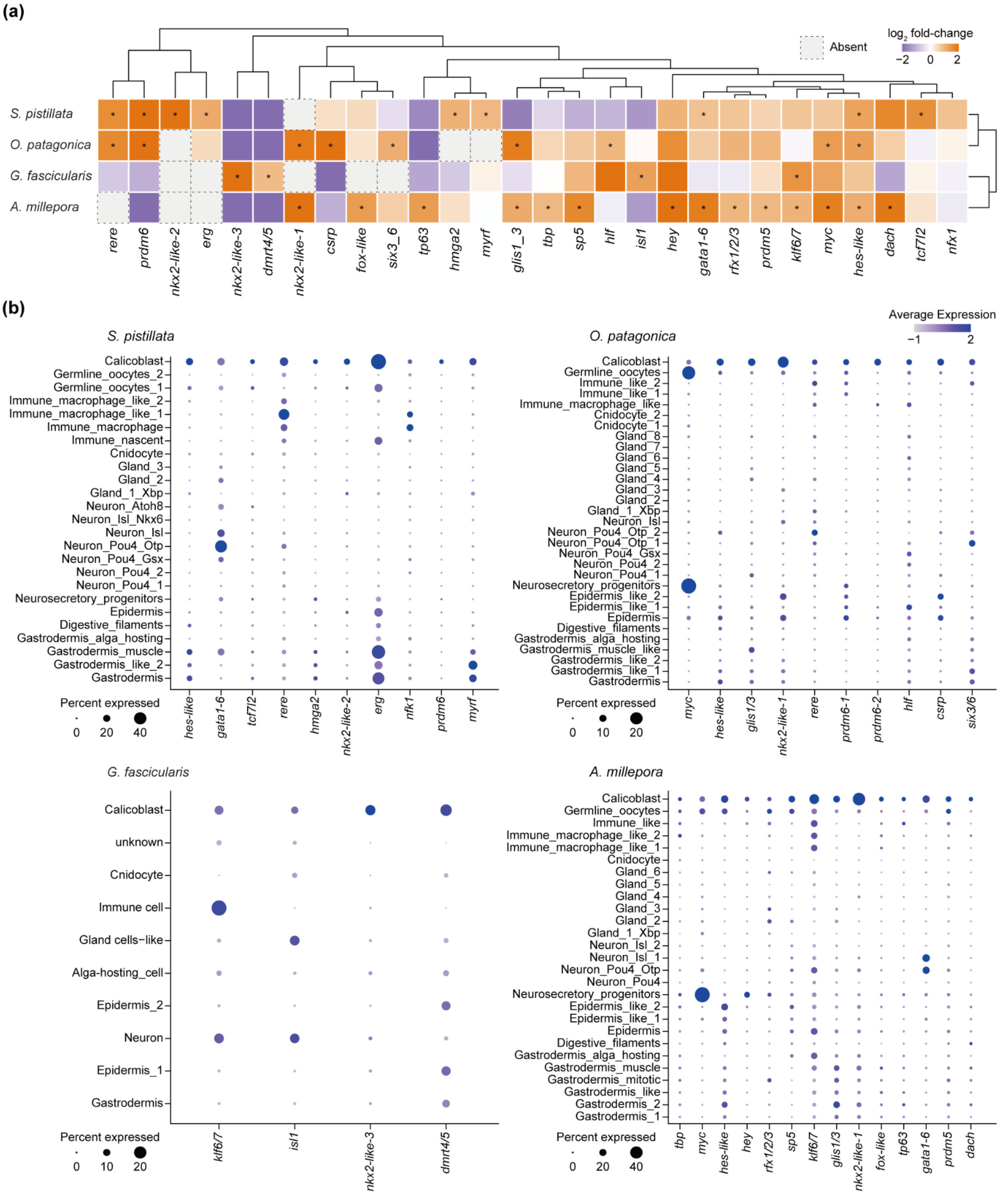


**Supplementary Figure 10.** Transcription factors differentially expressed in calicoblasts. (a) Heatmap with hierarchical clustering showing calicoblast-enriched transcription factors across scleractinian corals. Asterisks indicate significant enrichment ($\log_2$ fold-change > 1, expression detected in > 10% of cells, and $P < 0.05$). (b) Dot plots showing the distribution and expression levels of the enriched transcription factors across all cell types in each species. Transcription factor annotations were assigned based on best-hit KEGG orthologs and were revised using phylogenetic analyses and previous reports (Levy et al. 2021; Levy et al. 2025).

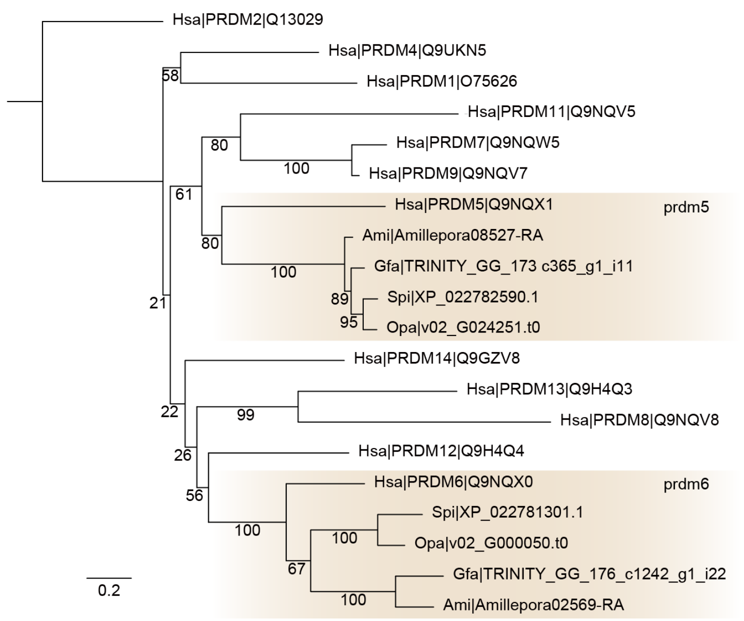


**Supplementary Figure 11.** Phylogenetic tree showing orthology assignment of *prdm* genes. Maximum-likelihood analysis was conducted under the WAG+F+R3 model with 1,000 bootstrap replicates. Node labels indicate the percentage of bootstrap support. Species codes: Hsa, *Homo sapiens*; Spi, *Stylophora pistillata*; Opa, *Oculina patagonica*; Gfa, *Galaxea fascicularis*; Ami, *Acropora millepora*.

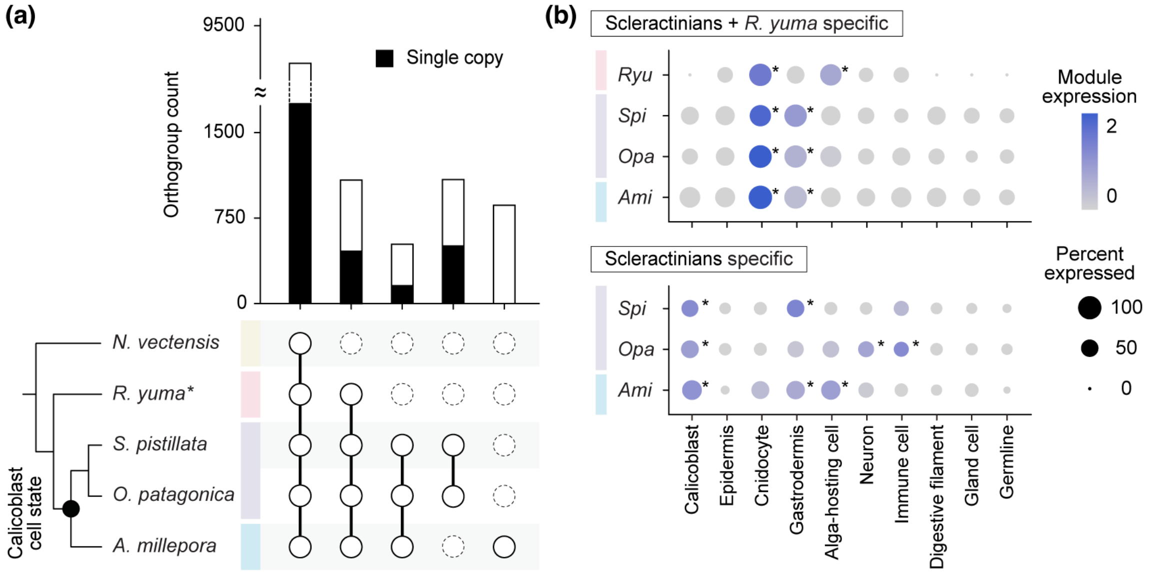


**Supplementary Figure 12.** Lineage-specific genes display cell type-specific expression patterns. (a) An UpSet plot shows the composition of lineage-specific orthogroups (OGs) identified by OrthoFinder across major clades. White bars indicate the total number of OGs, whereas black bars indicate the subset of these OGs that are single-copy. The matrix below indicates the combination of species corresponding to each bar. (b) Dot plot showing the average module scores calculated from lineage-specific single-copy orthogroups of the scleractinians + *R. yuma* clade (454 genes) and the scleractinian clade (147 genes). Asterisks indicate module scores that are significantly higher than those in all except two other cell types (*t*-test, $P < 0.01$). *Ryu*, *Ricordea yuma*; *Spi*, *Stylophora pistillata*; *Opa*, *Oculina patagonica*; *Ami*, *Acropora millepora*.

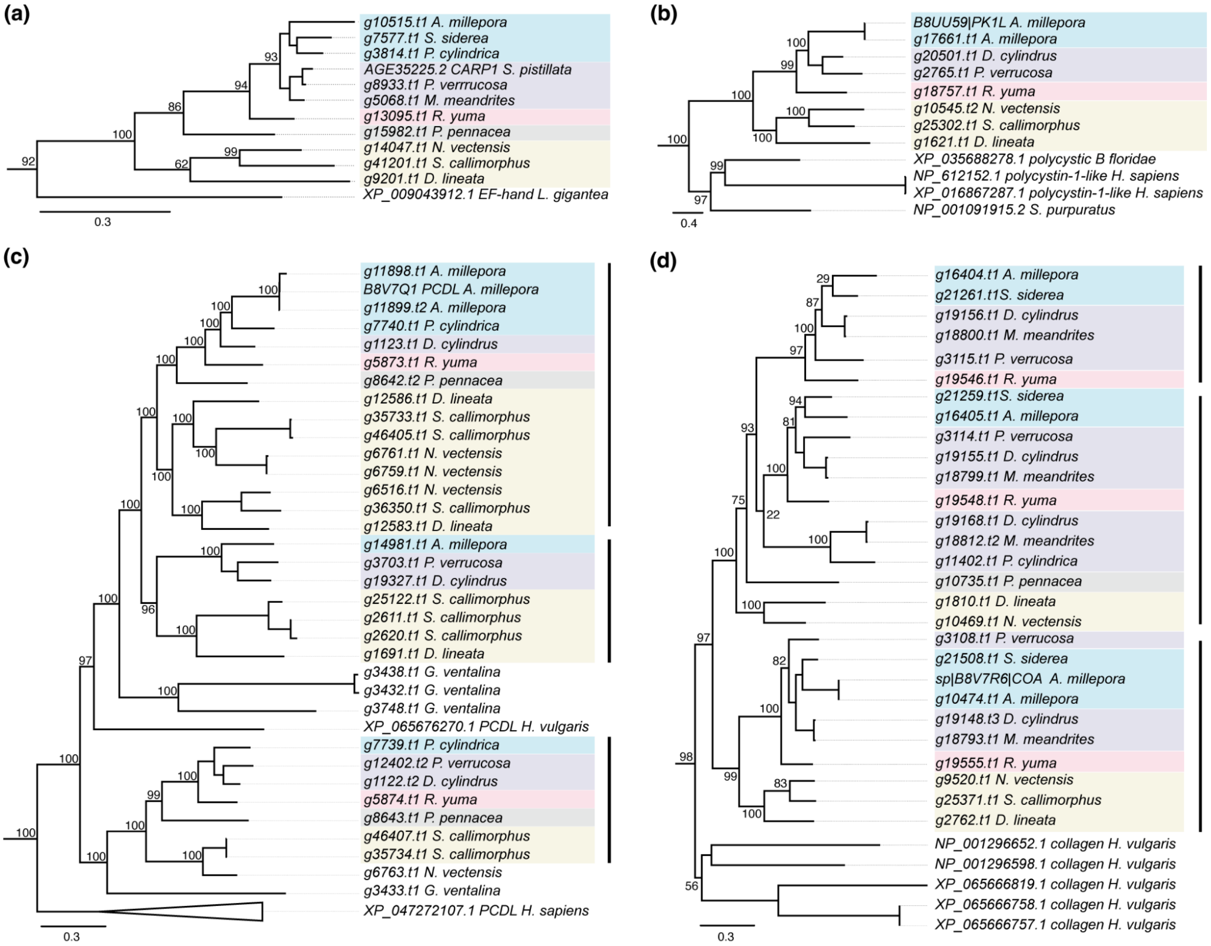


**Supplementary Figure 13.** Distribution of SOMP genes across Hexacorallian taxa inferred from gene tree analysis. Gene trees were constructed to assess the presence and absence of skeletal organic matrix protein (SOMP) genes across Hexacorallian lineages. (a) Distribution of the coral acid-rich protein 1 (CARP1) gene across Hexacorallian taxa. (b) Distribution of the polycystin-like 1 (PK1) gene across Hexacorallian taxa. (c) Gene tree of the protocadherin-like (PCDL) gene in Hexacorallians, with black bars indicating three hexacorallian PCDL copies. (d) Gene tree of the alpha collagen (COA) gene in Hexacorallians, with black bars indicating two broadly distributed hexacorallian COA copies and one copy restricted to Scleractinia and Corallimorpharia. Branch colors indicate taxonomic level: blue: Complexa (Scleractinia); purple: Robusta (Scleractinia); pink: Corallimorpharia; gray: Antipatharia; yellow: Actiniaria.

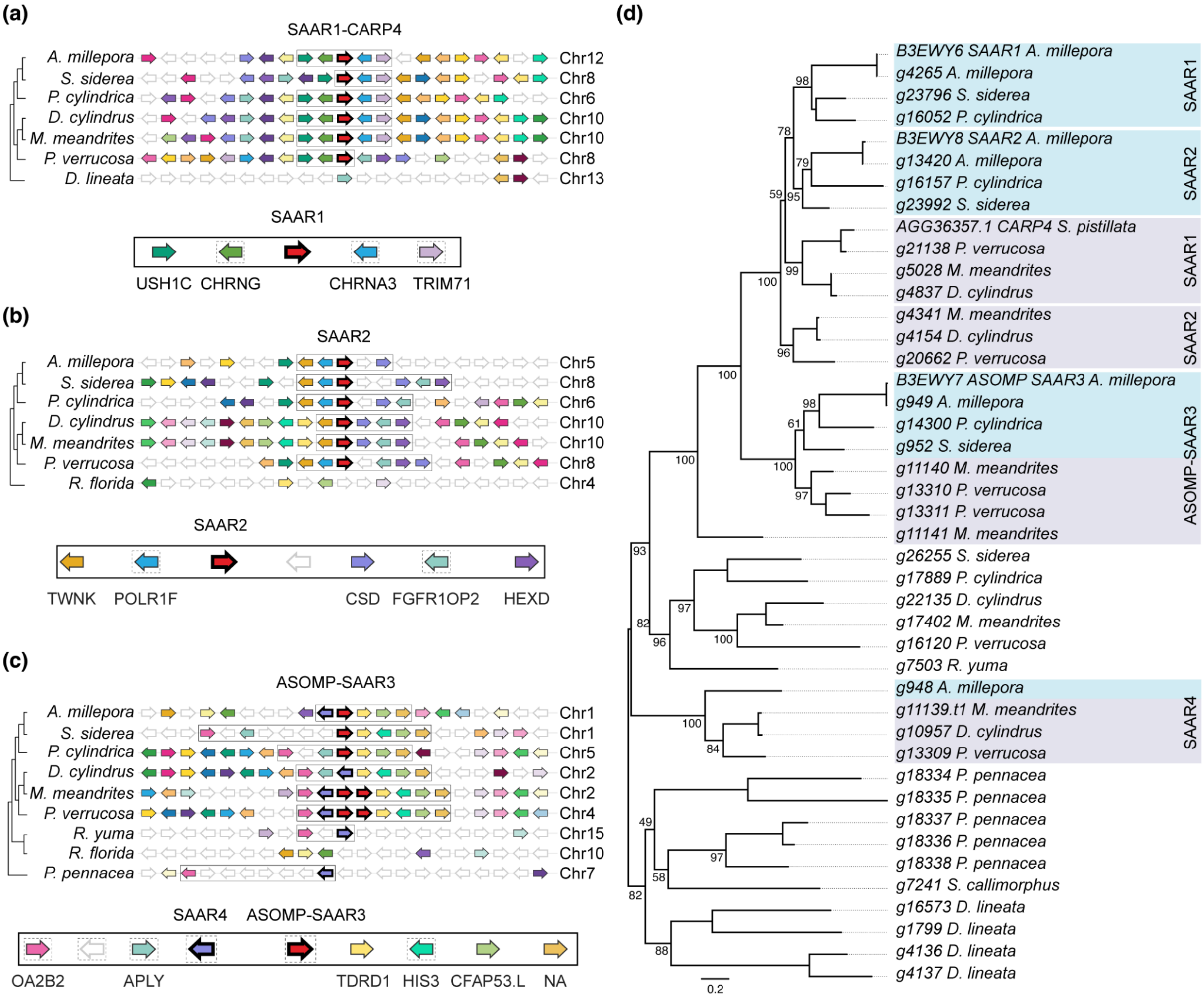


**Supplementary Figure 14**. Microsynteny and phylogenetic distribution of Skeletal Aspartic Acid-Rich (SAAR) genes across hexacorallian lineages. (a–c) Microsynteny plots showing the genomic neighbourhood of SAAR1, SAAR2, SAAR3 (ASOMP), and the putative SAAR4 across coral species. In each panel, the gene of interest is highlighted in red, and rectangles delimit conserved syntenic blocks. Arrow orientation indicates transcriptional strand, with rightward arrows representing the forward strand (+) and leftward arrows the reverse strand (−). Colored arrows represent orthologs across species, while white arrows with gray outlines indicate genes lacking a clear ortholog in the genomic region surveyed. Below each microsynteny plot, a schematic summarises the conserved syntenic neighbourhood of the gene of interest, where arrows with a dashed square indicate neighbour genes that may be absent in some species. (a) SAAR1 (also known as CARP4 in robusta species) is a coral-specific gene whose conserved syntenic neighbourhood includes *USH1C* (Harmonin), *CHRNG* (Acetylcholine receptor subunit gamma), *CHRNA3* (Neuronal acetylcholine receptor subunit alpha-3), and *TRIM71* (E3 ubiquitin-protein ligase TRIM71). (b) SAAR2 is likewise coral lineage-specific, with a conserved neighbourhood comprising *TWNK* (Twinkle mtDNA helicase), *POLR1F* (DNA-directed RNA polymerase I subunit RPA43), *CSD* (Cysteine desulfurase SufS), *FGFR1OP2* (FGFR1 oncogene partner 2), and *HEXD* (Hexosaminidase D). (c) SAAR3 (also known as ASOMP) and the putative novel paralogue SAAR4 share a syntenic neighbourhood anchored by *OA2B2* (Octopamine receptor beta-2R), *APLY* (Aplysianin-A), *TDRD1* (Tudor domain-containing protein 1), *HIS3* (Histone H3), *CFAP53.L* (Cilia- and flagella-associated protein 53), and an uncharacterised ortholog (NA). (d) Phylogenetic tree of SAAR1, SAAR2, SAAR3, and putative SAAR4 gene models. Blue and purple squares correspond to Complexa and Robusta clades, respectively. Gene models associated with SAAR4 are highlighted in red; although they form a coral-specific clade, the phylogenetic placement of the corallimorpharian and black coral sequences remains unresolved.

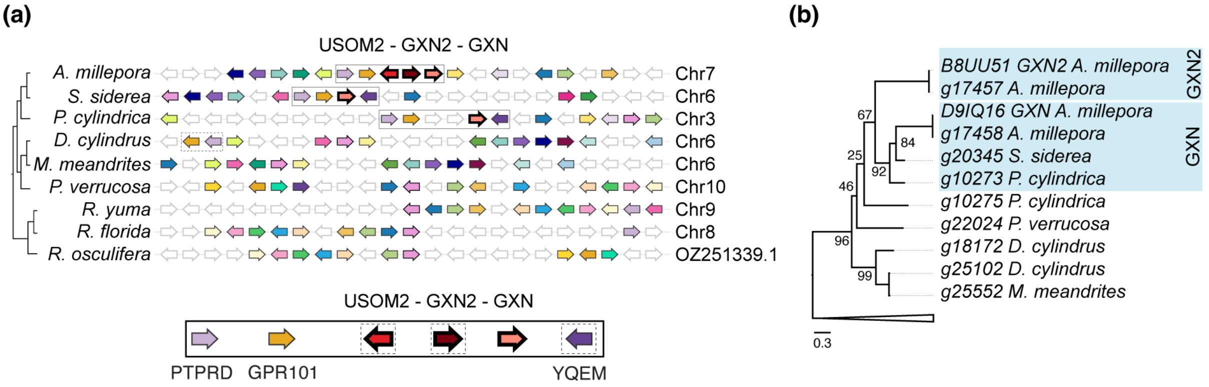


**Supplementary Figure 15**. Microsynteny and phylogenetic distribution of Galaxin 1 and 2 (GXN and GXN2) genes and Uncharacterized Skeletal Organic Matrix 2 (USOM2) genes across hexacorallian lineages. (a) Microsynteny plot showing the genomic neighbourhood of GXN, GXN2, and USOM2 across coral species. The genes of interest are highlighted in different tones of red with black outlines, and rectangles delimit its conserved syntenic block. Arrow orientation indicates transcriptional strand, with rightward arrows representing the forward strand (+) and leftward arrows the reverse strand (−). Colored arrows represent orthologs across species, while white arrows with gray outlines indicate genes lacking a clear ortholog in the genomic region surveyed. Below the microsynteny plot, a schematic summarises the conserved syntenic neighbourhood of the genes of interest, where arrows with a dashed square indicate neighbour genes that may be absent in some species. GXN, GXN, and USOM2 are arranged as a continuous genomic cluster, GXN2 and USOM2 are restricted to *Acropora*, while GXN (Galaxin 1) is found exclusively in Complexa corals. Their shared conserved syntenic neighbourhood includes *PTPRD* (Receptor-type tyrosine-protein phosphatase delta), *GPR101* (Probable G-protein coupled receptor 101), and *YQEM* (Putative methyltransferase YqeM). (b) Phylogenetic tree of GXN and GXN2 gene models. Blue squares correspond to Complexa clades. A phylogenetic tree for USOM2 is not presented, as BLAST searches recovered no additional hits beyond the *Acropora* sequence, precluding meaningful phylogenetic inference.

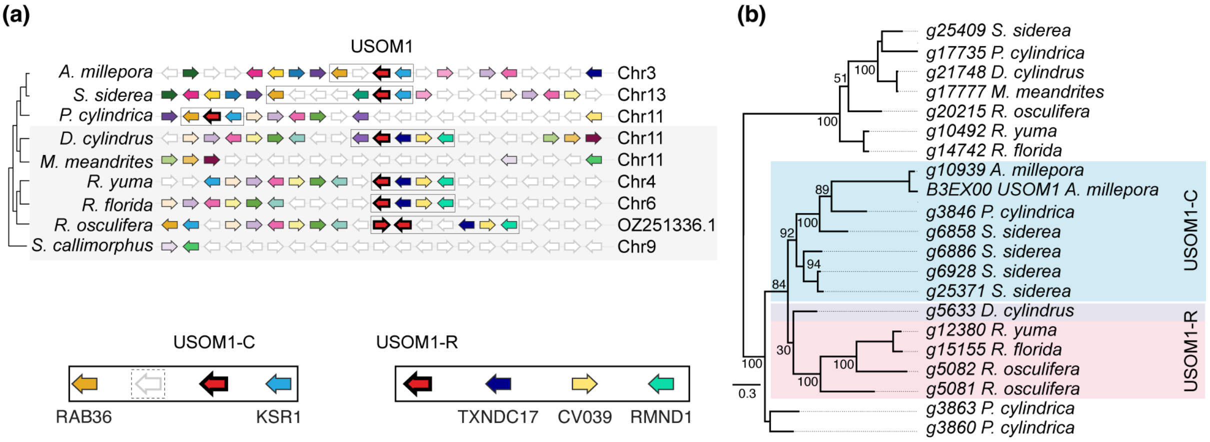


**Supplementary Figure 16**. Microsynteny and phylogenetic distribution of Uncharacterized Skeletal Organic Matrix protein 1 (USOM1) across hexacorallian lineages. (a) Microsynteny plot showing the genomic neighbourhood of USOM1 across hexacorallian species. The gene of interest is highlighted in red with black outlines, and rectangles delimit its conserved syntenic blocks. Arrow orientation indicates transcriptional strand, with rightward arrows representing the forward strand (+) and leftward arrows the reverse strand (−). Colored arrows represent orthologs across species, while white arrows with gray outlines indicate genes lacking a clear ortholog in the genomic region surveyed. Below the microsynteny plot, a schematic summarises the conserved syntenic neighbourhood of both Complexa and Robusta USOM1 variants (USOM1-C and USOM1-R), where arrows with a dashed square indicate neighbour genes that may be absent from the syntenic block in some species. Two distinct syntenic configurations of USOM1 were identified. The first is conserved across Complexa corals (USOM1-C) and is flanked by *RAB36* (Ras-related protein Rab-36) and *KSR1* (Kinase suppressor of Ras 1). The second syntenic configuration, shaded in gray, is shared by Robusta corals and corallimorpharians (USOM1-R), with a conserved neighbourhood comprising *TXNDC17* (Thioredoxin domain-containing protein 17), *CV039* (Synaptic plasticity regulator PANTS), and *RMND1* (Required for meiotic nuclear division protein 1 homolog). (b) Phylogenetic tree of USOM1 gene models. Blue, purple, and pink squares correspond to Complexa, Robusta, and corallimorpharian clades, respectively. Complexa sequences form a distinct branch separate from Robusta and corallimorpharian sequences, consistent with the two independent syntenic configurations observed in panel (a).

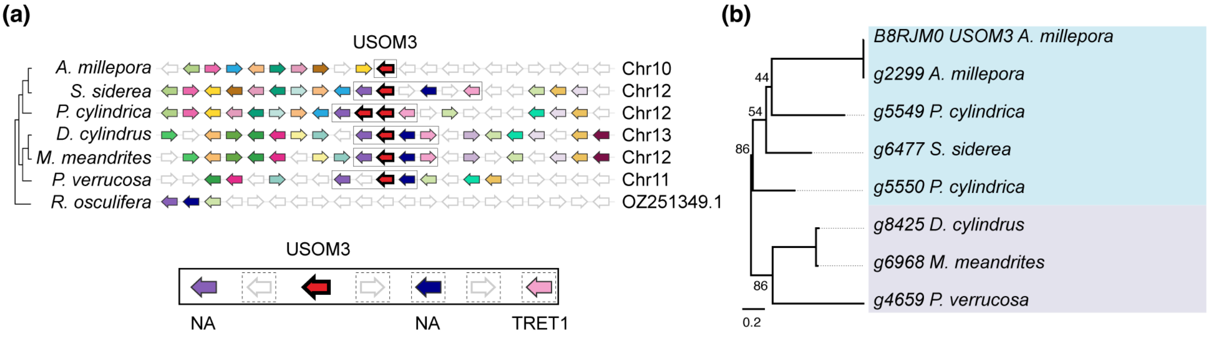


**Supplementary Figure 17**. Microsynteny and phylogenetic distribution of Uncharacterized Skeletal Organic Matrix protein 3 (USOM3) across coral lineages. (a) Microsynteny plot showing the genomic neighbourhood of USOM3 across coral species. The gene of interest is highlighted in red with black outlines, and rectangles delimit its conserved syntenic block. Arrow orientation indicates transcriptional strand, with rightward arrows representing the forward strand (+) and leftward arrows the reverse strand (−). Colored arrows represent orthologs across species, while white arrows with gray outlines indicate genes lacking a clear ortholog in the genomic region surveyed. Below the microsynteny plot, a schematic summarises the conserved syntenic neighbourhood of USOM3, where arrows with a dashed square indicate neighbour genes that may be absent in some species. USOM3 displays a conserved syntenic neighbourhood comprising two uncharacterised genes (NA) and *TRET1* (Facilitated trehalose transporter Tret1). (b) Phylogenetic tree of USOM3 gene models across coral species. Blue and purple squares correspond to Complexa and Robusta clades, respectively, which form distinct, separate branches consistent with a coral lineage-specific distribution of this gene.

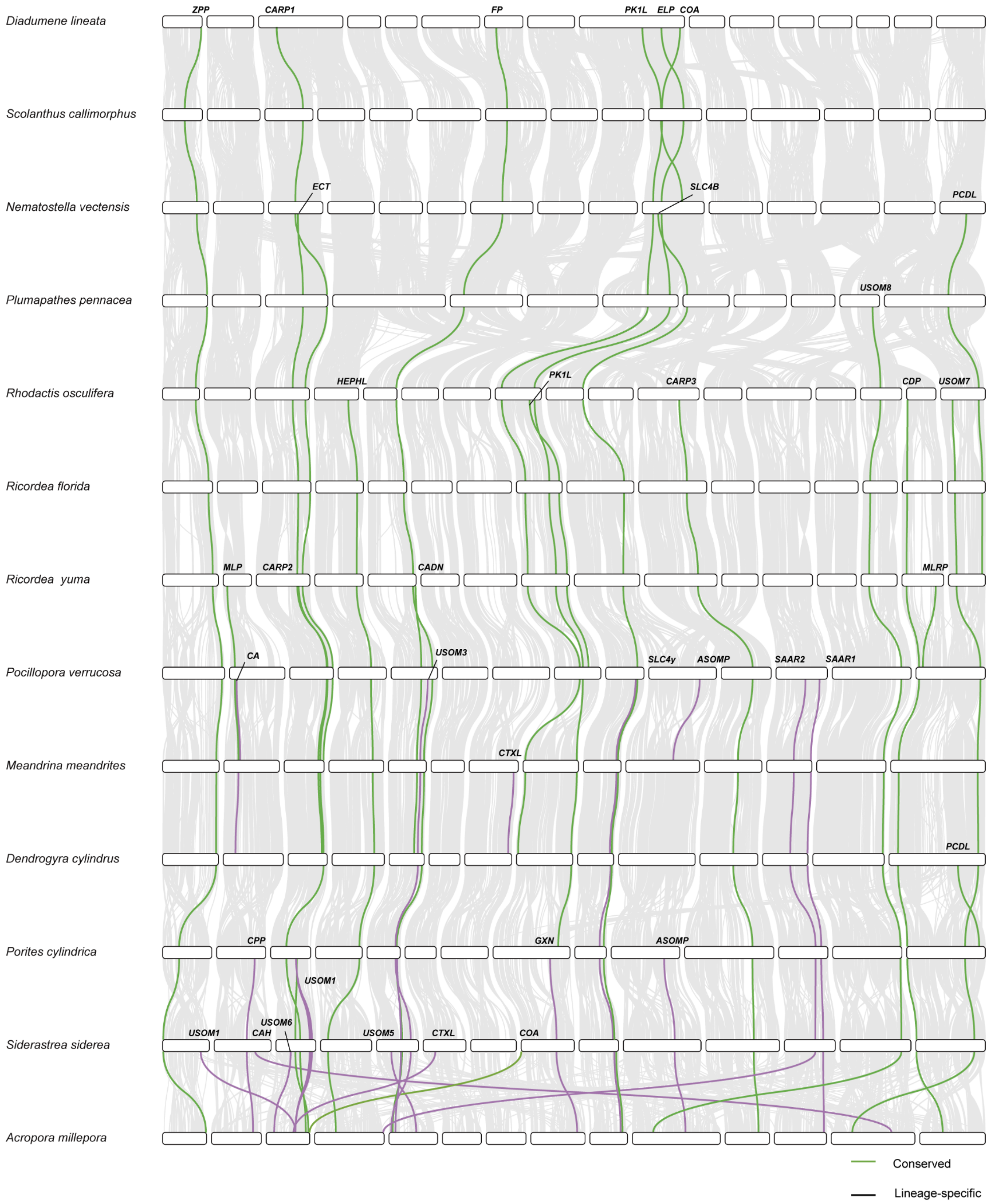


**Supplementary Figure 18.** SOMPs macrosynteny across hexacorallians. Vertical lines connect orthologous genes between species. Conserved and scleractinian-specific SOMPs are highlighted in green and purple, respectively, whereas other genes are shown in gray. Gene names are indicated adjacent to the corresponding lines.

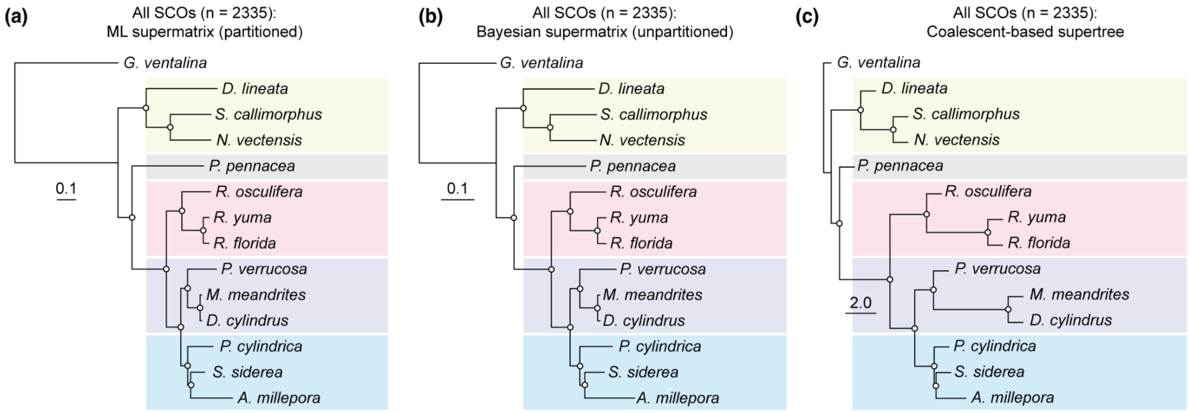


**Supplementary Figure 19.** Phylogenetic trees constructed using maximum likelihood supermatrix, Bayesian supermatrix and coalescent-based supertree methods all produce the same topology. (a) Maximum likelihood tree constructed using a partitioned analysis (-*p* option) using the phylogenetic terrace aware (PTA) data structure on a concatenated supermatrix of 2,335 single-copy proteins in IQ-TREE 3. Branch lengths represent substitution per site. Open circles represent 100% support from 1000 ultrafast bootstrap replicates. (b) Bayesian consensus tree constructed from the same dataset using MrBayes. Branch lengths represent substitutions per site. Open circles represent posterior probabilities of 1. (c) Coalescent-based supertree constructed using weighted ASTRAL. Branch lengths represent coalescent units. Open circles represent a local posterior probability of 1.

**Supplementary Table 1.** Repeat content of the *R. yuma* genome.

| **Class** | **Family** | **Count** | **Masked (bp)** | **Masked (%)** |
|---|---|---|---|---|
| LINE | CR1 | 4,956 | 1,653,192 | 0.29% |
| LINE | L1 | 4,905 | 2,675,233 | 0.46% |
| LINE | L2 | 30,747 | 12,245,979 | 2.12% |
| LINE | R2 | 66 | 80,952 | 0.01% |
| LINE | R4 | 2,763 | 1,268,805 | 0.22% |
| LINE | RTE | 2,419 | 1,369,039 | 0.24% |
| LINE | Rex | 3,686 | 1,896,675 | 0.33% |
| LINE | unknown | 12,420 | 2,354,355 | 0.41% |
| LTR | Bel_Pao | 2,902 | 965,492 | 0.17% |
| LTR | Copia | 1,590 | 436,567 | 0.08% |
| LTR | Gypsy | 110,840 | 65,555,611 | 11.36% |
| LTR | unknown | 204,485 | 38,954,363 | 6.75% |
| SINE | Alu | 56 | 18,423 | 0.00% |
| TIR | CACTA | 73,277 | 19,170,242 | 3.32% |
| TIR | Mutator | 102,319 | 39,393,335 | 6.82% |
| TIR | PIF_Harbinger | 59,365 | 15,599,506 | 2.70% |
| TIR | Tc1_Mariner | 9,906 | 2,364,800 | 0.41% |
| TIR | hAT | 9,948 | 2,242,933 | 0.39% |
| nonLTR | Penelope | 22,277 | 6,745,567 | 1.17% |
| nonTIR | helitron | 81,613 | 18,375,611 | 3.18% |
|  | repeat_fragment | 16,094 | 3,782,038 | 0.66% |
|  | TOTAL | 756,634 | 237,148,718 | 41.08% |

**Supplementary Table 2.** Number of orthologs used per pairwise comparison.

| **Species 1** | **Species 2** | **# orthologs used** | **Average # orthologs per ALG** |
|---|---|---|---|
| *Diadumene lineata* | *Scolanthus callimorphus* | 1,608 | 76.57142857 |
| *Scolanthus callimorphus* | *Nematostella vectensis* | 1,606 | 76.47619048 |
| *Nematostella vectensis* | *Plumapathes pennacea* | 2,358 | 112.2857143 |
| *Plumapathes pennacea* | *Rhodactis osculifera* | 2,330 | 110.952381 |
| *Rhodactis osculifera* | *Ricordea florida* | 2,361 | 112.4285714 |
| *Ricordea florida* | *Ricordea yuma* | 2,392 | 113.9047619 |
| *Ricordea yuma* | *Pocillopora verrucosa* | 2,317 | 110.3333333 |
| *Pocillopora verrucosa* | *Meandrina meandrites* | 2,252 | 107.2380952 |
| *Meandrina meandrites* | *Dendrogyra cylindrus* | 2,341 | 111.4761905 |
| *Dendrogyra cylindrus* | *Porites cylindrica* | 2,341 | 111.4761905 |
| *Porites cylindrica* | *Siderastrea siderea* | 2,299 | 109.4761905 |
| *Siderastrea siderea* | *Acropora millepora* | 1,728 | 82.28571429 |

**Supplementary Table 3.** Characterisation of the 8-gene location of ALG C1 in scleractinians. *Dcy*, *Dendrogyra cylindrus*; *Pcy*, *Porites cylindrica*; *Ryu*, *Ricordea yuma*.

| **OG** | **Dcy gene** | **Pcy gene** | **Ryu gene** | **Ryu gene position** | **Positioned on chr with ALG C1?** | **EggNOG-mapper description** | **Preferred name** | **PFAMs** |
|---|---|---|---|---|---|---|---|---|
| OG0003212 | g10944 | g14287 | g9269 | chr2:12724808-12737004 | Yes | Rho guanine nucleotide exchange factor | - | RhoGEF |
| OG0001604 | g10949 | g14292 | g9264 | chr2:12585795-12619511 | Yes | Belongs to the TRAFAC class myosin-kinesin ATPase superfamily. Myosin family | *MYO1E* | Myosin_TH1,Myosin_head,SH3_1,SH3_9 |
| OG0004920 | g10960 | g14302 | g9253 | chr2:12380290-12387893 | Yes | Cilia and flagella associated protein 53 | *CCDC11* | TPH |
| OG0003797 | g10978 | g14315 | g9575 | chr2:25376447-25377802 | Yes | Lysin motif | *LYSMD3* | LysM |
| OG0003790 | g10986 | g14321 | g9586 | chr2:25597834-25607375 | Yes | rRNA processing | *MPHOSPH10* | Mpp10 |
| OG0003788 | g10989 | g14325 | g9589 | chr2:25673206-25677834 | Yes | TeTratriCopeptide repeat | *TTC24* | TPR_10, TPR_12, TPR_16, TPR_2,TPR_4,TPR_7,TPR_8 |
| OG0003787 | g10990 | g14326 | g9590 | chr2:25678321-25686006 | Yes | cotranslational protein targeting to membrane | *SSR2* | TRAP_beta |

| | | | | | | | | |
|---|---|---|---|---|---|---|---|---|
| OG0004907 | g10993 | g14331 | g9594 | chr2:25813286-25844953 | Yes | SUMO transferase activity | *PIAS1* | PINIT,zf-MIZ |

**Supplementary Table 4.** RNA-Seq accession numbers used for BRAKER3 annotation.

| Order | Species | Genome Assembly Accession Number | RNA Accession Number(s) |
|---|---|---|---|
| Malacalcyonacea | *Gorgonia ventalina* | GCA_964194065.1 | SRR935083 |
| Actiniaria | *Diadumene lineata* | GCA_918843875.1 | ERR6688660; SRR8297741 |
| Actiniaria | *Scolanthus callimorphus* | GCA_033964015.1 | NA |
| Actiniaria | *Nematostella vectensis* | GCA_932526225.1 | SRR18908060; SRR18908065; SRR24348395; SRR24348403 |
| Antipatharia | *Plumapathes pennacea* | GCA_964036295.1 | SRR3407161 |
| Corallimorpharia | *Rhodactis osculifera* | GCA_965240485.1 | ERR13654243 |
| Corallimorpharia | *Ricordea florida* | GCA_949710005.1 | ERR10123701 |
| Scleractinia | *Pocillopora verrucosa* | GCA_036669915.2 | SRR27940236; SRR9648437 |
| Scleractinia | *Meandrina meandrites* | GCA_963693325.1 | ERR12321244 |
| Scleractinia | *Dendrogyra cylindrus* | GCA_964187815.1 | ERR13148289; SRR24899279 |
| Scleractinia | *Porites cylindrica* | GCA_964035525.1 | SRR14140469; SRR14140473 |
| Scleractinia | *Siderastrea siderea* | GCA_964030785.1 | SRR14295603; SRR14577702; SRR20761966 |
| Scleractinia | *Acropora millepora* | GCA_013753865.1 | SRR1853252; SRR3992866; SRR5374544 |

**Supplementary Table 5.** Fossil record data used in the MCMCTree for molecular dating.

| **Fossil** | **Age (Ma)** | **Reference** | **Node** | **Prior (Soft Bounds)** |
|---|---|---|---|---|
| *Bathycoenia statteri* | 168.2 - 165.3 | (Beauvais 1970) | 23 | B(0.66,1.65,1e-300,0.025) |
| *Maeandrina* sp. | 66.0 - 61.66 | (Wanner 1902) | 23 | B(0.66,1.65,1e-300,0.025) |
| *Eolympia pediculata* | 540 - NA | (Han et al. 2010) | 14 | B(5.4,7.0,1e-300,0.025) |
| *Oculina antiqua* | 266.9 - 264.2 | (De Gregorio et al. 1930) | 21 | B(2.6,4.2,1e-300,0.025) |

### Legends for Datasets

**Dataset S1 (separate file).** Metadata for genome assemblies used in this work.

**Dataset S2 (separate file).** Summary statistics of genome annotations.

**Dataset S3 (separate file).** Presence/absence of conserved gene blocks across hexacorallians. Values = number of copies.

**Dataset S4 (separate file).** Skeletal organic matrix protein orthologs across Hexacorallian species.

**Dataset S5 (separate file).** Annotation of genes differentially expressed during regeneration following fragmentation.

**Dataset S6 (separate file).** Categories of GO enrichment analysis.

### Supplementary References

Beauvais L. 1970. Sur quelques genres nouveaux ou peu connus de madreporaires jurassiques. Available from: https://doi.org/10.5169/seals-163878

De Gregorio A, De Gregorio A, De Gregorio A. 1930. Sul permiano Di Sicilia: (fossili Del calcare con fusulina Di Palazzo Adriano non descritti Dal prof. G. gemmellaro Conservati Nel Mio Privato Gabinetto). Annales de Géologie et de Paléontologie.

Han J, Kubota S, Uchida H-O, Stanley GD Jr, Yao X, Shu D, Li Y, Yasui K. 2010. Tiny sea anemone from the Lower Cambrian of China. *PLoS One* 5:e13276.

Levy S, Elek A, Grau-Bové X, Menéndez-Bravo S, Iglesias M, Tanay A, Mass T, Sebé-Pedrós A. 2021. A stony coral cell atlas illuminates the molecular and cellular basis of coral symbiosis, calcification, and immunity. *Cell* 184:2973–2987.e18.

Levy S, Grau-Bové X, Kim IV, Najle SR, Księżopolska E, Elek A, Montes-Espuña L, Montgomery SA, Mass T, Sebé-Pedrós A. 2025. The evolution of facultative symbiosis in stony corals. *Nature* 648:368–376.

Wanner J. 1902. Die Fauna der obersten weissen Kreide der lybischen Wüste. *Palaeontographica (1846-1933)*:91–152.
